# Effects of finfish farms on pelagic protist communities in a semi-closed stratified embayment

**DOI:** 10.1101/2022.08.08.503163

**Authors:** R.R.P. Da Silva, C.A. White, J.P. Bowman, D.J. Ross

## Abstract

Coastal aquaculture operations for feed additive species results in the release of waste into the surrounding environment, with the potential for adverse environmental change. Ubiquitous pelagic protists are sensitive to environmental changes making them potential sentinels for detecting and monitoring impacts. This study used 18S rRNA high-throughput amplicon sequencing as a molecular tool to study the pelagic protist community, with the aim of evaluating their potential as bioindicators of aquaculture activity in a low-oxygen, highly stratified marine embayment. Sampling occurred at three different depths along a distance gradient from two leases and at three control sites. Our results showed that the diversity and composition of both phytoplankton and other protist communities were more strongly influenced by depth stratification than the aquaculture activity. Nonetheless, differential abundance and machine learning analyses revealed a suite of potential bioindicators for aquaculture activity; this included the phytoplankton taxa *Chrysophyceae*, *Gymnodiniphycidae* (*Gyrodinium*), *Cryptomonadales* and *Ciliophora* (*Philasterides armatalis, Plagiopylida,* and *Strombidium).* Among the other protists, ciliates were also more abundant in closer proximity to the leases in both surface and bottom samples. Overall, our findings indicated that the use of 18S rRNA sequencing of protist communities is a promising tool for identifying environmental changes from aquaculture in the water column.

## Introduction

The aquaculture sector has been expanding globally over the last thirty years, raising concerns about the possible environmental effects (Stankus 2021). Carnivorous marine finfish aquaculture (e.g. Atlantic salmon - *Salmo salar*) is known to release significant volumes of waste as uneaten feed, faeces and metabolic products into the surrounding environment (Burridge et al. 2010; Navarro, Leakey & Black 2008; Zhang et al. 2015). These waste products contain high concentrations of carbon, nitrogen, and phosphorous which can lead to environmental changes such as eutrophication and oxygen depletion where they accumulate (Capone et al. 2008; Cloern 2001; Gardner et al. 2017). Previous studies have reported changes in the adjacent pelagic environment due to the release of such aquaculture waste (Alongi et al. 2003; McKinnon et al. 2010; Tičina, Katavić & Grubišić 2020; Tsagaraki et al. 2013; Welch et al. 2010), but knowledge around the biological significance of these changes remains limited. In semi-enclosed systems where water exchange rates are lower and nutrients are more likely to accumulate, the susceptibility to negative ecological impacts is likely to be greater (Husa et al. 2013; Jansen et al. 2018; Skogen et al. 2009). Understanding the response of the pelagic environments to additional nutrient loadings from aquaculture is critical to ensuring environmental sustainability, particularly in systems with limited water exchange (á Norði et al. 2011; Asami, Aida & Watanabe 2005; Belias et al. 2003; Elizondo-Patrone et al. 2015).

Protists are key components of aquatic ecosystems contributing to vital ecosystem services, such as nutrient and biogeochemical cycling (Azam et al. 1983; Azam & Malfatti 2007; Basu & Mackey 2018; Pierce & Turner 1992; Pomeroy et al. 2007). Protist communities are sensitive and can respond quickly to both natural and anthropogenic-driven changes in the environment, and as such can be used as sentinels of environmental perturbations (Nagasoe et al. 2010; Varkey et al. 2018; Zhang et al. 2018). The abundance and/or the diversity of specific taxa can be used to infer conditions and shifts of the aquatic environment (Das, Gauns & Naqvi 2019; Dorigo, Bérard & Humbert 2002; Needham & Fuhrman 2016). For instance, changes in the composition of phytoplankton and other protists such as ciliates have been reported in a variety of environments due to natural and anthropogenic changes such as water column stratification, salinity, light, temperature, nutrient concentration, pollution, eutrophication, and metal contamination (Anderson & Harvey 2020; Behnke et al. 2010; Edgcomb & Pachiadaki 2014; Kulaš et al. 2021; Kurobe et al. 2018; Leach et al. 2018; Oikonomou et al. 2015; Pawlowski et al. 2016; Weber et al. 2014). This includes aquaculture where the abundance of dinoflagellates and heterotrophic ciliates have been reported to increase in response to nutrient enrichment from fish farms (Buschmann et al. 2006; Forster et al. 2019; Olsen et al. 2014; Pitta et al. 2005).

Despite their importance, the diversity and ecology of most protistan taxa are still poorly understood. Moreover, environmental variation poses a challenge in efforts to untangle the effects of human induced change (e.g., nutrient enrichment) from natural variation (Jansen et al. 2016; McKinnon et al. 2010). Studies have used eukaryotic molecular biomarkers (i.e., 18S rRNA) along with more sophisticated data analysis techniques such as differential abundance and machine learning to enable a greater understanding of the effects of environmental changes and aquaculture on microbial ecology (Behnke et al. 2010; Dully et al. 2021; Edgcomb & Pachiadaki 2014; Laroche et al. 2021; Moncada et al. 2019; Verhoeven et al. 2018). Morever, high-throughput sequencing followed by bioinformatics pipeline is a faster and cost-effective alternative to traditional methods such as morphological identification (Apotheloz-Perret-Gentil et al. 2021; Cordier et al. 2018; Cordier et al. 2019). Therefore, the application of amplicon sequencing techniques provides the opportunity to enhance our understanding on the role of protist communities in the aquatic environment and their potential sentinels for environmental change.

Macquarie Harbour, located in Western Tasmania, is a highly stratified fjord-like system with restricted water exchange with the ocean. The water column is characterized by a brackish tannin-rich water layer on top of saltier waters. The major freshwater source carries a high amount of organic matter and flows into the southern end of the harbour. Mean monthly flows also vary according to the season (< 100 m^3^ sec^−1^ in summer and early autumn, and 500 m^3^ sec^−1^ late autumn, winter, and spring) (Carpenter et al. 1991; Hartstein et al. 2019; Teasdale et al. 2003). At the north end, marine waters enter through a narrow and shallow inlet allowing the renewal of the oxygen dissolved (DO) in the deeper waters. Marine intrusions in the harbour are likely driven by forces such as tide, wind, atmospheric pressure and freshwater inputs (Cresswell, Edwards & Barker 1989; Hartstein et al. 2019). The harbour has a history of anthropogenic activities and environmental stressors such as mining, damming and aquaculture (Crawford, Macleod & Mitchell 2003; King & Tyler 1982; Kirkpatrick, Kriwoken & Styger 2019; Koehnken 2005). Pelagic DO concentrations naturally drop with increasing depth reaching around 4 - 6 mg O_2_ l^−1^ at bottom layers. However, in recent years a considerable decline has been observed, with levels now 2 - 4 mg O_2_ L^−1^, and often lower at 25 m depth (Ross et al. 2021). This drawdown is thought to be related to a reduction in the frequency and scalability of DO recharge along with a high oxygen demand due to fish farm inputs. Atlantic salmon (*Salmo salar*) are the main species farmed in the Harbour, and in 2012, a major expansion of the industry in the harbour was approved (EPA 2012) with production peaking in 2015/16. In response to the declining oxygen levels and deterioration in environmental conditions (EPA 2017),the biomass cap for the Harbour has since been reduced to below pre-expansion levels. Previous studies have suggested that pelagic microbial oxygen demand is a key driver of the oxygen drawdown in the water column (Da Silva et al. 2021; Revill, Ross & Thompson 2016; Ross et al. 2021). For this reason, the aim of this study was to (i) investigate the composition and distribution of pelagic protists communities in the Harbour; (ii) the relationship with proximity to finfish farms; and (iii) to identify microorganisms that have the potential to be used as bioindicators of aquaculture activity.

## Material and methods

### Location and sample collection

In December of 2019, sampling was conducted at two fish farm leases (“Lease 1” and “Lease 2”) and three control sites (“Control”) in Macquarie Harbour (42°18’13.8”S; 145°22’22.7”E), Tasmania, Australia (Fig. 1). The two study leases were fully stocked at the time of sampling. At each of the leases, samples were collected at sites on replicate transects radiating from stocked cages using 10 L Niskin bottles (Tables 1 and S1). Samples were collected from the surface (0-4 m), middle (10 – 15 m) and bottom (20 – 37 m) of the water column at each site except for the site inside the cage. Each transect (Lease 1: n = 2; Lease 2: n=3) included 4 distances (inside the cage + 3 distances) at the surface, and 3 distances at middle and bottom layers. Sampling over distance (~ 0 – 85m; Table 1) was carried out by letting the boat drift along the direction of the current. Three control sites (>1500 m away from) were also sampled at the three depths mentioned above. Genetic material was collected on 0.22 µM polyethersulfone Sterivex^TM^ filter cartridges (Millipore, Darmstadt, Germany) using a multichannel peristaltic pump. Each sample was stored at −80°C until DNA extraction. The filtered water was used for dissolved inorganic nutrients and stored at −20°C until analyses. The Haversine formula was used to calculate the geographic distance between samples on the basis of latitude and longitude (Sinnott 1984).

**Fig. 1 –.**
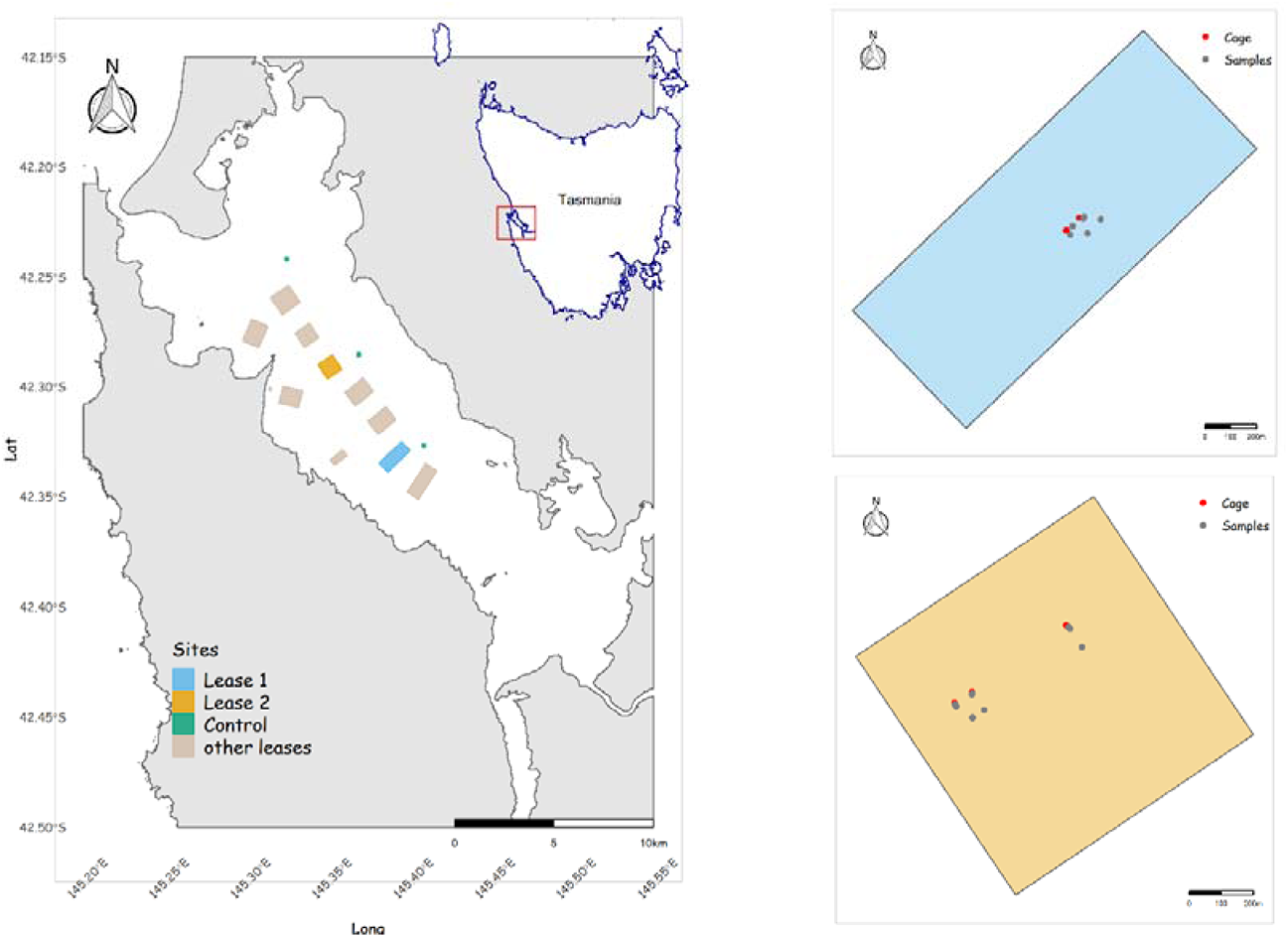
Map on the left displays the locations of sites sampled within this study. Green points are control sites and squares are leases located in the Macquarie Harbour. Sampled leases are the blue (Lease 1) and orange (Lease 2) ones. The two maps on the right show the sampled leases where the red points are the cages and the grey points are the samples collected along a distance gradient.

**Table 1 –.**
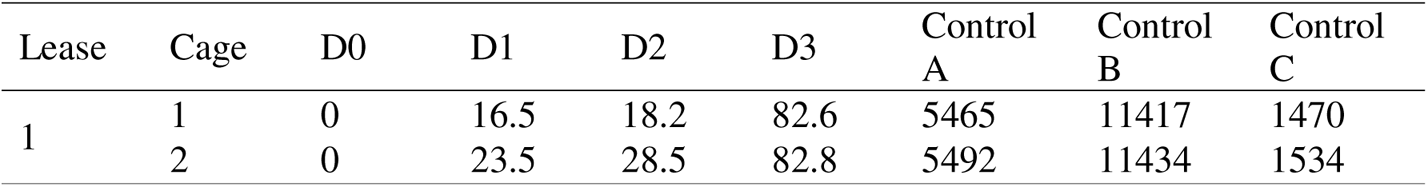
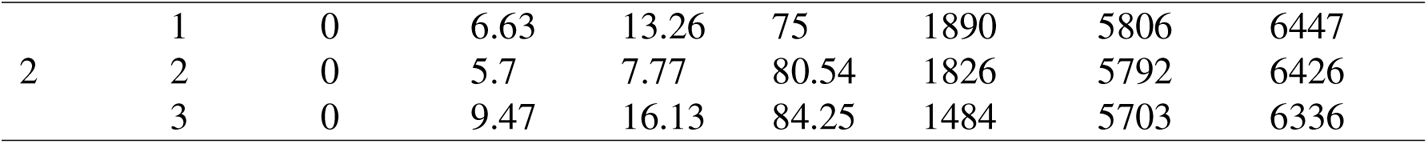
Distances (m) sampled from the cages in each lease calculated by the Haversine formula. D0 (0 m) is the water sample collected inside the cage.

### Environmental variables

Physiochemical parameters including temperature (°C), dissolved oxygen (DO, mg L^−1^), turbidity (NTU), salinity, and sample location (latitude & longitude) were measured *in situ* at each site using a YSI 6600 V2 Multi Parameter Water Quality Sonde with a YSI 650 MDS logger. Nutrient analyses were performed using a Bran + Luebbe AA3 HR segmented flow analyzer following standard spectrophotometric methods (Grasshoff, Kremling & Ehrhardt 2009). Detection limits for NO ^−^/NO_2_ ^−^ were 0.015 μmol L^−1^, for PO_4_ ^−3^ to 0.01 μmol L^−1^, and for NH_4_^+^ to 0.004 μmol L^−1^.

### DNA extraction and sequencing

DNA was extracted using a modified phenol:chloroform:isoamyl based DNA extraction protocol of the DNeasy PowerWater SterivexTM Kit (MoBio-Qiagen, vHilden, Germany) (Appleyard, Abell & Watson 2013) as described in Raes et al. (2018). Amplicon sequencing targeted the eukaryotic 18S rRNA V4 region (TAReuk454FWD1-TAReuk-Rev3) (Stoeck et al. 2010). Libraries were separately generated and sequenced for each sample at the Ramaciotti Centre for Genomics (University of New South Wales, Sydney) on the MiSeq sequencing platform (Illumina Inc., USA) using 300 bp paired-end sequencing. The raw sequences are publicly available at the Sequence Read Archive under the BioProject accession PRJNA803985 (https://www.ncbi.nlm.nih.gov/sra/PRJNA803985).

### Bioinformatic Pipeline

Paired end sequences of the 18S gene amplicons were analyzed through the DADA2 package pipeline, version 1.20.0 (Callahan et al. 2016) in R 4.1.0 environment (R Core Team 2021). Reads were filtered with the following parameters: truncLen = c(230,220), truncQ = 2, maxEE = c(2, 2). Merging of the forward and reverse reads was done with the *mergePairs* function using the default parameters (minOverlap = 50, maxMismatch = 0). Chimeras were removed using *removeBimeraDenovo* with default parameters. Taxonomic classification was performed using the *assignTaxonomy* function (default parameters) with the 18S SILVA v128 and v132 (Morien & Parfrey 2018) using the naive Bayesian classifier method described in Wang et al. (2007). ASVs abundance table, taxonomy and environmental data were imported into the phyloseq R package version 1.36.0 (McMurdie & Holmes 2013). After, data were separated into eukaryotic “Phytoplankton” and “Other Protists” groups. ASVs assigned to *Chlorophyta, Cryptophyta, Haptophyta, Ochrophyta, Cryptomonadales, Dinophyta, Chlorarachniophyta, Dinoflagellata,* and *Chrysophycea* were selected to be part of the “Phytoplankton” group. “Other Protists” groups encompassed all the other ASVs excluding the ones related to *Phaeophyceae* and *Xanthophyceae*, which are known to contain macroalgae as well as *Metazoa*, *Archaeplastida*, *Fungi*, and *Aphelidea’* that contain multicellular organisms. To reduce the data complexity (Cao et al. 2021), eukaryotic ASVs were filtered to keep taxa seen at least 2 times in at least 2 samples for further analyses, obtaining 409 out of 1688 ASVs in “Phytoplankton” group (2.76 % of total reads removed) and 772 out of 4351 ASVs in “Other Protists” group (2.28 % of total reads removed).

## Data Analysis and Statistics

### Alpha and beta diversity analysis

All data analysis and statistical tests were conducted in R 4.1.0 environment (R Core Team 2021). Alpha diversity indices (Shannon diversity index and Inverse Simpson) and Chao1 species richness were calculated using *estimate_richne*ss function from the phyloseq package. Both “Phytoplankton” and “Other Protists” groupings were centered log-ratio (CLR) transformed to address the negative correlation bias intrinsic to compositional data (Gloor et al. 2017) using CoDaSeq v.0.99.6 (Gloor 2016) and zCompositions v.1.3.4 (Palarea-Albaladejo & Martín-Fernández 2015) packages. To visualize differences in community structure among samples, Principal Composition Analysis (PCA) on Aitchison distance (Aitchison 1986) were performed using tidymodels v.0.1.3 (Kuhn & Wickham 2020). Environmental variables which could explain variation found on PCA were fitted using both the *envfit*, and *ordistep* functions, the latter with forward and backward stepwise selections. Autocorrelated environmental variables (i.e., r > 0.7, p-value ≤ 0.05) were combined into one for downstream analyses. To analyze the environmental and aquaculture effects on beta-diversity across all samples and at each depth separately, Mantel and partial Mantel (geographic distance and environmental variability controlled) tests were performed (Spearman rank correlation and 9999 permutations) using the vegan v.2.5.7 package (Oksanen et al. 2018).

### Feature selection and classification model

Pairwise quantitative analyses was performed using both ALDEx2 (Fernandes et al. 2013) and DESeq2 (Love, Huber & Anders 2014) packages to evaluate the differences in community composition between leases and control sites. As inputs, ASV table was clr-transformed for ALDEx2 whereas DESEq2 used variance-stabilized data. The combination of these two methods aims to overcome the drawbacks found in differential abundance methods, such as low power and high False Discovery Rate (FDR) (Hawinkel et al. 2019; Williams et al. 2017). ASVs with an effect size > |1.5| in ALDEx2 and an FDR-adjusted P value (FDR) < 0.05 and a log2foldchange > |2| in DESeq2 as well as with a relative abundance ≥ 60% were considered significantly associated to the examined site. Also, the random forest classification model was implement through the tidymodels v.0.1.3 (Kuhn & Wickham 2020) and randomForest v.4.6.14 (Liaw & Wiener 2002) packages to test the ability in classifying these sites using the pelagic eukaryotic 18S rRNA genes. ASVs were then used as predictors and sites (lease and control sites) as class labels. For comparison, these same ASVs were used as input to another random forest classification model to evaluate the ability in predicting depth as class label (surface, middle, and bottom water layers). A clr-transformed ASV table was first split into training and testing sets in a proportion of 3:4. Also, stratified sampling were performed only for the site classification model using “site” as the class label to maintain class ratios between train and test sets. Before training the site model, a Synthetic Minority Over-sampling Technique (SMOTE) step was applied to address the issue of imbalanced data (Chawla et al. 2002). The number of trees was set to 501 to all models. To tune the other hyperparameters (*mtry* and *min_n*), a k-fold cross validation scheme (k = 10 with 3 repeats and class label as strata argument) with 501 trees of the training set was used to estimate the best model based on the area under the ROC curve (ROC-AUC). The ROC-AUC metric informs how much the model is able to discriminate between classes. The model was finalized using the *finalize_workflow* and *select_best* functions. Model performance was tested in the training set and the metrics ROC-AUC, accuracy, sensitivity, and specificity were collected using the *collect_metrics* function. Variable importance was measured by Mean Decrease in Gini Index. The variable importance describes which features are relevant to describe a given class label. We considered potential aquaculture activity bioindicators the ASVs that were detected by at least two of these three methods (Shade & Handelsman 2012). All visualizations were created using vip v.0.3.2 (Greenwell & Boehmke 2020), tidyverse v.1.3.1 (Wickham et al. 2019) and ggtern v.3.3.5 (Hamilton & Ferry 2018) packages. Code and data used in this study can be found in the associated GitHub repository (https://github.com/ricrocha82/MH_2019_18s).

## Results

### Environmental variables

The water column of Macquarie Harbour showed a clear stratification of all environmental variables (Table 2; Fig. S1). NH_4_ and NO_2_ decreased with depth whereas NOx and PO_4_ displayed the opposite trend. Similarly, fluorescence, oxygen, PAR, and turbidity decreased with depth, while salinity and temperature all increased. There was no clear relationship observed between environmental parameters and distance from the cage, indicating that the natural stratification is strong enough to prevail over any sign of variation in physicochemical variables caused by aquaculture activity.

**Table 2 -.**
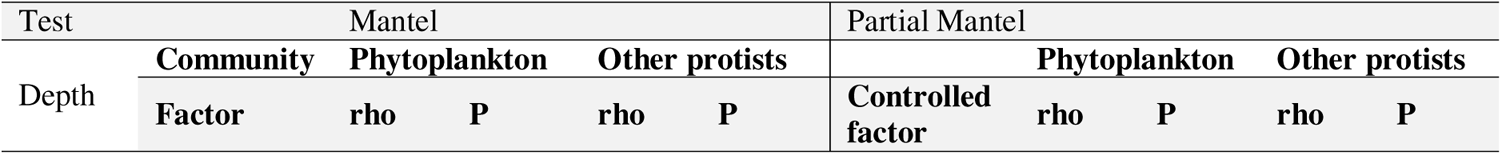
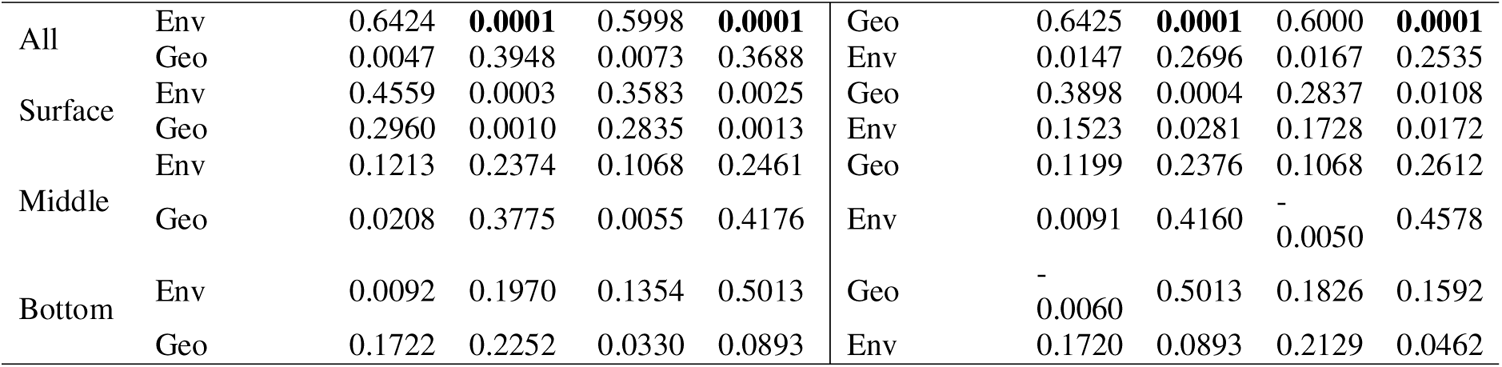
Mantel and Partial Mantel tests showing the correlation between beta-diversity (Euclidean distance matrices), geographic distance (Geo), and cumulative environmental condition (Env) considering all samples together and samples separately by depth. Significant strong correlations with |ρ| ≥ 0.5 and p-value (P) ≤ 0.01 are highlighted with bold type.

### Protist diversity and community composition

Alpha diversity indices and richness increased with depth. A trend can be observed at Lease 1 where diversity and richness increased from the cage to control site for both phytoplankton and other protists communities (Fig. 2; Table S2).

**Fig. 2 –.**
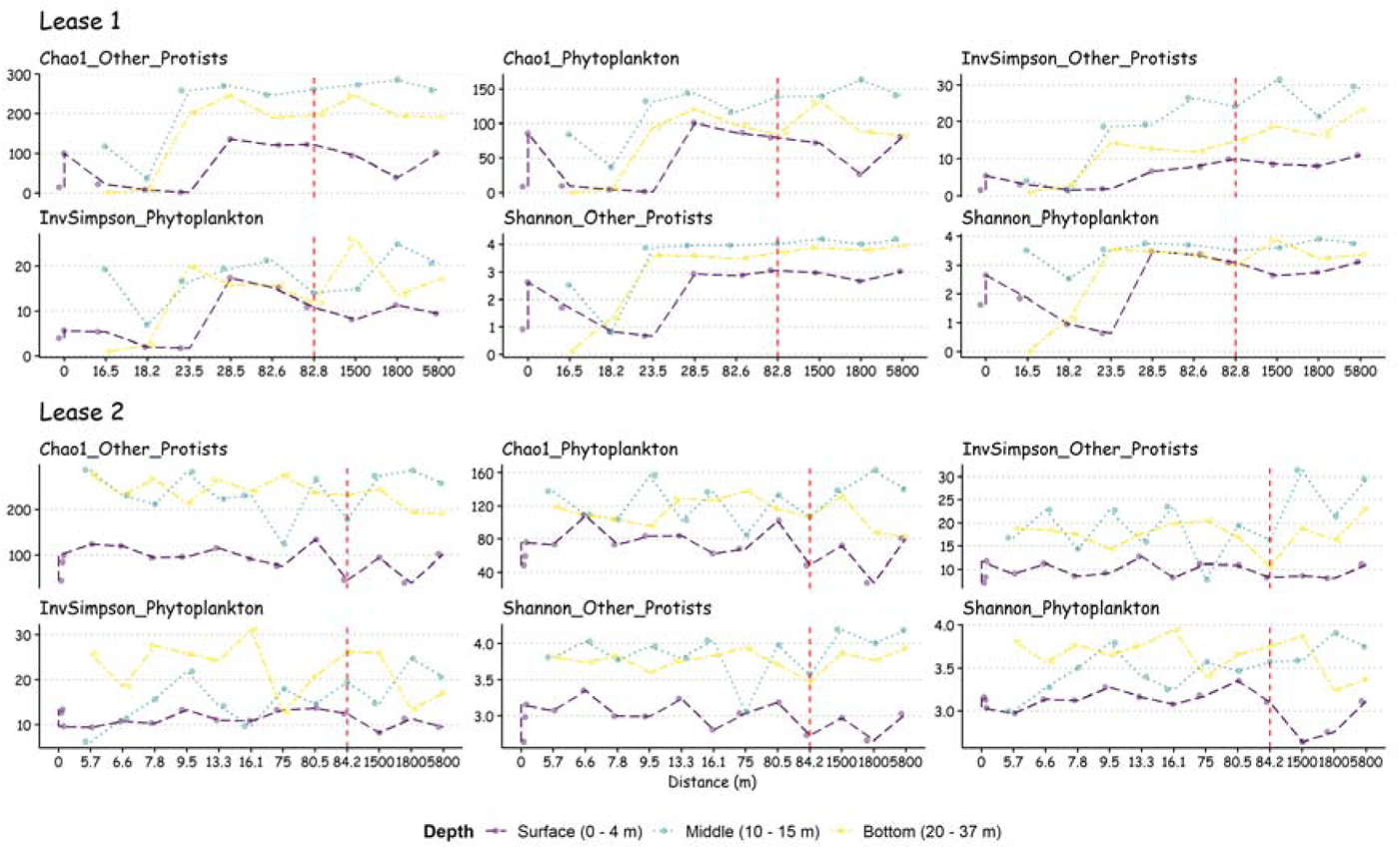
Alpha diversity indices along depth and distance gradients in both lease and control sites. The left-hand side of the dashed red line indicates the lease area whereas the control sites (distance > 1500 m from the fish cages) are at the right-hand side of the line. Distance = 0 m indicates surface water sample collected inside the fish cage.

Depth stratification of both communities was evident from PCA analysis and both simple and partial Mantel tests (Fig. 3; Tables 2 and 3 Table). Total variation explained by the first two principal components was 36.45% and 39.81% for phytoplankton and other protists communities, respectively. Beta diversity of both communities were significantly correlated (p < 0.01) to NOx and highly correlated peers (phytoplankton: r^2^ = 0.75; other protists: r^2^ = 0.69), NH_4_ and NO_2_ concentrations (phytoplankton: r^2^ = 0.49; other protists: r^2^ = 0.53) and turbidity (phytoplankton: r^2^ = 0.62; other protists: r^2^ = 0.46). When all samples were considered in the simple Mantel test, a significant correlation (rho = 0.64; P = 0.001, Table 2) was found in both communities indicating that beta diversity was significantly affected by the cumulative environment conditions even when geographic distance was controlled by the partial Mantel test. When depth was removed from the analysis by running both tests with samples from each water layer only, correlations became weaker and/or not significant. oth tests and ordination analysis were unable to detect variation in the community along a distance gradient from the cages. The results indicate that changes in protist beta diversity in Macquarie Harbour is more influenced by depth stratification than aquaculture activity.

**Fig. 3 –.**
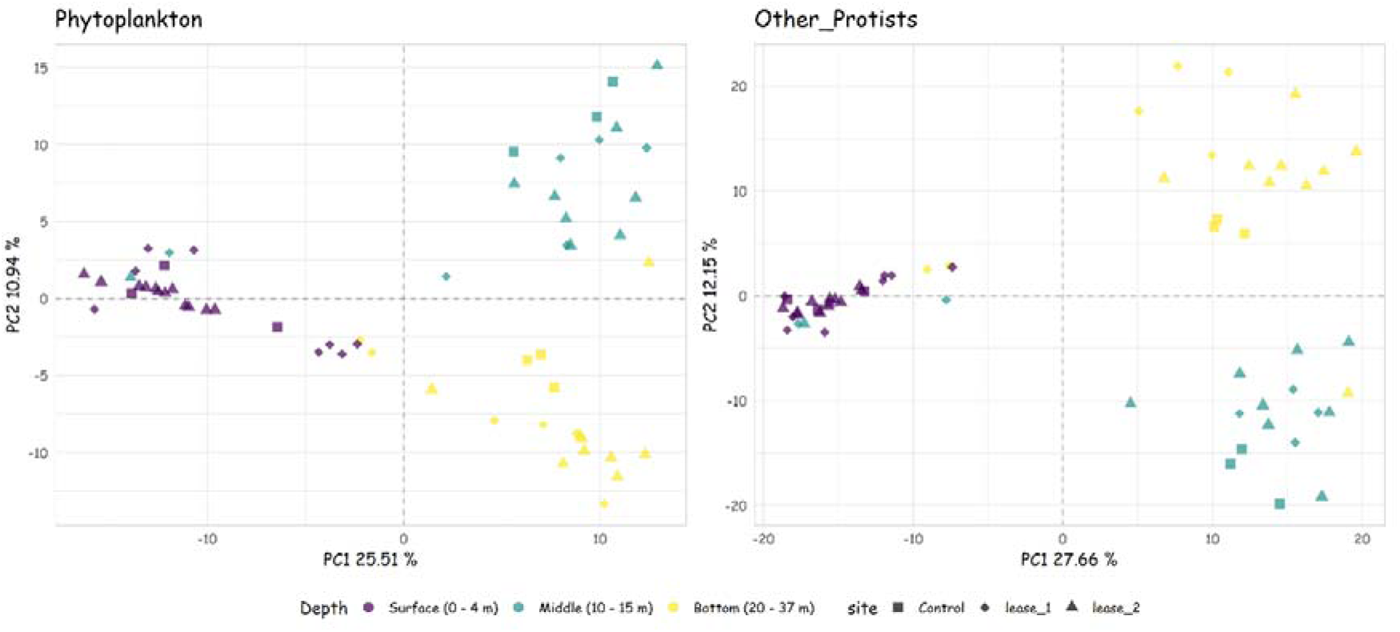
PCA plots showing eukaryotic community structure among samples from control and lease sites.

**Table 3 –.** Correlation between phytoplankton and other protists community beta diversity and environmental factors (*ordistep* and *envif* functions) in the Macquarie Harbour. We considered significant if the environmental variable had a p-value < 0.01 in both analyses. To avoid multicollinearity, NOx included conductivity, salinity, temperature and PO_4_. NH_4_ included NO_2_.

| Community | Factor | p-value<br>(ordistep) | p-value<br>(env.fit) | R2<br>(envfit) |
| --- | --- | --- | --- | --- |
| Phytoplankton | NOx | 0.005 | 0.001 | 0.75 |
|  | turbidity | 0.005 | 0.001 | 0.62 |
|  | NH<sub>4</sub> | 0.005 | 0.001 | 0.49 |
| Other protists | NOx | 0.005 | 0.001 | 0.69 |
|  | turbidity | 0.005 | 0.001 | 0.46 |
|  | NH<sub>4</sub> | 0.005 | 0.001 | 0.53 |

\*NOx included conductivity, salinity, temperature and PO<sub>4</sub>.
\*\*NH<sub>4</sub> included NO<sub>2</sub>.

At all sites, the phytoplankton dataset was dominated by *Dinoflagellata* and *Ochrophyta*, comprising more than 85 % mean relative abundance at Lease 1, 78.71 % at Lease 2 and 85 % at the control sites. Among the other protists, *Protalveolata* and *Ciliophora* were the most abundant groups at all sites, representing 79.93 %, 71.15 % and 69.51 % of total mean relative abundance at Lease 1, Lease 2 and control sites, respectively (Figs. S2 and S3).

## Feature Selection

### Differential Abundance Analyses

To detect differences in the eukaryotic community between the lease and control sites we used three approaches commonly used in microbial community analysis (ALDEx2, DESEq2 and random forest classification model). Two differential abundance analyses (ALDEx2 and DESeq2) were then used to examine the depth stratification effect previously observed in the multivariate analyses. Pairwise comparisons were performed grouping the ASVs by depth layer (i.e., surface, middle and bottom).

Among phytoplankton ASVs, a total of 13 and 21 ASVs were detected by ALDEx2 and DESeq2, respectively (Table S3). At Lease 1, an ASV related to *Lauderia annulata* was more abundant at the bottom whereas two *Dinoflagellata* ASVs (uncultured *Dinophyceae* and *Gyrodinium*) were enriched at the surface. A greater number of ASVs were found at Lease 2 (Table S3) where three ASVs assigned to *Chrysophyceae* and one ASV from uncultured *Ochrophyta* where more differentially abundant at the bottom. One ASV belonging to *Cryptomonadales* and two to *Crustomastix* (*Dolichomastigales*) and *Chrysophyceae* were more abundant at the middle and surface layers, respectively. Only one ASV, assigned to an uncultured *Dinoflagellata*, was more abundant (i.e., relative abundance ≥ 60%) in the bottom samples at control sites.

For the “other protists”, 28 and 46 ASVs were considered differentially abundant by ALDEx2 and DESeq2, respectively. At the bottom and middle layers of the control sites, the five most abundant ASVs were assigned to the *Syndiniales* group (Table S3). Seven ASVs related to *Ciolophora* and *Cercozoa* were found enriched at the surface and bottom waters at Lease 1. Following the same pattern as the phytoplankton group, a greater number of more abundant ASVs were detected at Lease 2 (n = 14). Again, ASVs assigned to *Ciliophora* and *Cercozoa* were more abundant at the bottom and surface. Groups that were enriched at the bottom included *Syndiniales*, *Codonosigidae* (*Choanoflagellida*), *Rhizaria* (*Retaria*) and *Stramenopiles* (MAST-2).

### Random forest Classification Models

Random forest classification models were used to identify important ASVs that could be used to classify samples according to site (i.e., Lease 1; Lease 2 or control) and depth (i.e., surface, middle and bottom). The top 10 most important phytoplankton ASVs to classify lease and control sites belonged to *Cryptomonadales*, *Dinoflagellata*, *Ochrophyta*, *Chlorophyta* and *Prymnesiales* (Fig. S4) while the top 10 most important from the other protists group were assigned to *Cercozoa*, *Syndiniales*, uncultured *Alveolata*, uncultured *Centrohelida*, *Euamoebida* (*Amoebozoa*), *Thraustochytriaceae* (*Stramenopiles*) and *Pirsonia* (*Stramenopiles*) (Fig. S5). Among the most important phytoplankton ASVs that classified the samples according to depth were related to Dinoflagellates and *Chrysophyceae*. Among the other protists, the important ASVs were assigned to *Protalveolata*, *Ciliophora*, *Bicosoecida*, *Syndiniales*, and *Leucocryptos* (Table S5).

Overall, depth classification models using both phytoplankton and other protists ASVs outperformed the site classification models (Table 4). Moreover, the top 20 most important ASVs in classifying depth displayed a clearer pattern relative to those that characterised lease and control sites (Figs. 4, 5, S6 and S7). This is evident from the ternary plots; most of the ASVs that classify depth are found at the edges of the plot denoting a clear separation between the communities of the surface and the deeper layers. On the other hand, the most important ASVs used to classify lease and control sites are more spread across the ternary plots suggesting more homogeneous distribution among these sites. The top 20 most important ASVs defined by the random forest classification models can be found in Table S4 (site) and Table S5 (depth).

**Fig. 4 –.**
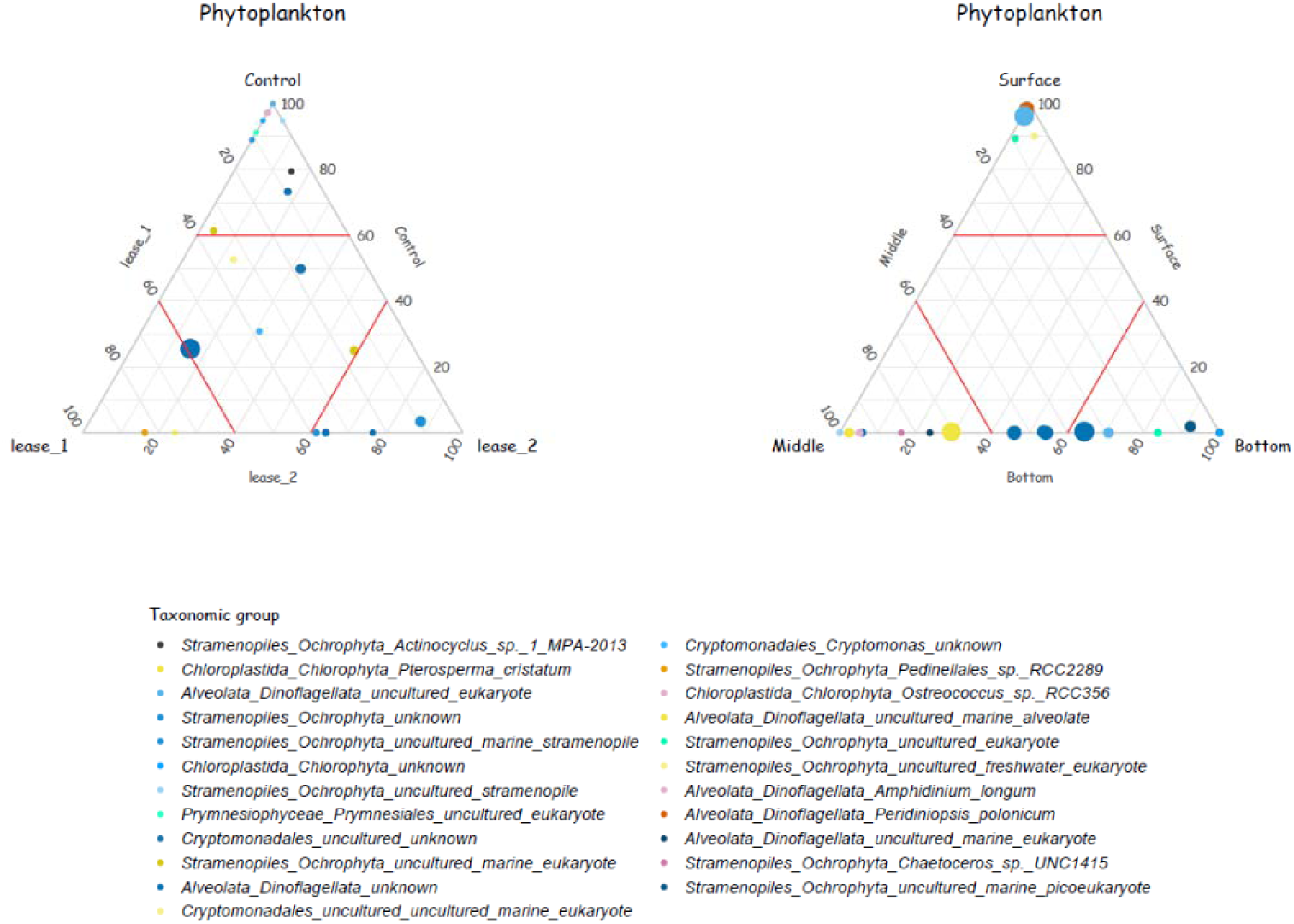
Ternary plots showing the distribution of the top 20 most important phytoplanktonic ASVs to classify sites (lease and control sites) and depth (surface, middle, and bottom water layers) according to random forest classification model. Each point represents an ASV, and its position indicates the proportion of its relative abundance at each site. Points closer to the ternary plot corners indicate that a greater proportion of the total relative abundance of this ASV was found in this site. Point colors indicate the ASV taxonomic group. The size of each point indicates the mean relative abundance of the ASV in all samples. The red lines inside the ternary plot indicate the 60% level of relative abundance of each of the sites.

**Fig. 5 –.**
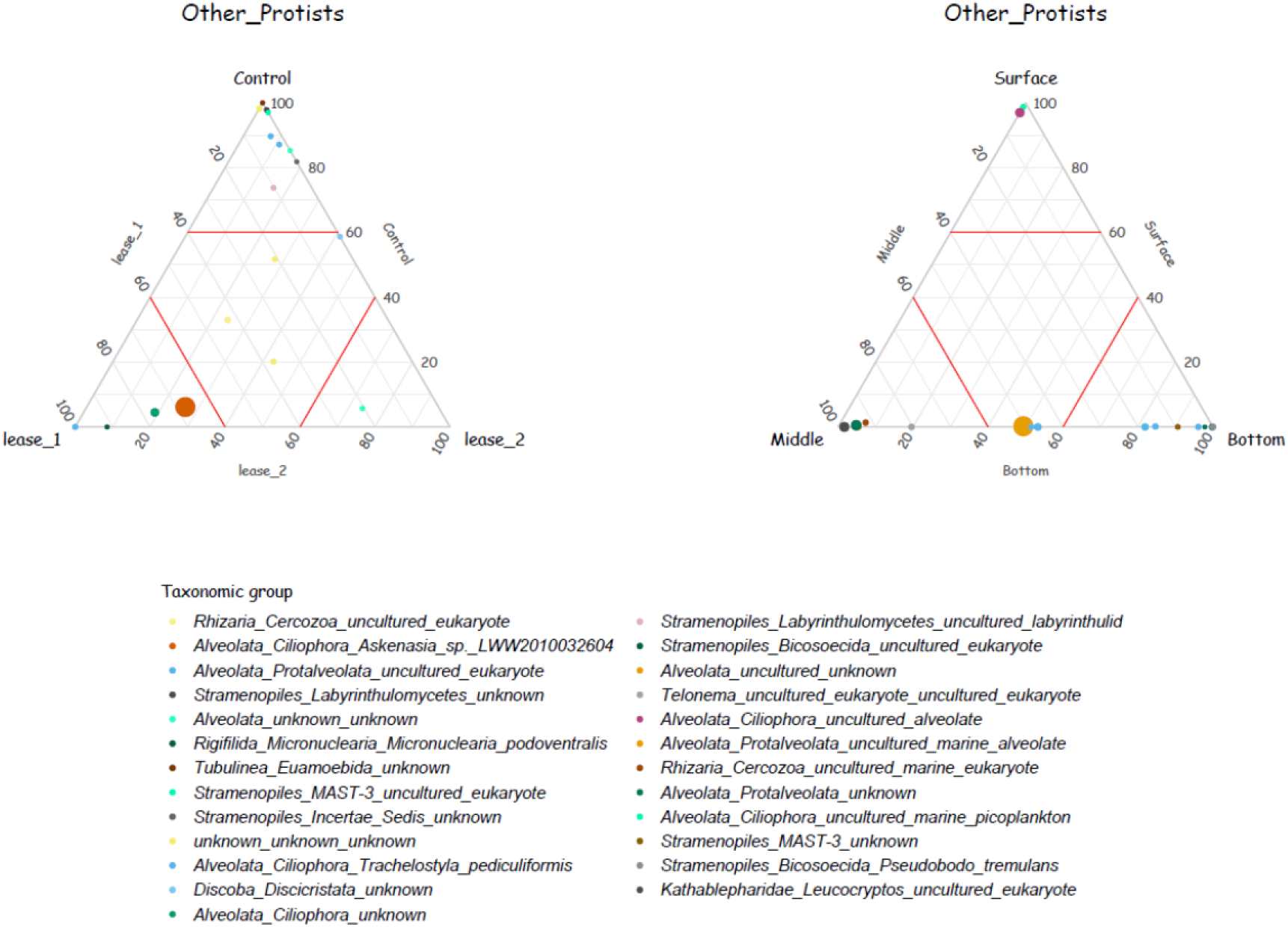
Ternary plots showing the distribution of the top 20 most important protists ASVs (other protists) to classify sites (lease and control sites) and depth (surface, middle, and bottom water layers) according to random forest classification model. Each point represents an ASV, and its position indicates the proportion of its relative abundance at each site. Points closer to the ternary plot corners indicate that a greater proportion of the total relative abundance of this ASV was found in this site. Point colors indicate the ASV taxonomic group. The size of each point indicates the mean relative abundance of the ASV in all samples. The red lines inside the ternary plot indicate the 60% level of relative abundance of each of the sites.

**Table 4 –.** Model classifier performance indices and tunned model hyperparameters of each microbial eukaryotic group to classify site (lease and control sites) and depth (surface, middle, and bottom water layers). Accuracy measures how many classes were correctly predicted by the model. In other words, it measures the proportion of true positives results in the selected data. Sensitivity (True Positive Rate) is the ability of the model to recognize a true positive that actually belongs to the true positive group. For instance, it measures the probability of classifying a sample from a site among those that belong to the same site. On the other hand, specificity (False Negative Rate) measures the classification capacity without giving false-positive results. For instance, it gives us the chance that a sample was correctly classified as not belonging to a particular site. mtry = number of features randomly selected as candidates for each split. min_n = minimal node size to control the depth and complexity of the individual trees. The seed used to run the models was 4848.

| **Classification class** | **Sites** |  | **Depth** |  |
| --- | --- | --- | --- | --- |
| **Group** | **Phytoplankton** | **Other Protists** | **Phytoplankton** | **Other Protists** |
| **Metrics** |  |  |  |  |
| **accuracy** | 0.750 | 0.688 | 0.938 | 0.875 |
| **sensitivity** | 0.625 | 0.583 | 0.933 | 0.867 |
| **specificity** | 0.879 | 0.837 | 0.967 | 0.936 |
| **ROC - AUC** | 0.785 | 0.751 | 1 | 1 |
| **Hyperparameters** |  |  |  |  |
| **number of trees** | 501 | 501 | 501 | 501 |
| **mtry** | 47 | 27 | 42 | 237 |
| **min_n** | 8 | 4 | 9 | 10 |

### Potential Pelagic Protists Bioindicators for Aquaculture Activity

To identify potential bioindicators, we assumed that common ASVs found by at least two of the three approaches could be an important indicator of aquaculture activity. The three methods showed divergent results, with ALDEx2 and DESeq2 having more shared ASVs than the random forest model (Fig. S8 and Table S6). For example, when the total of important ASVs determined by the random forest models was increased from 10 to 40 there was only a small increase in shared ASVs found by at least two of the three methods (from 5 to 7 phytoplankton ASVs and from 9 to 12 other protists ASVs). The distribution along the distance gradient of these common ASVs for each depth are displayed in Fig. 6 (phytoplankton) and Fig. 7 (other protists). Despite the variation in their distribution, some ASVs displayed interesting patterns (Figs. S9 and S10). In the phytoplankton community, ASVs 127 (*Phaeophyceae*) and 175 (*Chrysophyceae*) were more abundant at the surface of Lease 2 and decreased with distance from the cages. At Lease 1, ASVs assigned to *Chrysophyceae* (147, 181 and 97) increased in abundance with increasing distance from the cage while ASVs 268 (*Cryptomonadales*) and 78 (*Gyrodinium*) were more abundant in the middle and surface layers at the lease relative to control sites, respectively.

**Fig. 6 -.**
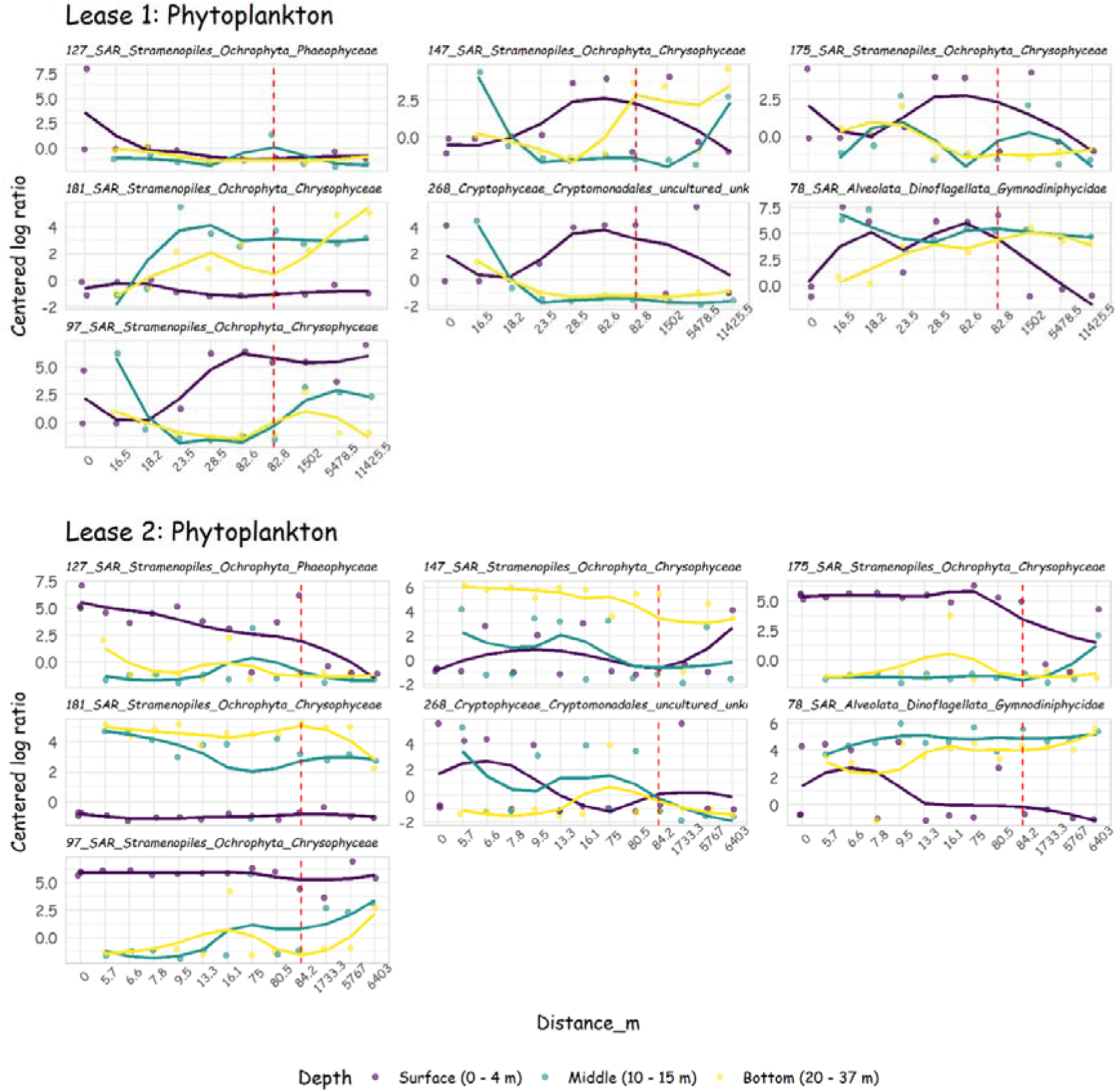
Distribution of phytoplanktonic ASVs commonly found in at least 2 out of the three methods (ALDEx2, DESEq2 and random forest classification model) along distance gradient. Each line represents the distribution of the ASVs in each depth. 0 m indicates the surface water sample collected inside the fish cage. Centered log ratio = log10(ASV read count / geometric mean). The SILVA nomenclature of each ASV is displayed above the plots.

**Fig. 7 -.**
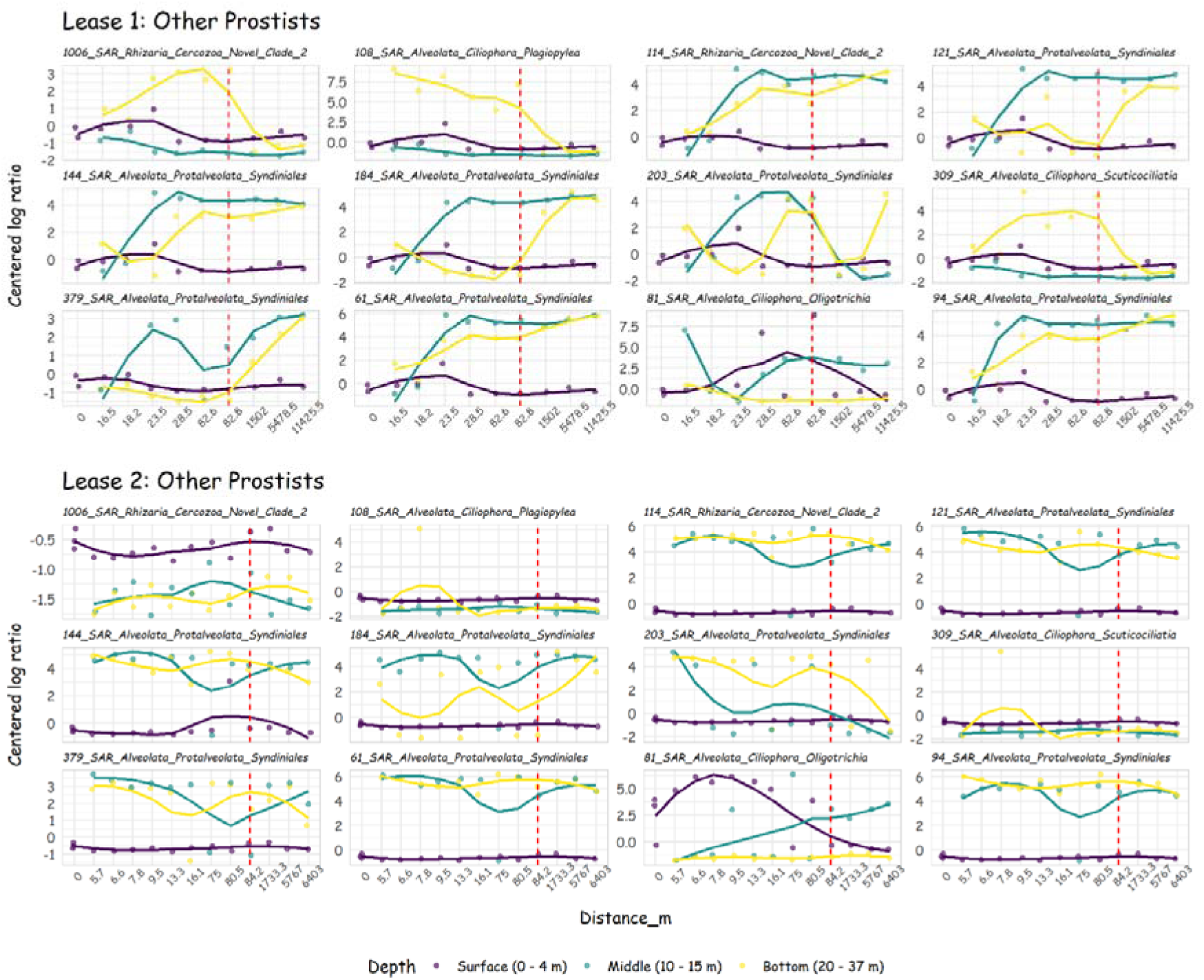
Distribution of protists ASVs (other protists) commonly found in at least 2 out of the three methods (ALDEx2, DESEq2 and random forest classification model) along distance gradient. Each line represents the distribution of the ASVs in each depth. 0 m indicates the surface water sample collected inside the fish cage. Centered log ratio = log10(ASV read count / geometric mean). The SILVA nomenclature of each ASV is displayed above the plots.

Among the other protist group, most ASVs related to *Syndiniales* were absent or were in relatively low abundance close to the cages at Lease 1. At Lease 2, *Syndiniales* ASVs were in greater abundance in the middle and bottom water layers but there was no clear pattern with proximity to the cage. A similar pattern was found for the cercozoan ASV 114; it was more abundant in the middle and bottom layers at both leases, but the increase in abundance further from the cages was evident for Lease 1 but not Lease 2. Another cercozoan ASV (1006) was comparatively more abundant in the bottom water samples of Lease 1 relative to the controls but was otherwise absent or in low abundance at the other depths and at Lease 2. Three *Ciolophora* related ASVs (ASVs 81, 108 and 309) were not detected at the control sites. ASVs 108 and 309 were both more abundant below the surface at Lease 1 while ASV 81 decreased with distance from the cage at the surface of Lease 2. From the results, the ASVs related to *Phaeophyceae* (*Ectocarpus* sp.), uncultured *Cryptomonadales, Gymnodiniphycidae* (*Gyrodinium*), and *Ciliophora* (*Philasterides armatalis, Plagiopylida,* and *Strombidiumi*) groups were not detected at control sites suggesting that they may be potential pelagic bioindicator candidates of aquaculture activity.

## Discussion

18S rRNA amplicon sequencing was used to describe the phytoplankton and other protist communities within the water column of Macquarie Harbour, a semi-enclosed coastal embayment in Tasmania, Australia. A site of significant finfish aquaculture activity, the study also assessed the distribution of pelagic communities in proximity to active leases compared to control sites. Through multivariate and supervised analyses, it was evident that depth stratification had a greater influence on phytoplankton and other protistan communities than did proximity to aquaculture activity. Using three techniques commonly used in targeted next generation sequencing (ALDEx2, DESEq2 and random forest classification model), several ASVs were identified as potential bioindicators for aquaculture activity. Given the “one-time” sampling strategy performed in this study and the natural heterogeneity of the Macquarie Harbour, the capacity to draw conclusions regarding the effects of the salmon aquaculture on phytoplankton communities in the Harbour is limited. Instead, the study offers an important insight into the environmental drivers of the pelagic protist community and a first glimpse of the potential effects of the aquaculture.

### Depth stratification and the influence on pelagic eukaryotic community structure

Depth was the major driver of both environmental variation and the diversity and composition of both phytoplankton and protist communities. Also, a vertical variation in the environmental variables was observed. Macquarie Harbor is a highly stratified system with the surface layer heavily influenced by inputs from the main freshwater source in the southern end of the harbour. The saltier layers below the surface only slowly mix with the riverine water, resulting in a distinct meromixis-like halocline stratification (Hartstein et al. 2019; Terry 2001). Alpha and beta diversity indices were affected by a vertical gradient in the water column suggesting the deep waters are inhabited by a eukaryotic community that is different to the upper; this is consistent with descriptions of the prokaryotic community in the Harbour (Da Silva et al. 2021). Depth-associated changes in environmental factors occur in highly stratified environments such as meromictic lakes and coastal embayments (Andrei et al. 2015; Cabello-Yeves et al. 2021; Oikonomou et al. 2015). Also, stratification reduces microbial dispersion among water layers separating microbial communities according to physiochemical conditions. (Costa, Santana & Neumann-Leitao 2018; DeLong et al. 2006; Mena et al. 2019). This changes in the water column influence both alpha and beta diversities. This result corroborates other studies showing that turbidity, oxygen, salinity, and nutrient concentrations are factors affecting protist community structure and distribution in coastal regions (Das, Gauns & Naqvi 2019; Jacquemot et al. 2021; Lallias et al. 2015; Parris et al. 2014; Sun et al. 2019). In the case of Macquarie Harbour, the higher turbidity, suspended particulate matter and NH_4_ concentrations at the surface could be related to the tannin-rich waters from Gordon River affecting light penetration, thereby favoring a heterotrophic community (Bartl et al. 2018; Da Silva et al. 2021; Dokulil 1994; Hartstein et al. 2019; Terry 2001).

At one of the study leases (Lease 1), an increase in richness and diversity of both communities with increasing distance from the cages was observed. Other studies on the impacts of aquaculture on aquatic microbiota (both eukaryotic and bacterial) have reported a similar trend. For instance, Jiang et al. (2012) studying three different aquaculture types in a semi-enclosed bay found significant differences in phytoplankton community diversity only in finfish farms when compared to oyster and kelp aquaculture. In the sediments, bacterial and eukaryotic benthic diversity increased from cage toward reference sites. High accumulation of organic matter due to faeces and feed input, environmental heterogeneity, and low water exchange were considered the major factors affecting these communities (Dully et al. 2021; He et al. 2019; Jiang et al. 2012). In this study, Lease 2 had a lower stocking density compared with Lease 1, which was at the maximum capacity (data not shown). As such, outputs of nutrient and organic material were likely to be higher at Lease 1, which may drive some of the differences observed between leases. Additionally, Lease 1 is situated further south in the Harbour where deep water exchange is less frequent and prominent, possibly leading to a less pronounced dispersion of nutrients (Hartstein et al. 2019; Ross et al. 2021). To further validate the role of aquaculture activity against natural environmental variation, repeat sampling across different stages of the production cycle is required.

### Vertical Distribution of Important Unicellular Eukaryotes

The multivariate and machine learning analyses also highlighted a much stronger influence of depth stratification than proximity to aquaculture activity on eukaryotes in Macquarie Harbour. The random forest classification models obtained a more accurate and precise classification of each water layer (i.e., surface, middle and bottom waters) in relation to the sites (i.e., Lease 1, Lease 2 and control).

Among the 10 most important phytoplankton ASVs in classifying depth, nine are associated with dinoflagellates. The most abundant dinoflagellates ASVs at the surface are *Thoracosphaeraceae* (*Peridiniopsis polonicum* and *Pfiesteria* sp.). Two of the species *Pfiesteria* sp. and *Peridiniopsis polonicum* are potential toxin producers (Burkholder, Glasgow & Deamer-Mella 2001; Burkholder et al. 2005; Hashimoto et al. 1968; Roset et al. 2002). Below the surface, ASVs assigned to an uncultured *Dinophyceae* and one to the genus *Heterocapsa* were enriched in the middle and bottom layer, respectively. *Dinophyceae* is a group commonly found in freshwater and estuarine habitats (Glasgow et al. 2001; Hashimoto et al. 1968; Iwataki 2008; Johnsen et al. 1994; Legrand, Graneli & Carlsson 1998). Dinoflagellates have been linked to high levels of ammonia and dissolved organic nitrogen in estuarine environments and aquaculture ponds, influenced by riverine and anthropogenic inputs (e.g., nutrient enrichment, light attenuation, humic content) (Dagenais-Bellefeuille & Morse 2013; Glibert & Terlizzi 1999; Gómez 2012; Prabhudessai, Vishal & Rivonker 2019; Prakash & Rashid 1968). As such, the dominant riverine influence in Macquarie Harbour with its high tannins, low light and humic content is naturally conducive for dinoflagellates, and the additional nutrient and organic matter inputs from aquaculture may be further promoting their abundance. Repeat sampling at different stage of production is required to better determine the role of aquaculture.

The ASVs characteristic of each depth strata provide insight into their functional role in the pelagic food web (Sherr & Sherr 2002; Steele et al. 2011; Worden et al. 2015). The dinoflagellates *Amphidinium longum* and katablepharid *Leucocryptos* are planktonic predators, and their predatory capacity is thought to be influenced by both light, by UV exposition, and prey availability (Havskum & Riemann 1996; Mikaelyan, Pautova & Fedorov 2021; Strom et al. 2003; Strom et al. 2020; Vørs 1992). This may help explain why they were predominately found at depth in this study. Important parasitic ASVs were enriched across the water column. The heterotrophic genus *Chytriodinium* were enriched both at the surface and in bottom waters while *Syndiniales* (Group II and *Amoebophrya*) were more dominant at depth. The distribution of *Chytriodinium* and *Syndiniales* rely on host and nutrient availability (Anderson & Harvey 2020; Fulton 1984; Gómez & Skovgaard 2015; Strassert et al. 2019). Copepods, a known host of *Chytriodinium* are more abundant in the surface waters of the Harbour (da Silva et al. 2022). *Syndiniales* on the other hand is a more flexible parasite infecting other dinoflagellates, radiolarians, and copepods in coastal environments (Bråte et al. 2012; Chambouvet et al. 2008; Guillou et al. 2008; Skovgaard 2014). The increased syndinian parasite abundance was also detected in stratified environments such as fjord and estuarine systems (Sehein et al. 2022; Torres-Beltrán et al. 2018) corroborating the results. The distribution and abundance of dinoflagellates and other non-phototrophic hosts inhabiting the water column of the Harbour influence the distribution and parasitic activities of both *Syndiniales* and *Chytriodinium*. Therefore, these predatory and parasitic groups may play role in the food web of the Harbour and in the organic carbon cycling by cell lysis.

Ciliates are an important component of marine plankton communities. They encompass heterotrophic and mixotrophic organisms and as such they can function as primary producers, consumers and as food for metazoans, playing an important role in linking energy transfer through trophic levels (Calbet & Saiz 2005; Doherty et al. 2010; Pierce & Turner 1992; Santoferrara, Alder & McManus 2017). These groups are typically abundant in the surface of estuarine and freshwater habitats (Hadas & Berman 1998; Jiang et al. 2021; Muylaert et al. 2000; Urrutxurtu, Orive & de la Sota 2003), and in this study the ciliates *Choreotrichia* and *Oligotrichia* (*Strombidium*) were characteristic of the plankton community found in the surface waters. Their predominance in the surface waters of Macquarie Harbour may be due to a range of factors, including differences in salinity and temperature, availability of humic matter, or prey availability (e.g., *cyanobacteria, small dinoflagellates and flagellates)* (Carlsson & Granéli 1993; Hughes et al. 2021; Jiang et al. 2021; Yang et al. 2020). Ciliates inhabiting the water column of the Harbour are likely to make a crucial contribution to nutrient cycling and plankton dynamics.

### Protists as Bioindicators for Aquaculture Activity

Despite the strong depth stratification effects, there were differences in abundance for some ASVs between the lease and control sites. Nineteen protistan ASVs were repeatedly found across the water column, and the majority of these ASVs were more abundant at the lease than the control sites. These included taxa related to *Chrysophyceae*, *Gymnodiniphycidae* (*Gyrodinium*), *Cryptomonadales* and *Ciliophora* (*Philasterides armatalis, Plagiopylida,* and *Strombidium*).

The higher abundance of *Chrysophyceae* and *Gyrodinium* ASVs in the leases is supported by previous studies reporting similar groups being affected by the presence of nutrient inputs, including from aquaculture (Findlay et al. 2009; Jimenez 1993; Jones et al. 1982; Mahmoud & Magdy 2021; Xiaoshan et al. 2004). For example, Findlay et al. (2009) detected increased abundance of chrysophytes associated with high levels of N and P in a lake heavily stocked with rainbow trout, and *Gyrodinium* species have been observed to thrive in estuaries with high nutrient inputs from anthropogenic sources such as aquaculture (Wang et al. 2017). Chrysophytes and *Gyrodinium* species have also more generally been associated to humic content and nutrient enrichment (Li, Stoecker & Coats 2000; Pålsson & Granéli 2004) and used as biological indicators to detect environmental changes (Bellinger & Sigee 2015; Bock et al. 2022; Gauthier et al. 2021; Gökçe 2016; Siver 1995; Xu et al. 2010). The abundance of chrysophytes and *Gyrodinium* species could be related to the presence of humic matter and nutrients concentrations due to riverine and fish farms inputs in the Harbour. Considering the environmental characteristic of the Harbour, these taxa are well suited to detect changes in this environment and further studies are needed to confirm them as potential bioindicators for aquaculture nutrient enrichment.

Among the “other protists” group, most of the ASVs that were differentially more abundant at the lease relative to the control sites were related to the ciliates. Characteristics such as high sensitivity to environmental change, short generation times, and ubiquity make ciliates good candidates as bioindicators of anthropogenic pressures. As morphological classification is often problematic for this group, the advance of molecular techniques has emphasized the discovery and knowledge of potential indicator taxa (Warren et al. 2017). Through molecular techniques, ciliates have now been used to assess the biological conditions of diverse aquatic environments, with variation in oxygen concentration, water masses, light and nutrient enrichment as major factors affecting their composition and distribution (Babko et al. 2022; Dias, Wieloch & D’Agosto 2008; Kulaš et al. 2021; Pawlowski et al. 2016). The methods applied in this study demonstrated that feature selection analyses and amplicon sequencing techniques could detect a depth stratified response of the pelagic protist communities to aquaculture. Previous studies have demonstrated that aquaculture activities lead to nutrient enrichment in the surrounding environment, potentially stimulating bacteria and phytoplankton growth (La Rosa et al. 2002; Olsen et al. 2014; Pitta et al. 2005; Tsagaraki et al. 2013). Since ciliates feed on bacteria, algae, heterotrophic flagellates and other protists (Pierce & Turner 1992) it is therefore not unexpected that ciliates can be found abundant surrounding fish farms.

The ciliate ASVs that were characteristic of the lease sites were assigned to *Philasterides armatalis*, *Plagiopylida*, and *Strombidium*. This is consistent with previous studies that have reported the free-living *Philasterides* and *Strombidium* in mariculture waters in China and Cyprus (Song 2000; Tsagaraki et al. 2013; Wu et al. 2018). *Plagiopylida* are ciliates found in anaerobic marine habitats containing high concentration of H_2_S (Edgcomb & Pachiadaki 2014; Rotterová et al. 2022). The abundance of these ciliates is probably related to nutrient enrichment leading to a highly reduced environment commonly found bellow fish farms in the benthos (Bissett, Bowman & Burke 2006; Keeley, Forrest & Macleod 2013; Quero et al. 2020). These conditions favor specific microorganisms capable of thriving in environments with low oxygen and/or high amount of organic matter and dissolved nutrients (Choi et al. 2018; La Rosa et al. 2002). It is likely that the presence of these abundant ciliates at the leases in this study is in response to the greater availability of organic material and low oxygen conditions at the bottom of the water column.

The results from this study are supported by Stoeck et al. (2018) who found that ciliates, dinoflagellates and chrysophytes could all be used to distinguish sites impacted by feed-additive aquaculture. Similarly, Pitta et al. (2005) also observed changes in the abundance of ciliates and phytoplankton in bottom waters near seabream and seabass fish cages in the Mediterranean Sea (Pitta et al. 2005), and at salmon farms in Chile, Norway and Scotland (Aranda et al. 2015; Forster et al. 2019; Fruhe et al. 2021).

Other important factors that should be considered when analysing the effects of aquaculture on the microbial community would be the management practices (e.g., feed input rate), production (e.g., stock density, faeces), hydrodynamics (e.g., shallow or deeper waters) and location (e.g., south or north of the harbour) of that specific aquaculture lease area (Belias et al. 2003; Jansen et al. 2016; Sanderson et al. 2008; Sarà et al. 2007; Skogen et al. 2009; Tsagaraki et al. 2011; Wu et al. 2018). For example, among leases, there was some variation in the ASVs distribution along the distance gradient from fish cages. This may be related to the location of each lease and/or the local hydrodynamical features. To illustrate, Lease 1 is located further south where the influence of the Gordon River is more accentuated (Teasdale et al. 2003) whereas Lease 2 is further north, near the deepest basin that is approximately 52 m deep. The region where Lease 1 is in an area with lower water exchange and may be receiving a higher concentration of organic carbon and other nutrients compared to Lease 2 due to river and water circulation patterns within the harbour (Hartstein et al. 2019; Kirkpatrick, Kriwoken & Styger 2019; Terry 2001). More detailed studies on environmental seasonality and its effects on the microbial community would elucidate possible fluctuations in the composition and function of microorganisms, in addition to obtaining a clearer picture of the possible impacts of aquaculture in the harbour.

## Conclusion

The study revealed highly vertical stratified communities of protists across the water column of the Harbour. The results also demonstrated that it was possible to detect differences between lease and control sites in the pelagic protist communities, despite the strong depth effect controlling the distribution. Vertical differences in relative abundance of taxa were related to nutrient concentrations and environmental changes in oxygen concentration, light, and salinity. The use of molecular methods such as amplicon sequencing demonstrated potential for monitoring aquaculture effects in the water column by identifying key groups (i.e., chrysophytes, *Gyrodinium* and ciliates) that had higher relative abundance in proximity to aquaculture. While this data was able to identify potential bioindicators of aquaculture activity, it also provided invaluable insight into the general ecology and distribution of protist communities within Macquarie Harbour and the role of environmental characteristics in shaping these communities. The location and operational practices of fish farms may also be important factors driving the distribution of pelagic protists. These findings could be further verified through a seasonal survey encompassing the production cycle to assess the spatial and temporal variation of protists community. In summary, this study provides important first insights of aquaculture effects on the pelagic protist community within a highly stratified semi-enclosed embayment. Also, the results highlighted the potential use of these microorganisms as bioindicator for aquaculture activity using the 18S rRNA amplicon sequencing.

## References

á Norði, G, Glud, RN, Gaard, E & Simonsen, K 2011, ‘Environmental impacts of coastal fish farming: carbon and nitrogen budgets for trout farming in Kaldbaksfjør7%%FONT_ERR%%ur (Faroe Islands)’, Marine Ecology Progress Series, vol. 431, pp. 223–241.

Aitchison, J 1986, The Statistical Analysis of Compositional Data, London; New York: Chapman and Hall.

Alongi, DM, Chong, VC, Dixon, P, Sasekumar, A & Tirendi, F 2003, ‘The influence of fish cage aquaculture on pelagic carbon flow and water chemistry in tidally dominated mangrove estuaries of peninsular Malaysia’, Marine Environmental Research, vol. 55, no. 4, pp. 313–333.

Anderson, SR & Harvey, EL 2020, ‘Temporal Variability and Ecological Interactions of Parasitic Marine Syndiniales in Coastal Protist Communities’, mSphere, vol. 5, no. 3.

Andrei, AS, Robeson, MS, 2nd, Baricz, A, Coman, C, Muntean, V, Ionescu, A, Etiope, G, Alexe, M, Sicora, CI, Podar, M & Banciu, HL 2015, ‘Contrasting taxonomic stratification of microbial communities in two hypersaline meromictic lakes’, ISME J, vol. 9, no. 12, pp. 2642–2656.

Apotheloz-Perret-Gentil, L, Bouchez, A, Cordier, T, Cordonier, A, Gueguen, J, Rimet, F, Vasselon, V & Pawlowski, J 2021, ‘Monitoring the ecological status of rivers with diatom eDNA metabarcoding: A comparison of taxonomic markers and analytical approaches for the inference of a molecular diatom index’, Mol Ecol, vol. 30, no. 13, pp. 2959–2968.

Appleyard, S, Abell, G & Watson, R 2013, Tackling microbial related issues in cultured shellfish via integrated molecular and water chemistry approaches. Seafood CRC Final Report (2011/729) (pp. 89). viewed 20 August 2019, <http://frdc.com.au/Archived-Reports/FRDC%20Projects/2011-729-DLD.pdf>.

Aranda, CP, Valenzuela, C, Matamala, Y, Godoy, FA & Aranda, N 2015, ‘Sulphur-cycling bacteria and ciliated protozoans in a Beggiatoaceae mat covering organically enriched sediments beneath a salmon farm in a southern Chilean fjord’, Mar Pollut Bull, vol. 100, no. 1, pp. 270–278.

Asami, H, Aida, M & Watanabe, K 2005, ‘Accelerated sulfur cycle in coastal marine sediment beneath areas of intensive shellfish aquaculture’, Appl Environ Microbiol, vol. 71, no. 6, pp. 2925–2933.

Azam, F, Fenchel, T, Field, J, Grey, J, Meyer-Reil, L & Thingstad, F 1983, ‘The ecological role of water-column microbes’, Mar. ecol. Prog. ser, vol. 10, pp. 257–263.

Azam, F & Malfatti, F 2007, ‘Microbial structuring of marine ecosystems’, Nat Rev Microbiol, vol. 5, no. 10, pp. 782–791.

Babko, R, Kuzmina, T, Danko, Y, Pliashechnyk, V, Szulżyk-Cieplak, J, Łazuka, E, Zaburko, J & Łagód, G 2022, ‘Spatial Distribution of Ciliate Assemblages in a Shallow Floodplain Lake with an Anaerobic Zone’, Water, vol. 14, no. 6, p. 898.

Bartl, I, Liskow, I, Schulz, K, Umlauf, L & Voss, M 2018, ‘River plume and bottom boundary layer – Hotspots for nitrification in a coastal bay?’, Estuarine, Coastal and Shelf Science, vol. 208, pp. 70–82.

Basu, S & Mackey, K 2018, ‘Phytoplankton as key mediators of the biological carbon pump: Their responses to a changing climate’, Sustainability, vol. 10, no. 3, p. 869.

Behnke, A, Barger, KJ, Bunge, J & Stoeck, T 2010, ‘Spatio-temporal variations in protistan communities along an O2/H2S gradient in the anoxic Framvaren Fjord (Norway)’, FEMS Microbiology Ecology, vol. 72, no. 1, pp. 89–102.

Belias, CV, Bikas, VG, Dassenakis, MJ & Scoullos, MJ 2003, ‘Environmental impacts of coastal aquaculture in eastern mediterranean bays the case of astakos gulf, Greece’, Environmental Science and Pollution Research, vol. 10, no. 5, p. 287.

Bellinger, EG & Sigee, DC 2015, Freshwater algae: identification, enumeration and use as bioindicators, John Wiley & Sons.

Bissett, A, Bowman, J & Burke, C 2006, ‘Bacterial diversity in organically-enriched fish farm sediments’, FEMS Microbiol Ecol, vol. 55, no. 1, pp. 48–56.

Bock, C, Olefeld, JL, Vogt, JC, Albach, DC & Boenigk, J 2022, ‘Phylogenetic and functional diversity of Chrysophyceae in inland waters’, Organisms Diversity & Evolution, pp. 1–15.

Bråte, J, Krabberød, AK, Dolven, JK, Ose, RF, Kristensen, T, Bjørklund, KR & Shalchian-Tabrizi, K 2012, ‘Radiolaria associated with large diversity of marine alveolates’, Protist, vol. 163, no. 5, pp. 767–777.

Burkholder, JM, Glasgow, HB & Deamer-Mella, N 2001, ‘Overview and present status of the toxic Pfiesteria complex (Dinophyceae)’, Phycologia, vol. 40, no. 3, pp. 186–214.

Burkholder, JM, Gordon, AS, Moeller, PD, Mac Law, J, Coyne, KJ, Lewitus, AJ, Ramsdell, JS, Marshall, HG, Deamer, NJ & Cary, SC 2005, ‘Demonstration of toxicity to fish and to mammalian cells by Pfiesteria species: comparison of assay methods and strains’, Proceedings of the National Academy of Sciences, vol. 102, no. 9, pp. 3471–3476.

Burridge, L, Weis, JS, Cabello, F, Pizarro, J & Bostick, K 2010, ‘Chemical use in salmon aquaculture: A review of current practices and possible environmental effects’, Aquaculture, vol. 306, no. 1, pp. 7–23.

Buschmann, A, Riquelme, V, Hernandezgonzalez, M, Varela, D, Jimenez, J, Henriquez, L, Vergara, P, Guinez, R & Filun, L 2006, ‘A review of the impacts of salmonid farming on marine coastal ecosystems in the southeast Pacific’, ICES Journal of Marine Science, vol. 63, no. 7, pp. 1338–1345.

Cabello-Yeves, PJ, Callieri, C, Picazo, A, Mehrshad, M, Haro-Moreno, JM, Roda-Garcia, JJ, Dzhembekova, N, Slabakova, V, Slabakova, N, Moncheva, S & Rodriguez-Valera, F 2021, ‘The microbiome of the Black Sea water column analyzed by shotgun and genome centric metagenomics’, Environ Microbiome, vol. 16, no. 1, p. 5.

Calbet, A & Saiz, E 2005, ‘The ciliate-copepod link in marine ecosystems’, Aquatic Microbial Ecology, vol. 38, no. 2, pp. 157–167.

Callahan, BJ, McMurdie, PJ, Rosen, MJ, Han, AW, Johnson, AJ & Holmes, SP 2016, ‘DADA2: High-resolution sample inference from Illumina amplicon data’, Nat Methods, vol. 13, no. 7, pp. 581–583.

Cao, Q, Sun, X, Rajesh, K, Chalasani, N, Gelow, K, Katz, B, Shah, VH, Sanyal, AJ & Smirnova, E 2021, ‘Effects of Rare Microbiome Taxa Filtering on Statistical Analysis’, Frontiers in Microbiology, vol. 11.

Capone, DG, Bronk, DA, Mulholland, MR & Carpenter, EJ 2008, Nitrogen in the marine environment, Elsevier.

Carlsson, P & Granéli, E 1993, ‘Availability of humic bound nitrogen for coastal phytoplankton’, Estuarine, Coastal and Shelf Science, vol. 36, no. 5, pp. 433–447.

Carpenter, PD, Butler, ECV, Higgins, HW, Mackey, DJ & Nichols, PD 1991, ‘Chemistry of trace elements, humic substances and sedimentary organic matter in Macquarie Harbour, Tasmania’, Marine and Freshwater Research, vol. 42, no. 6.

Chambouvet, A, Morin, P, Marie, D & Guillou, L 2008, ‘Control of toxic marine dinoflagellate blooms by serial parasitic killers’, Science, vol. 322, no. 5905, pp. 1254–1257.

Chawla, NV, Bowyer, KW, Hall, LO & Kegelmeyer, WP 2002, ‘SMOTE: synthetic minority over-sampling technique’, Journal of artificial intelligence research, vol. 16, pp. 321–357.

Choi, A, Cho, H, Kim, B, Kim, HC, Jung, RH, Lee, WC & Hyun, JH 2018, ‘Effects of finfish aquaculture on biogeochemistry and bacterial communities associated with sulfur cycles in highly sulfidic sediments’, Aquaculture Environment Interactions, vol. 10, pp. 413–427.

Cloern, JE 2001, ‘Our evolving conceptual model of the coastal eutrophication problem’, Marine Ecology Progress Series, vol. 210, pp. 223–253.

Cordier, T, Forster, D, Dufresne, Y, Martins, CIM, Stoeck, T & Pawlowski, J 2018, ‘Supervised machine learning outperforms taxonomy-based environmental DNA metabarcoding applied to biomonitoring’, Mol Ecol Resour, vol. 18, no. 6, pp. 1381–1391.

Cordier, T, Lanzen, A, Apotheloz-Perret-Gentil, L, Stoeck, T & Pawlowski, J 2019, ‘Embracing Environmental Genomics and Machine Learning for Routine Biomonitoring’, Trends Microbiol, vol. 27, no. 5, pp. 387–397.

Costa, AED, Santana, JR & Neumann-Leitao, S 2018, ‘Changes in microplanktonic protists assemblages promoted by the thermocline induced stratification around an oceanic archipelago’, Anais da Academia Brasileira de Ciências, vol. 90, pp. 2249–2266.

Crawford, CM, Macleod, CKA & Mitchell, IM 2003, ‘Effects of shellfish farming on the benthic environment’, Aquaculture, vol. 224, no. 1-4, pp. 117–140.

Cresswell, G, Edwards, R & Barker, B 1989, ’Macquarie Harbour, Tasmania-seasonal oceanographic surveys in 1985’, in Papers and Proceedings of the Royal Society of Tasmania, vol. 123, pp. 63–66.

da Silva, RR, White, C, Bowman, J, Bodrossy, L, Bissett, A, Revill, A, Eriksen, R & Ross, D 2022, ‘The importance of environmental parameters and mixing zone in shaping estuarine microbial communities along a freshwater-marine gradient’, bioRxiv.

Da Silva, RRP, White, CA, Bowman, JP, Raes, E, Bisset, A, Chapman, C, Bodrossy, L & Ross, DJ 2021, ‘Environmental influences shaping microbial communities in a low oxygen, highly stratified marine embayment’, Aquatic Microbial Ecology, vol. 87, pp. 185–203.

Dagenais-Bellefeuille, S & Morse, D 2013, ‘Putting the N in dinoflagellates’, Front Microbiol, vol. 4, p. 369.

Das, PB, Gauns, M & Naqvi, SWA 2019, ‘Ecological diversity of planktonic protists in spatial regimes of the Arabian Sea revealed through next-generation sequencing’, Regional Studies in Marine Science, vol. 25.

DeLong, EF, Preston, CM, Mincer, T, Rich, V, Hallam, SJ, Frigaard, N-U, Martinez, A, Sullivan, MB, Edwards, R, Brito, BR, Chisholm, SW & Karl, DM 2006, ’Community Genomics Among Stratified Microbial Assemblages in the Ocean’s Interior’, Science, vol. 311, no. 5760, pp. 496–503.

Dias, R, Wieloch, A & D’Agosto, M 2008, ‘The influence of environmental characteristics on the distribution of ciliates (Protozoa, Ciliophora) in an urban stream of southeast Brazil’, Brazilian Journal of Biology, vol. 68, pp. 287–295.

Doherty, M, Tamura, M, Vriezen, JA, McManus, GB & Katz, LA 2010, ‘Diversity of Oligotrichia and Choreotrichia ciliates in coastal marine sediments and in overlying plankton’, Applied and Environmental Microbiology, vol. 76, no. 12, pp. 3924–3935.

Dokulil, MT 1994, ’Environmental control of phytoplankton productivity in turbulent turbid systems’, in *Phytoplankton in Turbid Environments: Rivers and Shallow Lakes*, Springer, pp. 65–72.

Dorigo, U, Bérard, A & Humbert, JF 2002, ‘Comparison of Eukaryotic Phytobenthic Community Composition in a Polluted River by Partial 18S rRNA Gene Cloning and Sequencing’, Microbial Ecology, vol. 44, no. 4, pp. 372–380.

Dully, V, Balliet, H, Frühe, L, Däumer, M, Thielen, A, Gallie, S, Berrill, I & Stoeck, T 2021, ‘Robustness, sensitivity and reproducibility of eDNA metabarcoding as an environmental biomonitoring tool in coastal salmon aquaculture – An inter-laboratory study’, Ecological Indicators, vol. 121.

Edgcomb, VP & Pachiadaki, M 2014, ‘Ciliates along oxyclines of permanently stratified marine water columns’, Journal of Eukaryotic Microbiology, vol. 61, no. 4, pp. 434–445.

Elizondo-Patrone, C, Hernández, K, Yannicelli, B, Olsen, LM & Molina, V 2015, ‘The response of nitrifying microbial assemblages to ammonium (NH4+) enrichment from salmon farm activities in a northern Chilean Fjord’, Estuarine, Coastal and Shelf Science, vol. 166, pp. 131–142.

P Department of Primary Industries, Water and Environment 2012, Macquarie Harbour Marine Farming Development Plan October 2005 by EPA, Environmental Protection Authority, <https://nre.tas.gov.au/Documents/Mac-Harbour-MFDP-October-2005.pdf>.

EP AUTHORITY 2017, *EPA Director’s statement* by EPA, EPA, <https://epa.tas.gov.au/Documents/Macquarie%20Harbour%20Statement%20of%20Reasons%20-%2031%20May2017%20_%20Final.pdf>.

Fernandes, AD, Macklaim, JM, Linn, TG, Reid, G & Gloor, GB 2013, ‘ANOVA-like differential expression (ALDEx) analysis for mixed population RNA-Seq’, PLOS ONE, vol. 8, no. 7, p. e67019.

Findlay, DL, Smith, R, Podemski, CL & Kasian, SEM 2009, ‘Aquaculture impacts on the algal and bacterial communities in a small boreal forest lakeThis paper is part of the series “Forty Years of Aquatic Research at the Experimental Lakes Area”’, Canadian Journal of Fisheries and Aquatic Sciences, vol. 66, no. 11, pp. 1936–1948.

Forster, D, Filker, S, Kochems, R, Breiner, HW, Cordier, T, Pawlowski, J & Stoeck, T 2019, ‘A Comparison of Different Ciliate Metabarcode Genes as Bioindicators for Environmental Impact Assessments of Salmon Aquaculture’, J Eukaryot Microbiol, vol. 66, no. 2, pp. 294–308.

Fruhe, L, Cordier, T, Dully, V, Breiner, HW, Lentendu, G, Pawlowski, J, Martins, C, Wilding, TA & Stoeck, T 2021, ‘Supervised machine learning is superior to indicator value inference in monitoring the environmental impacts of salmon aquaculture using eDNA metabarcodes’, Mol Ecol, vol. 30, no. 13, pp. 2988–3006.

Fulton, RS 1984, ‘Distribution and community structure of estuarine copepods’, Estuaries, vol. 7, no. 1, pp. 38–50.

Gardner, WS, Newell, SE, McCarthy, MJ, Hoffman, DK, Lu, K, Lavrentyev, PJ, Hellweger, FL, Wilhelm, SW, Liu, Z, Bruesewitz, DA & Paerl, HW 2017, ‘Community Biological Ammonium Demand: A Conceptual Model for Cyanobacteria Blooms in Eutrophic Lakes’, Environ Sci Technol, vol. 51, no. 14, pp. 7785–7793.

Gauthier, J, Walsh, D, Selbie, DT, Bourgeois, A, Griffiths, K, Domaizon, I & GregoryLEaves, I 2021, ‘Evaluating the congruence between DNA Lbased and morphological taxonomic approaches in water and sediment trap samples: Analyses of a 36Lmonth time series from a temperate monomictic lake’, Limnology and Oceanography, vol. 66, no. 8, pp. 3020–3039.

Glasgow, HB, Burkholder, JM, Mallin, MA, Deamer-Melia, NJ & Reed, RE 2001, ‘Field ecology of toxic Pfiesteria complex species and a conservative analysis of their role in estuarine fish kills’, Environmental Health Perspectives, vol. 109, no. suppl 5, pp. 715–730.

Glibert, PM & Terlizzi, DE 1999, ‘Cooccurrence of elevated urea levels and dinoflagellate blooms in temperate estuarine aquaculture ponds’, Applied and Environmental Microbiology, vol. 65, no. 12, pp. 5594–5596.

Gloor, G 2016, ‘CoDaSeq: Analyzing HTS using compositional data analysis’, F1000Research, vol. 5.

Gloor, GB, Macklaim, JM, Pawlowsky-Glahn, V & Egozcue, JJ 2017, ‘Microbiome Datasets Are Compositional: And This Is Not Optional’, Front Microbiol, vol. 8, p. 2224.

Gökçe, D 2016, ‘Algae as an indicator of water quality’, Algae-Organisms for Imminent Biotechnology, pp. 81–101.

Gómez, F 2012, ‘A quantitative review of the lifestyle, habitat and trophic diversity of dinoflagellates (Dinoflagellata, Alveolata)’, Systematics and Biodiversity, vol. 10, no. 3, pp. 267–275.

Gómez, F & Skovgaard, A 2015, ‘Molecular phylogeny of the parasitic dinoflagellate Chytriodinium within the Gymnodinium clade (Gymnodiniales, Dinophyceae)’, Journal of Eukaryotic Microbiology, vol. 62, no. 3, pp. 422–425.

Grasshoff, K, Kremling, K & Ehrhardt, M 2009, Methods of seawater analysis, John Wiley & Sons.

Greenwell, BM & Boehmke, BC 2020, ‘Variable Importance Plots—An Introduction to the vip Package.’, The R Journal, vol. 12, no. 1, pp. 343–366.

Guillou, L, Viprey, M, Chambouvet, A, Welsh, RM, Kirkham, AR, Massana, R, Scanlan, DJ & Worden, AZ 2008, ‘Widespread occurrence and genetic diversity of marine parasitoids belonging to Syndiniales (Alveolata)’, Environ Microbiol, vol. 10, no. 12, pp. 3349–3365.

Hadas, O & Berman, T 1998, ‘Seasonal abundance and vertical distribution of Protozoa (flagellates, ciliates) and bacteria in Lake Kinneret, Israel’, Aquatic Microbial Ecology, vol. 14, no. 2, pp. 161–170.

Hamilton, NE & Ferry, M 2018, ‘ggtern: Ternary diagrams using ggplot2’, Journal of statistical software, vol. 87, no. 1, pp. 1–17.

Hartstein, ND, Maxey, JD, Loo, JCH & Then, AY-H 2019, ‘Drivers of deep water renewal in Macquarie Harbour, Tasmania’, Journal of Marine Systems, vol. 199, no. 103226.

Hashimoto, Y, Okaichi, T, Dang, LD & Noguchi, T 1968, ‘Glenodinine, an ichthyotoxic substance produced by a dinoflagellate, Peridinium polonicum’, Bulletin of the Japanese Society of Scientific Fisheries, vol. 34, no. 6, pp. 528–534.

Havskum, H & Riemann, B 1996, ‘Ecological importance of bacterivorous, pigmented flagellates (mixotrophs) in the Bay of Aarhus, Denmark’, Marine Ecology Progress Series, vol. 137, pp. 251–263.

Hawinkel, S, Mattiello, F, Bijnens, L & Thas, O 2019, ‘A broken promise: microbiome differential abundance methods do not control the false discovery rate’, Brief Bioinform, vol. 20, no. 1, pp. 210–221.

He, X, Sutherland, TF, Pawlowski, J & Abbott, CL 2019, ‘Responses of foraminifera communities to aquaculture-derived organic enrichment as revealed by environmental DNA metabarcoding’, Mol Ecol.

Hughes, EA, Maselli, M, Sørensen, H & Hansen, PJ 2021, ‘Metabolic reliance on photosynthesis depends on both irradiance and prey availability in the mixotrophic ciliate, Strombidium cf. basimorphum’, Frontiers in Microbiology, vol. 12, p. 1577.

Husa, V, Kutti, T, Ervik, A, Sjøtun, K, Hansen, PK & Aure, J 2013, ‘Regional impact from fin-fish farming in an intensive production area (Hardangerfjord, Norway)’, Marine Biology Research, vol. 10, no. 3, pp. 241–252.

Iwataki, M 2008, ‘Taxonomy and identification of the armored dinoflagellate genus Heterocapsa (Peridiniales, Dinophyceae)’, Plankton and Benthos Research, vol. 3, no. 3, pp. 135–142.

Jacquemot, L, Kalenitchenko, D, Matthes, LC, Vigneron, A, Mundy, CJ, Tremblay, J-É & Lovejoy, C 2021, ‘Protist communities along freshwater–marine transition zones in Hudson Bay (Canada)’, Elementa: Science of the Anthropocene, vol. 9, no. 1.

Jansen, HM, Broch, OJ, Bannister, R, Cranford, P, Handå, A, Husa, V, Jiang, Z, Strohmeier, T & Strand, Ø 2018, ‘Spatio-temporal dynamics in the dissolved nutrient waste plume from Norwegian salmon cage aquaculture’, Aquaculture Environment Interactions, vol. 10, pp. 385–399.

Jansen, HM, Reid, GK, Bannister, RJ, Husa, V, Robinson, SMC, Cooper, JA, Quinton, C & Strand, Ø 2016, ‘Discrete water quality sampling at open-water aquaculture sites: limitations and strategies’, Aquaculture Environment Interactions, vol. 8, pp. 463–480.

Jiang, C, Liu, B, Zhang, J, Gu, S, Liu, Z, Wang, X, Chen, K, Xiong, J, Lu, Y & Miao, W 2021, ‘Diversity and seasonality dynamics of ciliate communities in four estuaries of Shenzhen, China (South China Sea)’, Journal of Marine Science and Engineering, vol. 9, no. 3, p. 260.

Jiang, ZB, Chen, QZ, Zeng, JN, Liao, YB, Shou, L & Liu, J 2012, ‘Phytoplankton community distribution in relation to environmental parameters in three aquaculture systems in a Chinese subtropical eutrophic bay’, Marine Ecology Progress Series, vol. 446, pp. 73–89.

Jimenez, R 1993, ‘Ecological factors related to Gyrodinium instriatum bloom in the inner estuary of the Gulf of Guayaquil’, Toxic Phytoplankton Blooms in the Sea, pp. 257–262.

Johnsen, G, Nelson, NB, Jovine, RV & Prezelin, BB 1994, ‘dinoflagellates, Prorocentrum minimum and Heterocapsa pygmaea’, Marine Ecology Progress Series, vol. 114, pp. 245–258.

Jones, K, Ayres, P, Bullock, A, Roberts, R & Tett, P 1982, ‘A red tide of gyrod inium aurelum in sea lochs of the firth of clyde and associated mortality of pond-reared salmon’, Journal of the Marine Biological Association of the United Kingdom, vol. 62, no. 4, pp. 771–782.

Keeley, NB, Forrest, BM & Macleod, CK 2013, ‘Novel observations of benthic enrichment in contrasting flow regimes with implications for marine farm monitoring and management’, Marine Pollution Bulletin, vol. 66, no. 1-2, pp. 105–116.

King, RD & Tyler, PA 1982, ‘Downstream effects of the Gorden River Power Development, south-west Tasmania’, Marine and Freshwater Research, vol. 33, no. 3, pp. 431–442.

Kirkpatrick, JB, Kriwoken, LK & Styger, J 2019, ‘The reverse precautionary principle: science, the environment and the salmon aquaculture industry in Macquarie Harbour, Tasmania, Australia’, Pacific Conservation Biology, vol. 25, no. 1.

Koehnken, L 2005, ‘Overview of Water Quality in Macquarie Harbour and Assessment of Risks due to Copper Levels’.

Kuhn, M & Wickham, H 2020, *Tidymodels: a collection of packages for modeling and machine learning using tidyverse principles*. https://www.tidymodels.org.

Kulaš, A, Gulin, V, Matoničkin Kepčija, R, Žutinić, P, Sertić Perić, M, Orlić, S, Kajan, K, Stoeck, T, Lentendu, G, Čanjevac, I, Martinić, I & Gligora Udovič, M 2021, ‘Ciliates (Alveolata, Ciliophora) as bioindicators of environmental pressure: A karstic river case’, Ecological Indicators, vol. 124.

Kurobe, T, Lehman, PW, Hammock, BG, Bolotaolo, MB, Lesmeister, S & Teh, SJ 2018, ‘Biodiversity of cyanobacteria and other aquatic microorganisms across a freshwater to brackish water gradient determined by shotgun metagenomic sequencing analysis in the San Francisco Estuary, USA’, PLOS ONE, vol. 13, no. 9, p. e0203953.

La Rosa, T, Mirto, S, Favaloro, E, Savona, B, Sarà, G, Danovaro, R & Mazzola, A 2002, ‘Impact on the water column biogeochemistry of a Mediterranean mussel and fish farm’, Water Research, vol. 36, no. 3, pp. 713–721.

Lallias, D, Hiddink, JG, Fonseca, VG, Gaspar, JM, Sung, W, Neill, SP, Barnes, N, Ferrero, T, Hall, N, Lambshead, PJ, Packer, M, Thomas, WK & Creer, S 2015, ‘Environmental metabarcoding reveals heterogeneous drivers of microbial eukaryote diversity in contrasting estuarine ecosystems’, ISME J, vol. 9, no. 5, pp. 1208–1221.

Laroche, O, Meier, S, Mjøs, SA & Keeley, N 2021, ‘Effects of fish farm activities on the sponge Weberella bursa, and its associated microbiota’, Ecological Indicators, vol. 129.

Leach, TH, Beisner, BE, Carey, CC, Pernica, P, Rose, KC, Huot, Y, Brentrup, JA, Domaizon, I, Grossart, H-P, Ibelings, BW, Jacquet, S, Kelly, PT, Rusak, JA, Stockwell, JD, Straile, D & Verburg, P 2018, ‘Patterns and drivers of deep chlorophyll maxima structure in 100 lakes: The relative importance of light and thermal stratification’, Limnology and Oceanography, vol. 63, no. 2, pp. 628–646.

Legrand, C, Graneli, E & Carlsson, P 1998, ‘Induced phagotrophy in the photosynthetic dinoflagellate Heterocapsa triquetra’, Aquatic Microbial Ecology, vol. 15, no. 1, pp. 65–75.

Li, A, Stoecker, DK & Coats, DW 2000, ‘Mixotrophy in Gyrodinium galatheanum (Dinophyceae): grazing responses to light intensity and inorganic nutrients’, Journal of Phycology, vol. 36, no. 1, pp. 33–45.

Liaw, A & Wiener, M 2002, ‘Classification and regression by randomForest’, R news, vol. 2, no. 3, pp. 18–22.

Love, MI, Huber, W & Anders, S 2014, ‘Moderated estimation of fold change and dispersion for RNA-seq data with DESeq2’, Genome Biol, vol. 15, no. 12, p. 550.

Mahmoud, MAA & Magdy, M 2021, ‘Metabarcoding profiling of microbial diversity associated with trout fish farming’, Sci Rep, vol. 11, no. 1, p. 421.

McKinnon, AD, Trott, LA, Brinkman, R, Duggan, S, Castine, S, O’Leary, RA & Alongi, DM 2010, ‘Seacage aquaculture in a World Heritage Area: the environmental footprint of a Barramundi farm in tropical Australia’, Mar Pollut Bull, vol. 60, no. 9, pp. 1489–1501.

McMurdie, PJ & Holmes, S 2013, ‘phyloseq: an R package for reproducible interactive analysis and graphics of microbiome census data’, PLOS ONE, vol. 8, no. 4, p. e61217.

Mena, C, Reglero, P, Hidalgo, M, Sintes, E, Santiago, R, Martín, M, Moyà, G & Balbín, R 2019, ‘Phytoplankton community structure is driven by stratification in the oligotrophic Mediterranean Sea’, Frontiers in Microbiology, p. 1698.

Mikaelyan, AS, Pautova, LA & Fedorov, AV 2021, ‘Seasonal evolution of deep phytoplankton assemblages in the Black Sea’, Journal of Sea Research, vol. 178, p. 102125.

Moncada, C, Hassenruck, C, Gardes, A & Conaco, C 2019, ‘Microbial community composition of sediments influenced by intensive mariculture activity’, FEMS Microbiol Ecol, vol. 95, no. 2.

Morien, E & Parfrey, L 2018, *SILVA v128 and v132 dada2 formatted 18s “train sets.”*, <10.5281/ZENODO.1447330>.

Muylaert, K, Van Mieghem, R, Sabbe, K, Tackx, M & Vyverman, W 2000, ‘Dynamics and trophic roles of heterotrophic protists in the plankton of a freshwater tidal estuary’, Hydrobiologia, vol. 432, no. 1, pp. 25–36.

Nagasoe, S, Shikata, T, Yamasaki, Y, Matsubara, T, Shimasaki, Y, Oshima, Y & Honjo, T 2010, ‘Effects of nutrients on growth of the red-tide dinoflagellate Gyrodinium instriatum Freudenthal et Lee and a possible link to blooms of this species’, Hydrobiologia, vol. 651, no. 1, pp. 225–238.

Navarro, N, Leakey, RJG & Black, KD 2008, ‘Effect of salmon cage aquaculture on the pelagic environment of temperate coastal waters: seasonal changes in nutrients and microbial community’, Marine Ecology Progress Series, vol. 361, pp. 47–58.

Needham, DM & Fuhrman, JA 2016, ‘Pronounced daily succession of phytoplankton, archaea and bacteria following a spring bloom’, Nat Microbiol, vol. 1, p. 16005.

Oikonomou, A, Filker, S, Breiner, HW & Stoeck, T 2015, ‘Protistan diversity in a permanently stratified meromictic Lake (Lake a latsee, SW G ermany)’, Environmental Microbiology, vol. 17, no. 6, pp. 2144–2157.

Oksanen, JF, Blanchet, G, Friendly, M, Kindt, R, Legendre, P, McGlinn, D, Peter, R & Minchin, R 2018, ‘O’Hara; Gavin, L.; et al’, The Vegan Community Ecology Package. Available online: https://cran.r-project.org/web/packages/vegan/ *(accessed on 1 March 2019)*.

Olsen, LM, Hernández, KL, Van Ardelan, M, Iriarte, JL, Sánchez, N, González, HE, Tokle, N & Olsen, Y 2014, ‘Responses in the microbial food web to increased rates of nutrient supply in a southern Chilean fjord: possible implications of cage aquaculture’, Aquaculture Environment Interactions, vol. 6, no. 1, pp. 11–27.

Palarea-Albaladejo, J & Martín-Fernández, JA 2015, ‘zCompositions—R package for multivariate imputation of left-censored data under a compositional approach’, Chemometrics and Intelligent Laboratory Systems, vol. 143, pp. 85–96.

Pålsson, C & Granéli, W 2004, ‘Nutrient limitation of autotrophic and mixotrophic phytoplankton in a temperate and tropical humic lake gradient’, Journal of Plankton Research, vol. 26, no. 9, pp. 1005–1014.

Parris, DJ, Ganesh, S, Edgcomb, VP, DeLong, EF & Stewart, FJ 2014, ‘Microbial eukaryote diversity in the marine oxygen minimum zone off northern Chile’, Front Microbiol, vol. 5, p. 543.

Pawlowski, J, Lejzerowicz, F, Apotheloz-Perret-Gentil, L, Visco, J & Esling, P 2016, ‘Protist metabarcoding and environmental biomonitoring: Time for change’, Eur J Protistol, vol. 55, no. Pt A, pp. 12–25.

Pierce, RW & Turner, JT 1992, ‘Ecology of planktonic ciliates in marine food webs’, Rev. Aquat. Sci, vol. 6, no. 2, pp. 139–181.

Pitta, P, Apostolaki, ET, Giannoulaki, M & Karakassis, I 2005, ‘Mesoscale changes in the water column in response to fish farming zones in three coastal areas in the Eastern Mediterranean Sea’, Estuarine, Coastal and Shelf Science, vol. 65, no. 3, pp. 501–512.

Pomeroy, LR, leB. WILLIAMS, PJ, Azam, F & Hobbie, JE 2007, ‘The microbial loop’, Oceanography, vol. 20, no. 2, pp. 28–33.

Prabhudessai, S, Vishal, C & Rivonker, C 2019, ‘Biotic interaction as the triggering factor for blooms under favourable conditions in tropical estuarine systems’, Environmental monitoring and assessment, vol. 191, no. 2, pp. 1–11.

Prakash, Aa & Rashid, M 1968, ‘INFLUENCE OF HUMIC SUBSTANCES ON THE GROWTH OF MARINE PHYTOPLANKTON: DINOFLAGELLATES 1’, Limnology and Oceanography, vol. 13, no. 4, pp. 598–606.

Quero, GM, Ape, F, Manini, E, Mirto, S & Luna, GM 2020, ‘Temporal Changes in Microbial Communities Beneath Fish Farm Sediments Are Related to Organic Enrichment and Fish Biomass Over a Production Cycle’, Frontiers in Marine Science, vol. 7.

R Core Team 2021, *R: A language and environment for statistical computing. R Foundation for Statistical Computing, Vienna, Austria. URL* https://www.R-project.org/.

Raes, EJ, Bodrossy, L, van de Kamp, J, Bissett, A, Ostrowski, M, Brown, MV, Sow, SLS, Sloyan, B & Waite, AM 2018, ‘Oceanographic boundaries constrain microbial diversity gradients in the South Pacific Ocean’, Proc Natl Acad Sci U S A, vol. 115, no. 35, pp. E8266–E8275.

Revill, AT, Ross, Ja & Thompson, PA 2016, *Investigating Dissolved Oxygen Drawdown in Macquarie Harbour. CSIRO, Australia. Report to Huon Aquaculture*, Institute for Marine and Antarctic Studies (IMAS), Hobart, viewed 16 September 2019, <https://www.huonaqua.com.au/wp-content/uploads/2015/08/Appendix-4-part-4.pdf>.

Roset, J, Gibello, A, Aguayo, S, Domínguez, L, Álvarez, M, FernándezLGarayzabal, J, Zapata, A & Muñoz, M 2002, ‘Mortality of rainbow trout [Oncorynchus mykiss (Walbaum)] associated with freshwater dinoflagellate bloom [Peridinium polonicum (Woloszynska)] in a fish farm’, Aquaculture Research, vol. 33, no. 2, pp. 141–145.

Ross, J, Beard, J, Wild-Allen, K, Andrewartha, Moreno, D, Semmens, J, Davey, A, Hortle, J, Pender, Quigley B., Andrewartha, J, Stehfest, K, Durand, A & Macleod, Ca 2021, Understanding oxygen dynamics and the importance for benthic recovery in Macquarie Harbour. Hobart, Australia CC BY 3.0, Institute for Marine and Antarctic Studies, University of Tasmania.

Rotterová, J, Edgcomb, VP, Čepička, I & Beinart, R 2022, ‘Anaerobic Ciliates as a Model Group for Studying Symbioses in OxygenLdepleted Environments’, Journal of Eukaryotic Microbiology, p. e12912.

Sanderson, JC, Cromey, CJ, Dring, MJ & Kelly, MS 2008, ‘Distribution of nutrients for seaweed cultivation around salmon cages at farm sites in north–west Scotland’, Aquaculture, vol. 278, no. 1-4, pp. 60–68.

Santoferrara, LF, Alder, VV & McManus, GB 2017, ‘Phylogeny, classification and diversity of Choreotrichia and Oligotrichia (Ciliophora, Spirotrichea)’, Molecular Phylogenetics and Evolution, vol. 112, pp. 12–22.

Sarà, G, Lo Martire, M, Buffa, G, Mannino, AM & Badalamenti, F 2007, ‘The fouling community as an indicator of fish farming impact in Mediterranean’, Aquaculture Research, vol. 38, no. 1, pp. 66–75.

Sehein, TR, Gast, RJ, Pachiadaki, M, Guillou, L & Edgcomb, VP 2022, ‘Parasitic infections by Group II Syndiniales target selected dinoflagellate host populations within diverse protist assemblages in a model coastal pond’, Environ Microbiol, vol. 24, no. 4, pp. 1818–1834.

Shade, A & Handelsman, J 2012, ‘Beyond the Venn diagram: the hunt for a core microbiome’, Environ Microbiol, vol. 14, no. 1, pp. 4–12.

Sherr, EB & Sherr, BF 2002, ‘Significance of predation by protists in aquatic microbial food webs’, Antonie Van Leeuwenhoek, vol. 81, no. 1, pp. 293–308.

Sinnott, RW 1984, ‘Virtues of the Haversine’, S&T, vol. 68, no. 2, p. 158.

Siver, PA 1995, ‘The distribution of chrysophytes along environmental gradients: their use as biological indicators’, Chrysophyte algae, pp. 232–268.

Skogen, MD, Eknes, M, Asplin, LC & Sandvik, AD 2009, ‘Modelling the environmental effects of fish farming in a Norwegian fjord’, Aquaculture, vol. 298, no. 1-2, pp. 70–75.

Skovgaard, A 2014, ‘Dirty tricks in the plankton: diversity and role of marine parasitic protists’, Acta Protozoologica, vol. 53, no. 1.

Song, W 2000, ‘Morphological and taxonomical studies on some marine scuticociliates from China Sea, with description of two new species, Philasterides armatalis sp. n. and Cyclidium varibonneti sp. n.[Protozoa: Ciliophora: Scuticociliatida]’, Acta Protozoologica, vol. 39, no. 4.

Stankus, A 2021, ‘State of world aquaculture 2020 and regional reviews: FAO webinar series’, FAO Aquaculture Newsletter, no. 63, pp. 17–18.

Steele, JA, Countway, PD, Xia, L, Vigil, PD, Beman, JM, Kim, DY, Chow, C-ET, Sachdeva, R, Jones, AC & Schwalbach, MS 2011, ‘Marine bacterial, archaeal and protistan association networks reveal ecological linkages’, The ISME Journal, vol. 5, no. 9, pp. 1414–1425.

Stoeck, T, Bass, D, Nebel, M, Christen, R, Jones, MD, Breiner, HW & Richards, TA 2010, ‘Multiple marker parallel tag environmental DNA sequencing reveals a highly complex eukaryotic community in marine anoxic water’, Mol Ecol, vol. 19 Suppl 1, pp. 21–31.

Stoeck, T, Kochems, R, Forster, D, Lejzerowicz, F & Pawlowski, J 2018, ‘Metabarcoding of benthic ciliate communities shows high potential for environmental monitoring in salmon aquaculture’, Ecological Indicators, vol. 85, pp. 153–164.

Strassert, JF, Hehenberger, E, Del Campo, J, Okamoto, N, Kolisko, M, Richards, TA, Worden, AZ, Santoro, AE & Keeling, PJ 2019, ‘Phylogeny, evidence for a cryptic plastid, and distribution of Chytriodinium parasites (Dinophyceae) infecting copepods’, Journal of Eukaryotic Microbiology, vol. 66, no. 4, pp. 574–581.

Strom, S, Wolfe, G, Holmes, J, Stecher, H, Shimeneck, C & Sarah, L 2003, ‘Chemical defense in the microplankton I: Feeding and growth rates of heterotrophic protists on the DMSLproducing phytoplankter Emiliania huxleyi’, Limnology and Oceanography, vol. 48, no. 1, pp. 217–229.

Strom, SL, Barberi, O, Mazur, C, Bright, K & Fredrickson, K 2020, ‘High light stress reduces dinoflagellate predation on phytoplankton through both direct and indirect responses’, Aquatic Microbial Ecology, vol. 84, pp. 43–57.

Sun, X, Kop, LFM, Lau, MCY, Frank, J, Jayakumar, A, Lucker, S & Ward, BB 2019, ‘Uncultured Nitrospina-like species are major nitrite oxidizing bacteria in oxygen minimum zones’, ISME J, vol. 13, no. 10, pp. 2391–2402.

Teasdale, PR, Apte, SC, Ford, PW, Batley, GE & Koehnken, L 2003, ‘Geochemical cycling and speciation of copper in waters and sediments of Macquarie Harbour, Western Tasmania’, Estuarine, Coastal and Shelf Science, vol. 57, no. 3, pp. 475–487.

Terry, C 2001, ‘Numerical Modelling of Macquarie Harbour, Tasmania’, in In: Conference on Hydraulics in Civil Engineering (6th : 2001 : Hobart, Tas.). 6th Conference on Hydraulics in Civil Engineering: The State of Hydraulics; Proceedings. Barton, A.C.T.: Institution of Engineers, Australia, 2001: 345-354., DOI 10.3316/informit.521499006787667, <https://search.informit.org/doi/10.3316/informit.521499006787667>.

Tičina, V, Katavić, I & Grubišić, L 2020, ‘Marine Aquaculture Impacts on Marine Biota in Oligotrophic Environments of the Mediterranean Sea – A Review’, Frontiers in Marine Science, vol. 7.

Torres-Beltrán, M, Sehein, T, Pachiadaki, MG, Hallam, SJ & Edgcomb, V 2018, ‘Protistan parasites along oxygen gradients in a seasonally anoxic fjord: A network approach to assessing potential host-parasite interactions’, Deep Sea Research Part II: Topical Studies in Oceanography, vol. 156, pp. 97–110.

Tsagaraki, TM, Petihakis, G, Tsiaras, K, Triantafyllou, G, Tsapakis, M, Korres, G, Kakagiannis, G, Frangoulis, C & Karakassis, I 2011, ‘Beyond the cage: ecosystem modelling for impact evaluation in aquaculture’, Ecological modelling, vol. 222, no. 14, pp. 2512–2523.

Tsagaraki, TM, Pitta, P, Frangoulis, C, Petihakis, G & Karakassis, I 2013, ‘Plankton response to nutrient enrichment is maximized at intermediate distances from fish farms’, Marine Ecology Progress Series, vol. 493, pp. 31–42.

Urrutxurtu, I, Orive, E & de la Sota, A 2003, ’Seasonal dynamics of ciliated protozoa and their potential food in an eutrophic estuary (Bay of Biscay)’, Estuarine, Coastal and Shelf Science, vol. 57, no. 5-6, pp. 1169–1182.

Varkey, D, Mazard, S, Jeffries, TC, Hughes, DJ, Seymour, J, Paulsen, IT & Ostrowski, M 2018, ‘Stormwater influences phytoplankton assemblages within the diverse, but impacted Sydney Harbour estuary’, PLOS ONE, vol. 13, no. 12, p. e0209857.

Verhoeven, JTP, Salvo, F, Knight, R, Hamoutene, D & Dufour, SC 2018, ‘Temporal Bacterial Surveillance of Salmon Aquaculture Sites Indicates a Long Lasting Benthic Impact With Minimal Recovery’, Front Microbiol, vol. 9, p. 3054.

Vørs, N 1992, ‘Ultrastructure and autecology of the marine, heterotrophic flagellate Leucocryptos marina (Braarud) Butcher 1967 (Katablepharidaceae/Kathablepharidae), with a discussion of the genera Leucocryptos and Katablepharis/Kathablepharis’, European journal of protistology, vol. 28, no. 4, pp. 369–389.

Wang, Q, Garrity, GM, Tiedje, JM & Cole, JR 2007, ‘Naive Bayesian classifier for rapid assignment of rRNA sequences into the new bacterial taxonomy’, Appl Environ Microbiol, vol. 73, no. 16, pp. 5261–5267.

Wang, Z, Guo, X, Qu, L & Lin, L 2017, ‘Effects of nitrogen and phosphorus on the growth of Levanderina fissa: How it blooms in Pearl River Estuary’, Journal of Ocean University of China, vol. 16, no. 1, pp. 114–120.

Warren, A, Patterson, DJ, Dunthorn, M, Clamp, JC, AchillesLDay, UE, Aescht, E, AlLFarraj, SA, AlLQuraishy, S, AlLRasheid, K & Carr, M 2017, ‘Beyond the “Code”: a guide to the description and documentation of biodiversity in ciliated protists (Alveolata, Ciliophora)’, Journal of Eukaryotic Microbiology, vol. 64, no. 4, pp. 539–554.

Weber, F, Anderson, R, Foissner, W, Mylnikov, AP & Jürgens, K 2014, ‘Morphological and molecular approaches reveal highly stratified protist communities along Baltic Sea pelagic redox gradients’, Aquatic Microbial Ecology, vol. 73, no. 1, pp. 1–16.

Welch, A, Hoenig, R, Stieglitz, J, Benetti, D, Tacon, A, Sims, N & O’Hanlon, B 2010, ‘From Fishing to the Sustainable Farming of Carnivorous Marine Finfish’, Reviews in Fisheries Science, vol. 18, no. 3, pp. 235–247.

Wickham, H, Averick, M, Bryan, J, Chang, W, McGowan, L, François, R, Grolemund, G, Hayes, A, Henry, L, Hester, J, Kuhn, M, Pedersen, T, Miller, E, Bache, S, Müller, K, Ooms, J, Robinson, D, Seidel, D, Spinu, V, Takahashi, K, Vaughan, D, Wilke, C, Woo, K & Yutani, H 2019, ‘Welcome to the Tidyverse’, Journal of Open Source Software, vol. 4, no. 43, p. 1686.

Williams, CR, Baccarella, A, Parrish, JZ & Kim, CC 2017, ‘Empirical assessment of analysis workflows for differential expression analysis of human samples using RNA-Seq’, BMC Bioinformatics, vol. 18, no. 1, pp. 1–12.

Worden, AZ, Follows, MJ, Giovannoni, SJ, Wilken, S, Zimmerman, AE & Keeling, PJ 2015, ‘Rethinking the marine carbon cycle: factoring in the multifarious lifestyles of microbes’, Science, vol. 347, no. 6223, p. 1257594.

Wu, F, Dai, M, Huang, H & Qi, Z 2018, ‘Plankton ciliate community responses to different aquatic environments in Nan’ao Island, a representative mariculture base in the South China Sea’, Marine and Freshwater Research, vol. 70, no. 3, pp. 426–436.

Xiaoshan, Z, Bin, Y, Yanhong, D & Lianfeng, Y 2004, ‘A primary study on one of “bilateral” red tide at Chi Bay of Pearl River Estuary’, Marine Environmental Science, vol. 23, no. 4, pp. 41–44.

Xu, H, Min, G-S, Choi, J-K, Al-Rasheid, KA, Lin, X & Zhu, M 2010, ‘Temporal dynamics of phytoplankton communities in a semi-enclosed mariculture pond and their responses to environmental factors’, Chinese Journal of Oceanology and Limnology, vol. 28, no. 2, pp. 295–303.

Yang, J, Huang, S, Fan, W, Warren, A, Jiao, N & Xu, D 2020, ‘Spatial distribution patterns of planktonic ciliate communities in the East China Sea: potential indicators of water masses’, Marine Pollution Bulletin, vol. 156, p. 111253.

Zhang, H, Huang, X, Huang, L, Bao, F, Xiong, S, Wang, K & Zhang, D 2018, ‘Microeukaryotic biogeography in the typical subtropical coastal waters with multiple environmental gradients’, Sci Total Environ, vol. 635, pp. 618–628.

Zhang, Q, Tang, F, Zhou, Y, Xu, J, Chen, H, Wang, M & Laanbroek, HJ 2015, ‘Shifts in the pelagic ammonia-oxidizing microbial communities along the eutrophic estuary of Yong River in Ningbo City, China’, Front Microbiol, vol. 6, p. 1180.

