## Supplementary material for "Effects of finfish farms on pelagic protist communities in a semi-closed stratified embayment": sup_material_figures

### Supplementary Material - Figures

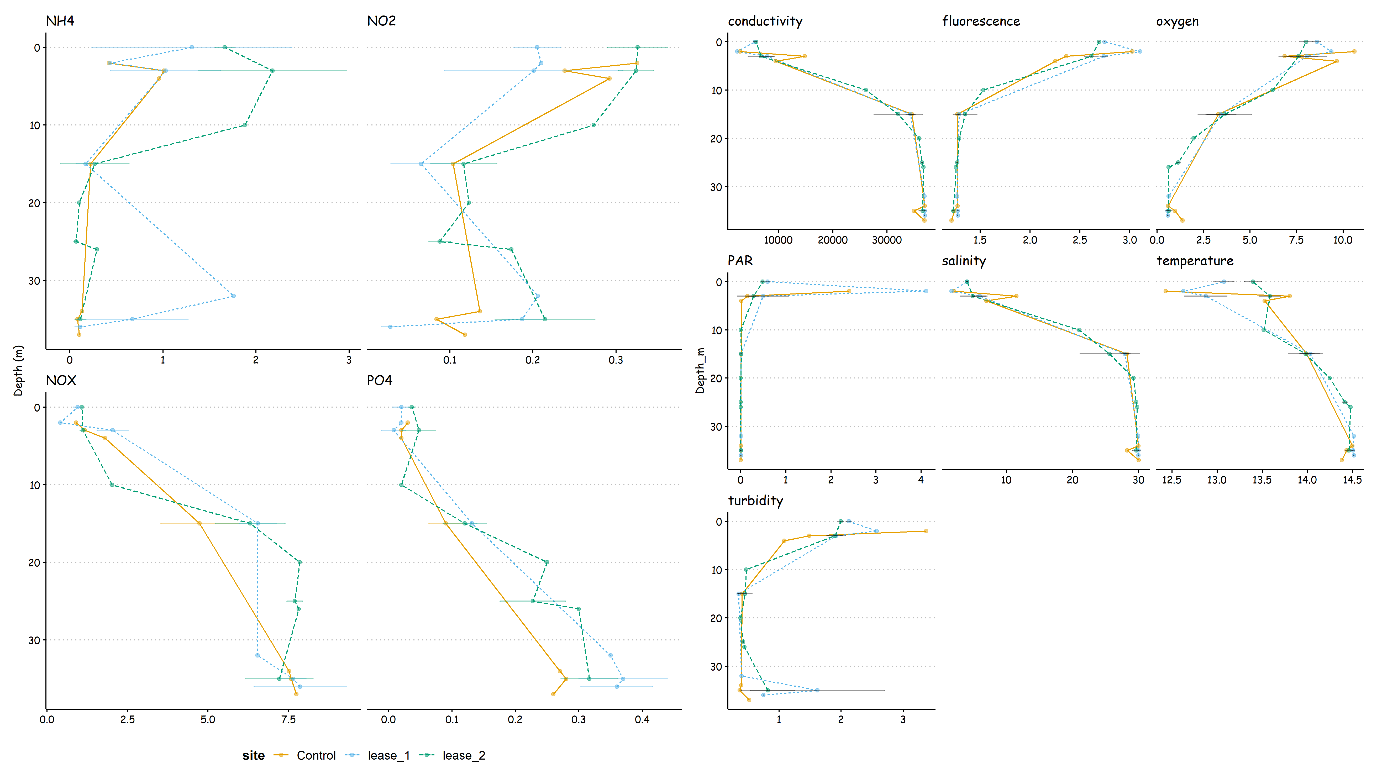

Fig. S1 – Depth profile of the environmental variables. Colors represent samples sites (both leases and control sites).

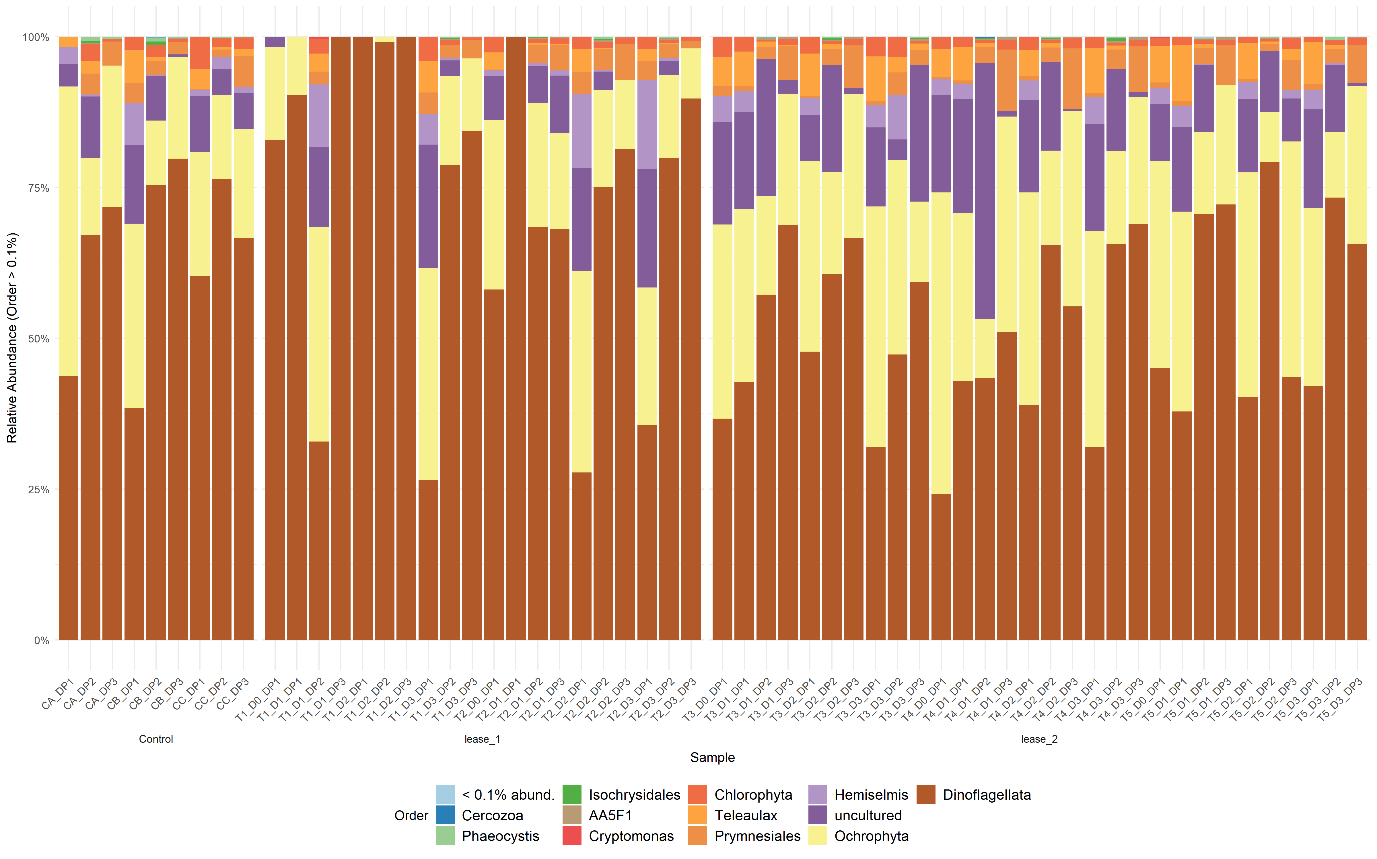

Fig. S2 - Bar plot showing the relative abundance distribution of phytoplankton community across sites. ASVs with abundance less than 0.1% were agglomerate together.

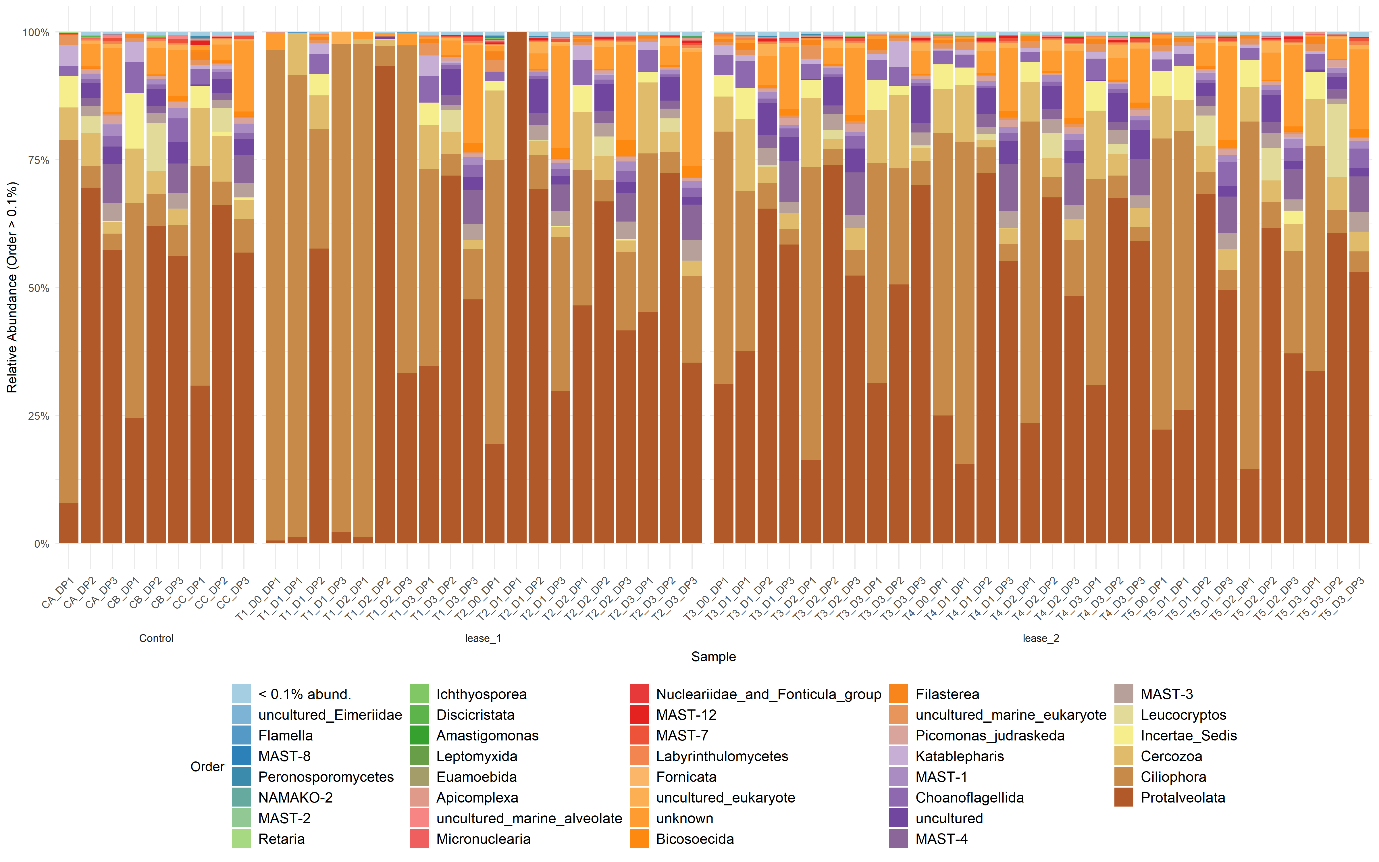

Fig. S3 - Bar plot showing the relative abundance distribution of protists (non-phytoplankton) community across sites. ASVs with abundance less than 0.1% were agglomerate together.

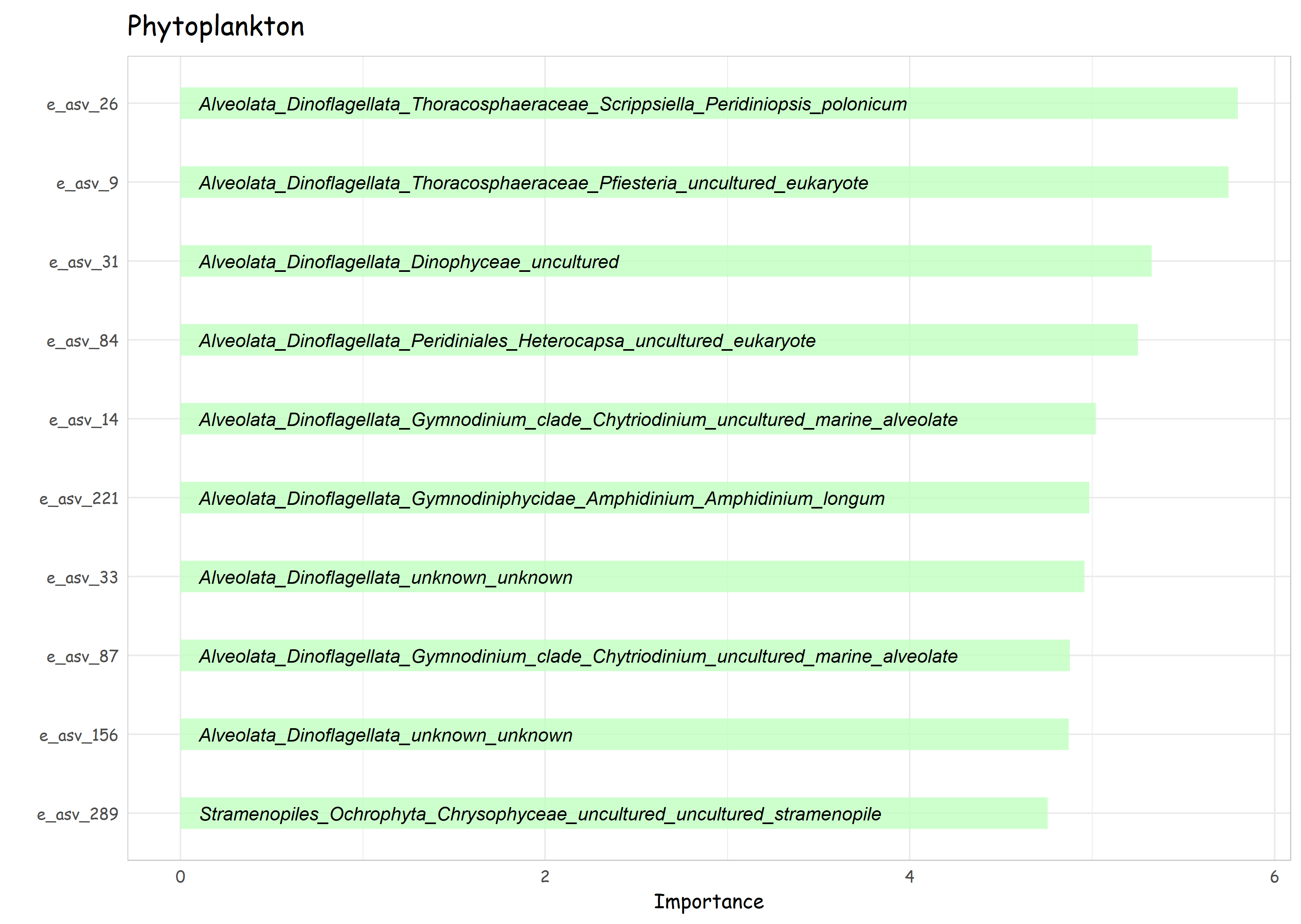

Fig. S4 - The 10 most important phytoplankton ASVs based on mean decrease in Gini index in classification-based prediction model of the lease and control sites. ASV identifications are displayed in y-axis along with the respective taxonomic levels inside of each bar.

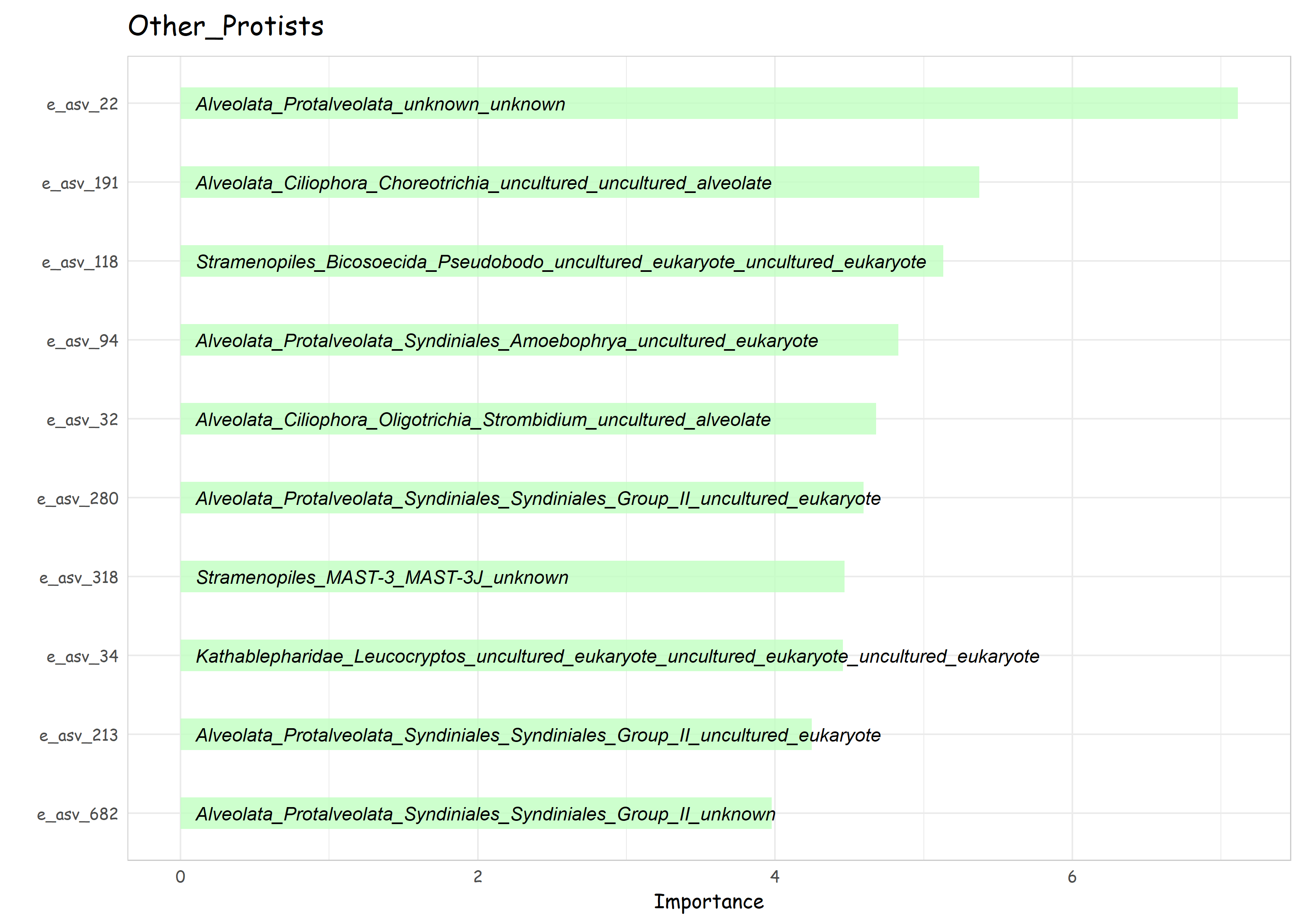

Fig. S5 - The 10 most important protists ASVs (non-phytoplankton) based on mean decrease in Gini index in classification-based prediction model of the lease and control sites. ASV identifications are displayed in y-axis along with the respective taxonomic levels inside of each bar.

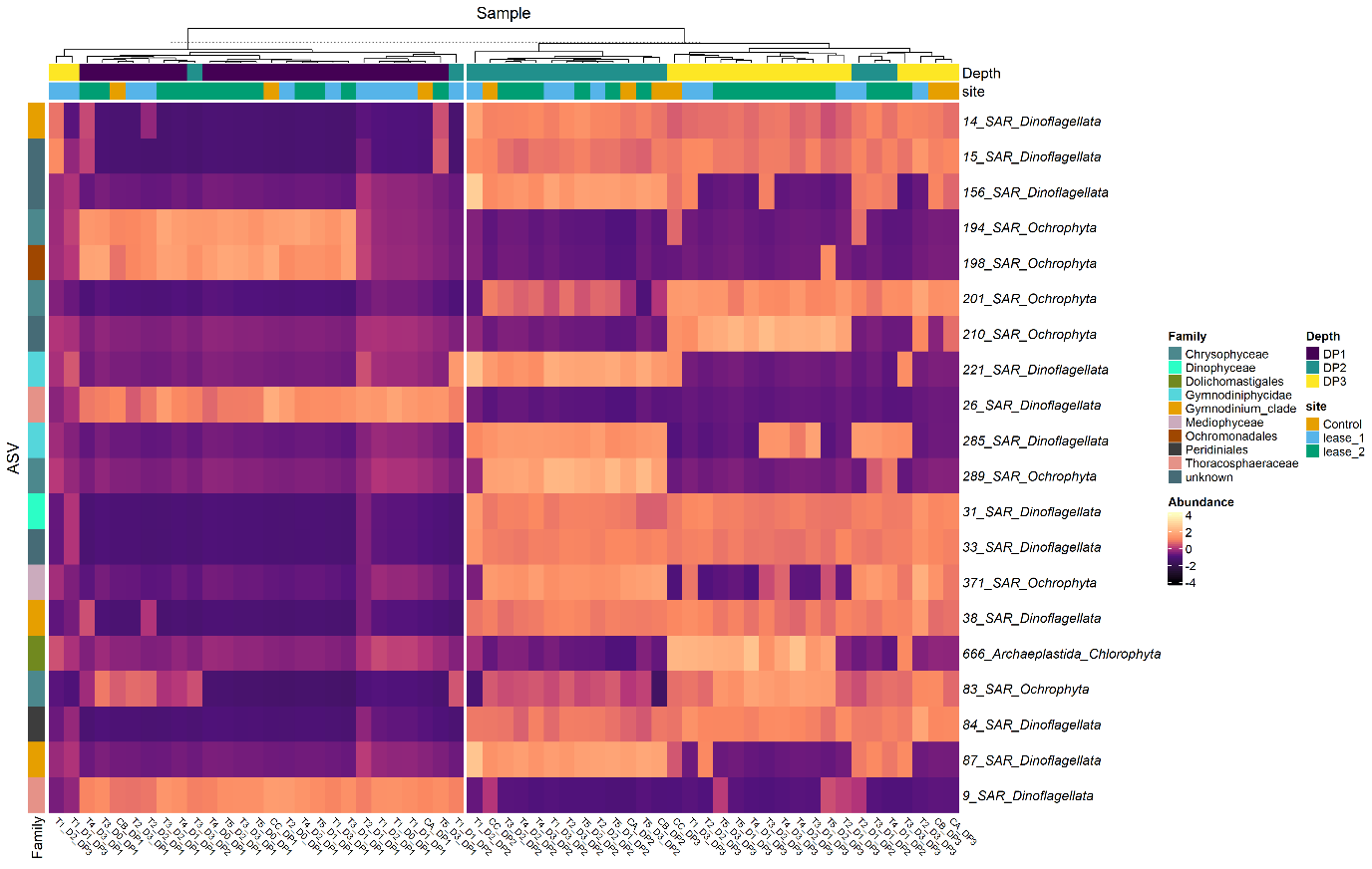

Fig. S6- Heatmap displaying the relative abundance of the top 20 most important phytoplankton ASVs found by the random forest model classifying water layers. The color code indicates the centered log-ratio (clr) relative abundance for a given ASV.

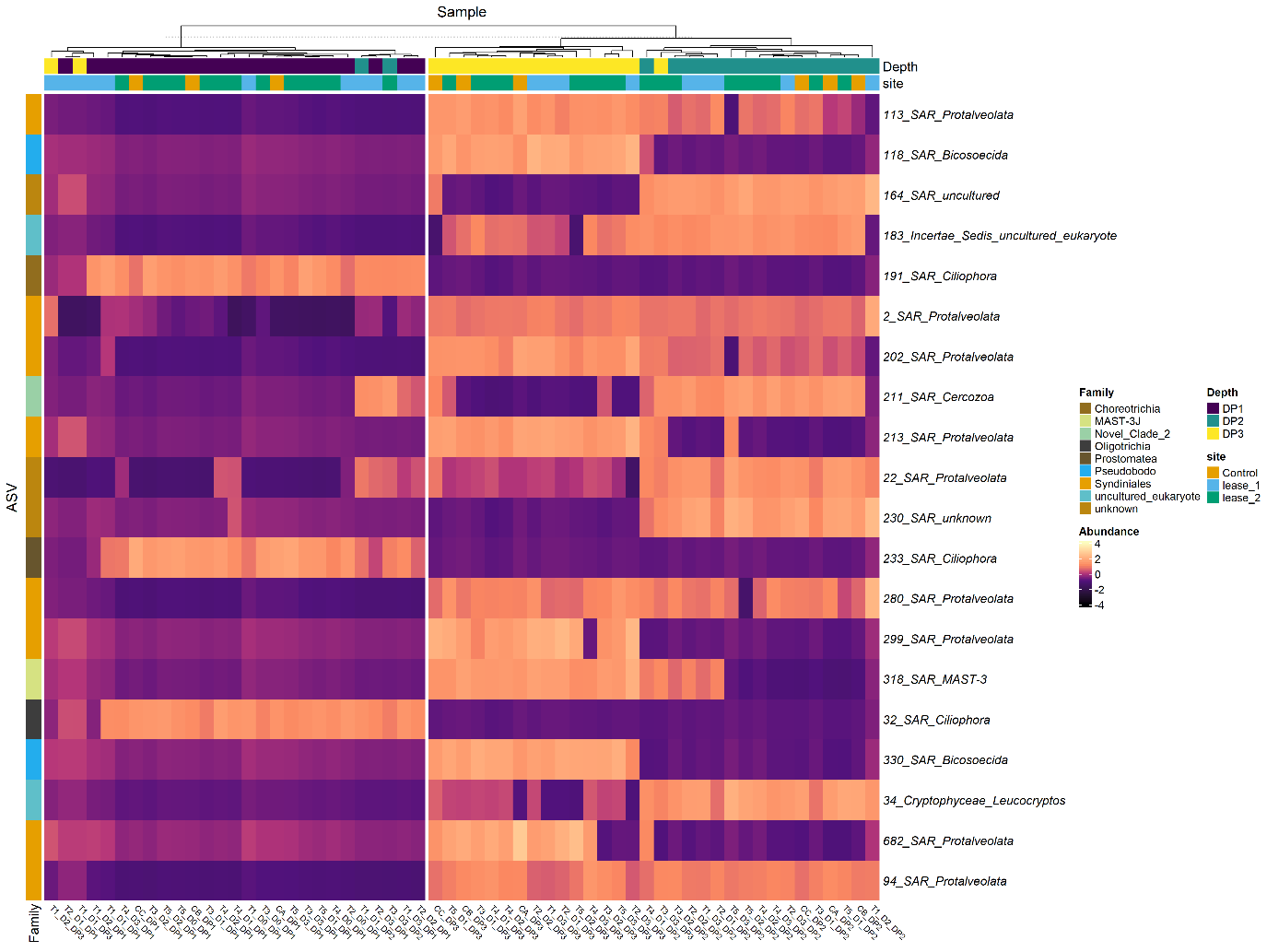

Fig. S7 - Heatmap displaying the relative abundance of the protists ASVs (non-phytoplankton) found by both differential abundant and random forest classification (top 40 vips) approaches comparing water layers. The color code indicates the centered log-ratio (clr) relative abundance for a given ASV.

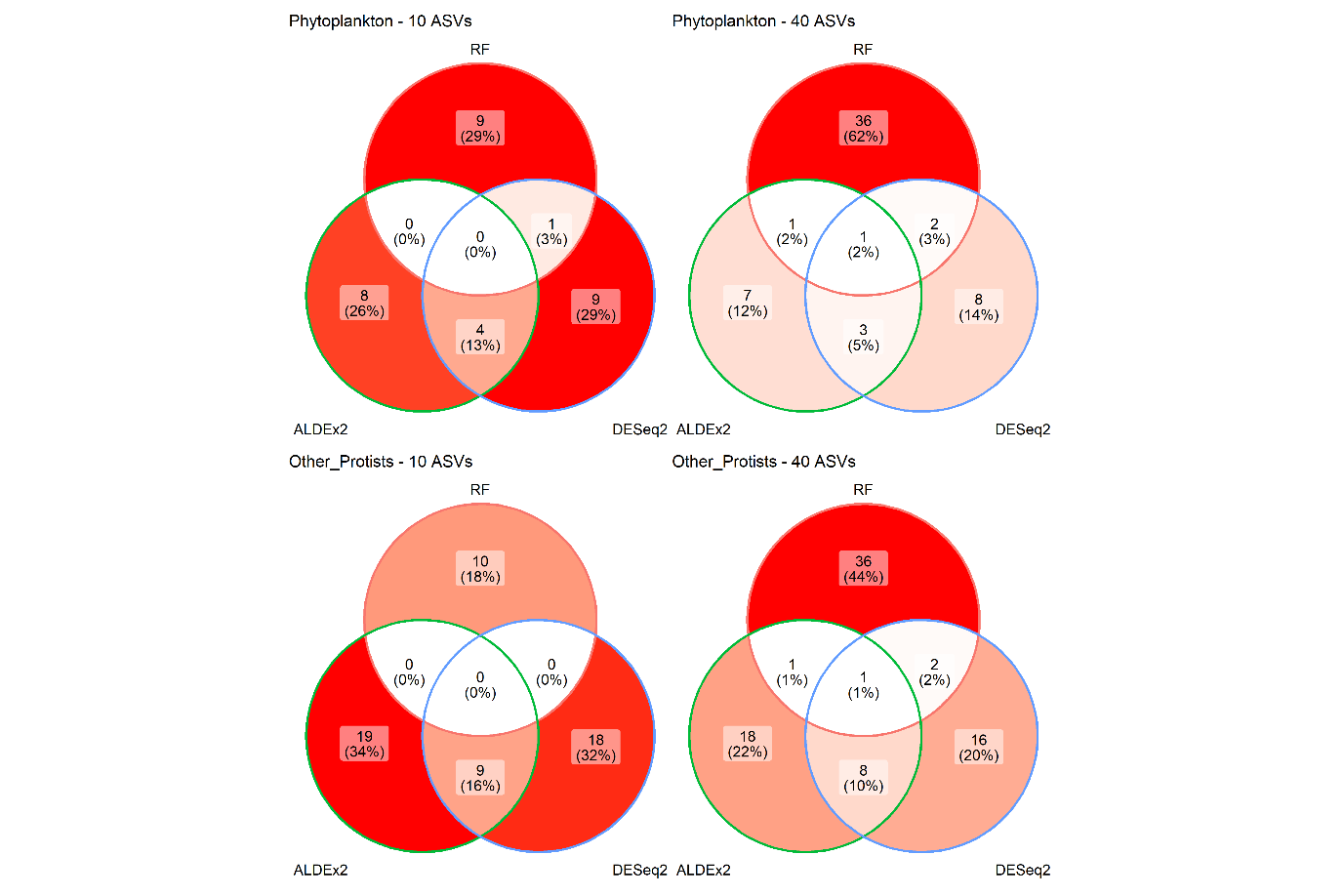

Fig. S8 – Venn diagram displaying the number of shared ASVs found among the three feature selection methods used in this study (ALDEx2, DESEq2 and random forest classification model). We compared the top 10 (left-hand site) and top 40 (right-hand side) most important predictive ASVs according to random forest with the ASVs found in the two differential abundance approaches.

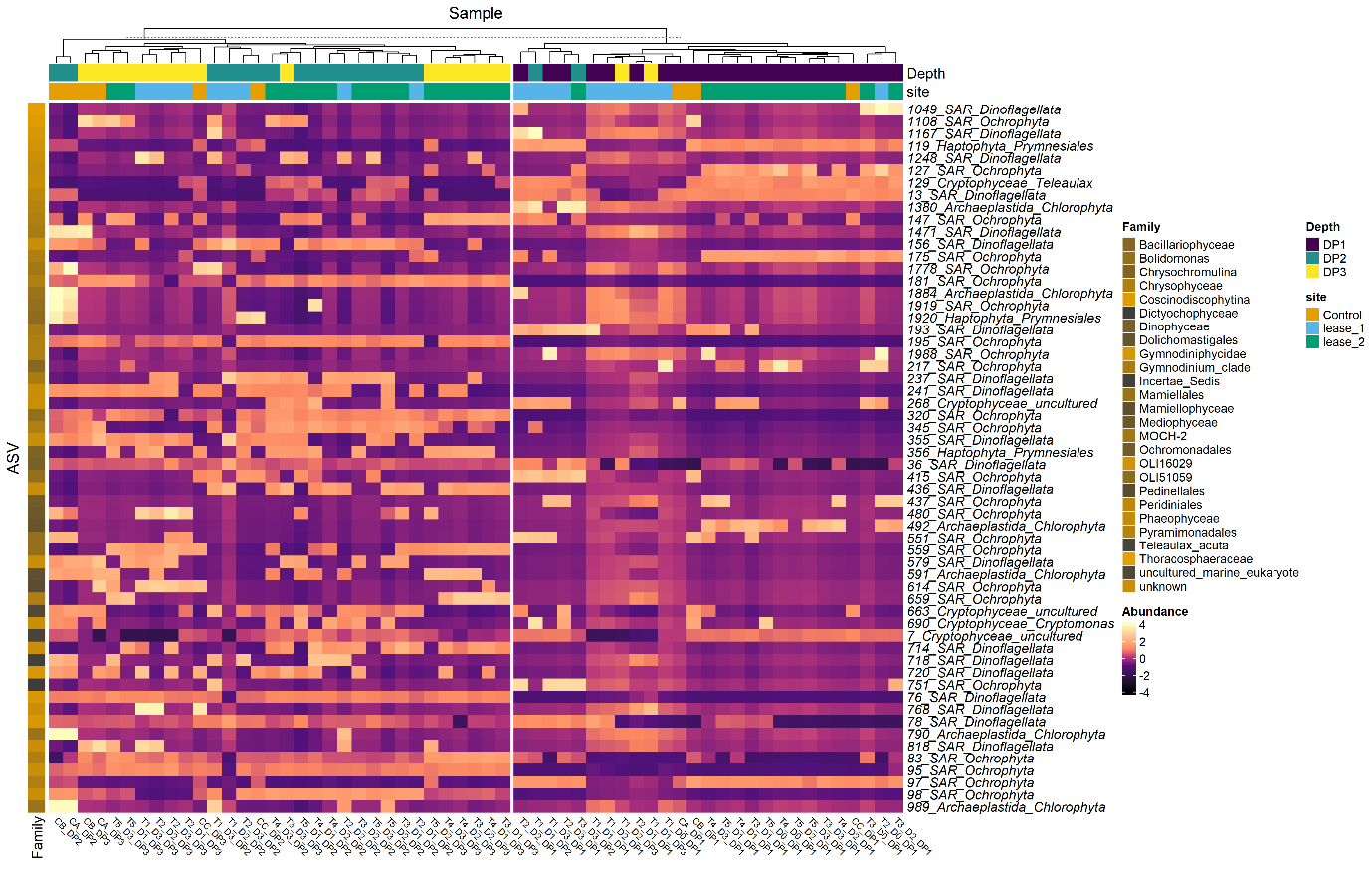

Fig. S9 – Heatmap displaying the relative abundance of the phytoplanktonic ASVs found by both differential abundant and random forest classification (top 40 vips) approaches comparing lease and control sites. The color code indicates the centered log-ratio (clr) relative abundance for a given ASV.

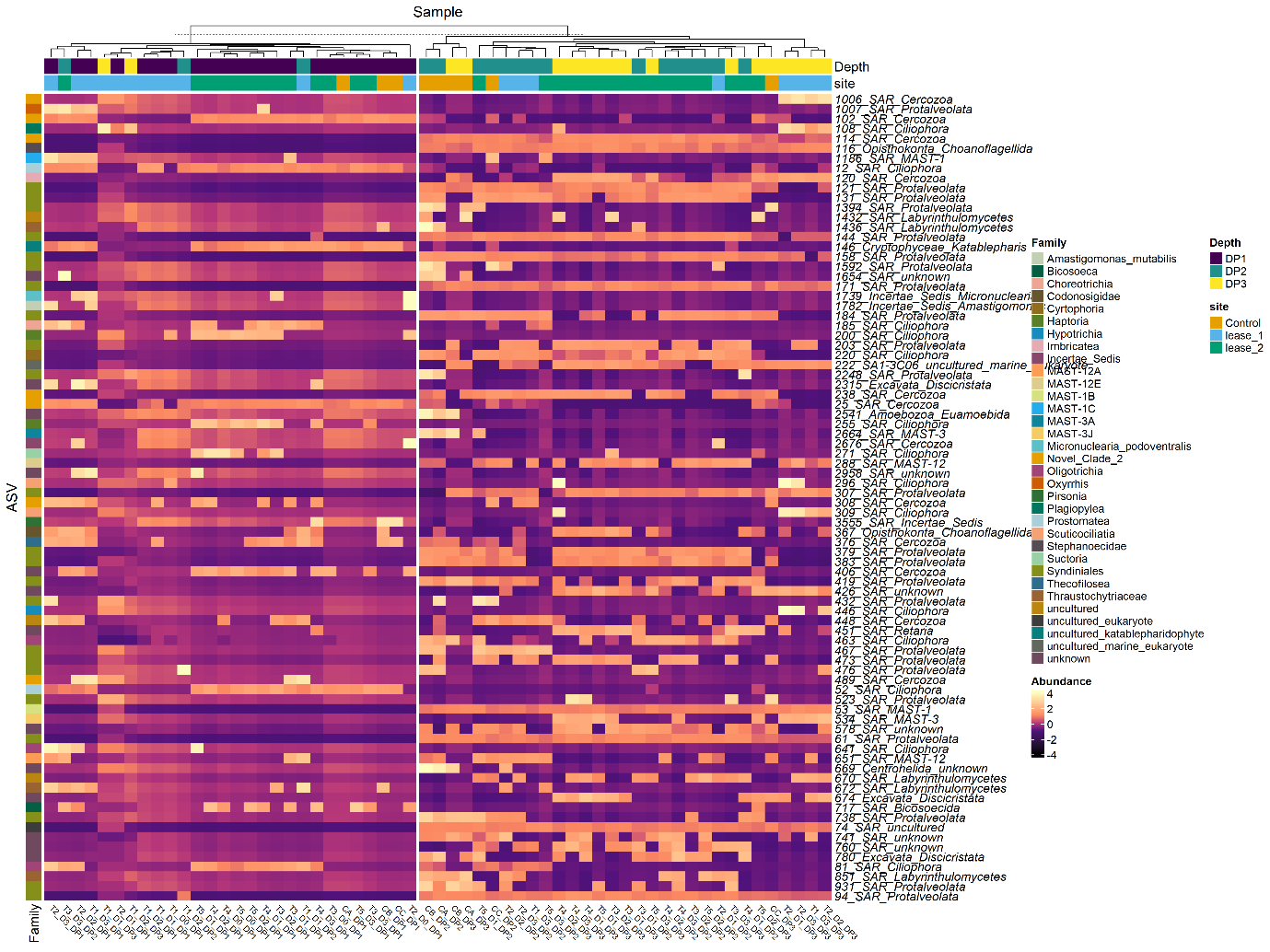

Fig. S10 - Heatmap displaying the relative abundance of the protists ASVs (non-phytoplankton) found by both differential abundant and random forest classification (top 40 vips) approaches comparing lease and control sites. The color code indicates the centered log-ratio (clr) relative abundance for a given ASV.
