## Supplementary material for "Effects of finfish farms on pelagic protist communities in a semi-closed stratified embayment": sup_material_tables

### Supplementary Material - Tables

Table S1 - Environmental variables measurements.

| Sample id | site | Dist(m) | Depth(m) | lat | long | conductivity | oxygen | salinity | temperature | PAR | fluorescence | turbidity | NH4 | NOX | NO2 | PO4 |
| --- | --- | --- | --- | --- | --- | --- | --- | --- | --- | --- | --- | --- | --- | --- | --- | --- |
| T1_D0_DP1 | lease_1 | 0 | 0 | -42.3315 | 145.3922 | 5862.10 | 8.45 | 4.08 | 12.99 | 0.57 | 2.75 | 2.10 | 0.55 | 1.05 | 0.185 | 0.03 |
| T1_D1_DP1 | lease_1 | 16.5 | 3 | -42.3315 | 145.3924 | 6185.44 | 9.72 | 4.57 | 12.50 | 0.46 | 2.77 | 1.84 | 0.54 | 2.35 | 0.116 | -0.02 |
| T1_D1_DP2 | lease_1 | 16.5 | 15 | -42.3315 | 145.3924 | 34352.33 | 3.64 | 27.98 | 14.05 | 0.00 | 1.29 | 0.34 | 0.08 | 7.25 | 0.007 | 0.13 |
| T1_D1_DP3 | lease_1 | 16.5 | 36 | -42.3315 | 145.3924 | 36953.40 | 0.59 | 29.98 | 14.51 | 0.00 | 1.27 | 0.75 | 0.07 | 8.88 | 0.021 | 0.4 |
| T1_D2_DP1 | lease_1 | 18.2 | 3 | -42.3314 | 145.3924 | 9411.00 | 7.41 | 7.29 | 12.85 | 1.41 | 2.76 | 1.90 | 0.65 | 2.73 | 0.058 | 0.03 |
| T1_D2_DP2 | lease_1 | 18.2 | 15 | -42.3314 | 145.3924 | 34550.25 | 3.29 | 28.13 | 14.10 | 0.00 | 1.29 | 0.35 | 0.26 | 6.42 | 0.035 | 0.14 |
| T1_D2_DP3 | lease_1 | 18.2 | 36 | -42.3314 | 145.3924 | 36956.00 | 0.56 | 29.99 | 14.51 | 0.00 | 1.28 | 0.75 | 0.15 | 6.85 | 0.036 | 0.32 |
| T1_D3_DP1 | lease_1 | 82.6 | 3 | -42.3315 | 145.3932 | 7485.40 | 7.10 | 5.66 | 12.90 | 0.02 | 2.74 | 1.90 | 0.61 | 1.54 | 0.302 | 0 |
| T1_D3_DP2 | lease_1 | 82.6 | 15 | -42.3315 | 145.3932 | 35030.75 | 3.19 | 28.54 | 14.14 | 0.00 | 1.29 | 0.35 | 0.18 | 6.62 | 0.097 | 0.13 |
| T1_D3_DP3 | lease_1 | 82.6 | 35 | -42.3315 | 145.3932 | 36935.33 | 0.66 | 29.97 | 14.51 | 0.00 | 1.27 | 0.43 | 0.11 | 7.99 | 0.163 | 0.3 |
| T2_D0_DP1 | lease_1 | 0 | 0 | -42.3319 | 145.3916 | 5455.67 | 8.74 | 3.94 | 13.15 | 0.62 | 2.75 | 2.15 | 2.07 | 0.85 | 0.225 | 0.01 |
| T2_D1_DP1 | lease_1 | 23.5 | 2 | -42.3321 | 145.3918 | 2434.73 | 9.36 | 1.70 | 12.62 | 4.11 | 3.11 | 2.58 | 0.43 | 0.41 | 0.21 | 0.02 |
| T2_D1_DP2 | lease_1 | 23.5 | 15 | -42.3321 | 145.3918 | 32751.33 | 4.90 | 26.70 | 13.83 | 0.00 | 1.31 | 0.36 | 0.09 | 5.52 | 0.077 | 0.14 |
| T2_D1_DP3 | lease_1 | 23.5 | 32 | -42.3321 | 145.3918 | 36889.00 | 0.63 | 29.93 | 14.51 | 0.00 | 1.27 | 0.40 | 1.76 | 6.54 | 0.206 | 0.35 |
| T2_D2_DP1 | lease_1 | 28.5 | 3 | -42.3318 | 145.3919 | 7128.50 | 8.46 | 5.32 | 13.17 | 0.57 | 2.76 | 2.18 | 1.8 | 1.75 | 0.274 | 0.02 |
| T2_D2_DP2 | lease_1 | 28.5 | 15 | -42.3318 | 145.3919 | 34202.00 | 3.14 | 27.89 | 14.00 | 0.00 | 1.28 | 0.34 | 0.09 | 6.78 | 0.097 | 0.13 |
| T2_D2_DP3 | lease_1 | 28.5 | 35 | -42.3318 | 145.3919 | 36967.40 | 0.52 | 29.99 | 14.51 | 0.00 | 1.28 | 2.53 | 0.59 | 7.76 | 0.212 | 0.37 |
| T2_D3_DP1 | lease_1 | 82.8 | 3 | -42.332 | 145.3926 | 9358.57 | 7.63 | 7.06 | 12.95 | 0.01 | 2.67 | 1.78 | 1.54 | 1.83 | 0.254 | 0.01 |
| T2_D3_DP2 | lease_1 | 82.8 | 15 | -42.332 | 145.3926 | 34745.50 | 2.44 | 28.33 | 14.07 | 0.00 | 1.28 | 0.36 | 0.32 | 6.71 | 0.08 | 0.12 |
| T2_D3_DP3 | lease_1 | 82.8 | 35 | -42.332 | 145.3926 | 36935.50 | 0.51 | 29.99 | 14.49 | 0.00 | 1.27 | 1.90 | 1.31 | 7.19 | 0.186 | 0.44 |
| T3_D0_DP1 | lease_2 | 0 | 0 | -42.2912 | 145.3474 | 5904.89 | 8.41 | 4.12 | 13.28 | 0.49 | 2.70 | 2.05 | 1.61 | 1.04 | 0.294 | 0.03 |
| T3_D1_DP1 | lease_2 | 6.63 | 3 | -42.2913 | 145.3475 | 5790.11 | 7.76 | 3.95 | 13.62 | 0.28 | 2.64 | 1.91 | 3 | 1.16 | 0.302 | 0.04 |
| T3_D1_DP2 | lease_2 | 6.63 | 15 | -42.2913 | 145.3475 | 24159.79 | 5.35 | 17.91 | 13.69 | 0.05 | 1.57 | 0.67 | 0.56 | 6.02 | 0.083 | 0.14 |
| T3_D1_DP3 | lease_2 | 6.63 | 25 | -42.2913 | 145.3475 | 36287.82 | 1.13 | 29.58 | 14.42 | 0.00 | 1.27 | 0.42 | 0.07 | 7.88 | 0.089 | 0.25 |
| T3_D2_DP1 | lease_2 | 13.26 | 3 | -42.2913 | 145.3475 | 7205.33 | 7.48 | 5.33 | 13.44 | 0.27 | 2.63 | 1.90 | 1.54 | 1.05 | 0.344 | 0.02 |
| T3_D2_DP2 | lease_2 | 13.26 | 15 | -42.2913 | 145.3475 | 25555.04 | 4.97 | 18.67 | 13.69 | 0.03 | 1.51 | 0.59 | 0.05 | 6.87 | 0.072 | 0.12 |
| T3_D2_DP3 | lease_2 | 13.26 | 25 | -42.2913 | 145.3475 | 36436.19 | 1.05 | 29.60 | 14.42 | 0.00 | 1.26 | 0.42 | 0.06 | 7.86 | 0.076 | 0.26 |
| T3_D3_DP1 | lease_2 | 75 | 3 | -42.2916 | 145.3481 | 7114.97 | 7.38 | 5.25 | 13.38 | 0.34 | 2.59 | 1.92 | 0.86 | 1.03 | 0.32 | 0.03 |
| T3_D3_DP2 | lease_2 | 75 | 10 | -42.2916 | 145.3481 | 26112.00 | 6.21 | 21.00 | 13.52 | 0.00 | 1.53 | 0.47 | 1.88 | 2.02 | 0.273 | 0.02 |
| T3_D3_DP3 | lease_2 | 75 | 25 | -42.2916 | 145.3481 | 36400.86 | 1.28 | 29.60 | 14.37 | 0.00 | 1.28 | 0.40 | 0.04 | 7.35 | 0.081 | 0.15 |
| T4_D0_DP1 | lease_2 | 0 | 0 | -42.2909 | 145.3481 | 5859.57 | 7.87 | 3.98 | 13.44 | 0.51 | 2.69 | 1.97 | 1.78 | 1.09 | 0.318 | 0.04 |
| T4_D1_DP1 | lease_2 | 5.7 | 3 | -42.291 | 145.3481 | 4351.44 | 9.20 | 3.02 | 13.72 | 0.23 | 2.71 | 2.02 | 3.2 | 1.17 | 0.343 | 0.1 |
| T4_D1_DP2 | lease_2 | 5.7 | 15 | -42.291 | 145.3481 | 34323.75 | 3.98 | 28.02 | 13.98 | 0.00 | 1.28 | 0.42 | 0.06 | 7.47 | 0.101 | 0.17 |
| T4_D1_DP3 | lease_2 | 5.7 | 20 | -42.291 | 145.3481 | 35891.00 | 1.96 | 29.23 | 14.25 | 0.00 | 1.29 | 0.38 | 0.1 | 7.85 | 0.123 | 0.25 |
| T4_D2_DP1 | lease_2 | 7.77 | 3 | -42.291 | 145.3481 | 4710.39 | 7.88 | 3.29 | 13.62 | 0.42 | 2.72 | 1.82 | 3.1 | 1.06 | 0.304 | 0.08 |
| T4_D2_DP2 | lease_2 | 7.77 | 15 | -42.291 | 145.3481 | 35176.71 | 1.65 | 28.67 | 14.14 | 0.00 | 1.29 | 0.48 | 0.08 | 6.91 | 0.104 | 0.13 |
| T4_D2_DP3 | lease_2 | 7.77 | 26 | -42.291 | 145.3481 | 36677.40 | 0.63 | 29.77 | 14.47 | 0.00 | 1.26 | 0.44 | 0.29 | 7.82 | 0.174 | 0.3 |
| T4_D3_DP1 | lease_2 | 80.54 | 3 | -42.2914 | 145.3486 | 12194.40 | 5.79 | 9.50 | 13.51 | 0.36 | 2.56 | 1.77 | 1.9 | 1.24 | 0.335 | 0.05 |
| T4_D3_DP2 | lease_2 | 80.54 | 15 | -42.2914 | 145.3486 | 35174.50 | 3.10 | 28.65 | 14.15 | 0.00 | 1.28 | 0.38 | 0.06 | 6.94 | 0.102 | 0.13 |
| T4_D3_DP3 | lease_2 | 80.54 | 25 | -42.2914 | 145.3486 | 36568.33 | 1.15 | 29.69 | 14.45 | 0.00 | 1.27 | 0.45 | 0.1 | 7.7 | 0.106 | 0.25 |
| T5_D0_DP1 | lease_2 | 0 | 0 | -42.2891 | 145.3517 | 5932.86 | 7.76 | 4.08 | 13.46 | 0.46 | 2.69 | 1.96 | 1.6 | 1.11 | 0.365 | 0.04 |
| T5_D1_DP1 | lease_2 | 9.47 | 3 | -42.2891 | 145.3518 | 5555.19 | 7.19 | 3.94 | 13.77 | 0.02 | 2.62 | 1.89 | 2.31 | 1.19 | 0.35 | 0.04 |
| T5_D1_DP2 | lease_2 | 9.47 | 15 | -42.2891 | 145.3518 | 33720.00 | 1.47 | 27.34 | 14.19 | 0.00 | 1.28 | 0.37 | 1.08 | 4.04 | 0.162 | 0.07 |
| T5_D1_DP3 | lease_2 | 9.47 | 35 | -42.2891 | 145.3518 | 35801.40 | 0.45 | 28.97 | 14.49 | 0.00 | 1.24 | 0.61 | 0.14 | 6.01 | 0.254 | 0.27 |
| T5_D2_DP1 | lease_2 | 16.13 | 3 | -42.2891 | 145.3518 | 8038.60 | 6.69 | 5.99 | 13.56 | 0.33 | 2.55 | 1.76 | 1.78 | 1.14 | 0.327 | 0.03 |
| T5_D2_DP2 | lease_2 | 16.13 | 15 | -42.2891 | 145.3518 | 34273.50 | 4.40 | 27.94 | 14.02 | 0.00 | 1.29 | 0.36 | 0.07 | 6.64 | 0.119 | 0.13 |
| T5_D2_DP3 | lease_2 | 16.13 | 35 | -42.2891 | 145.3518 | 36860.00 | 0.68 | 29.93 | 14.47 | 0.00 | 1.24 | 0.53 | 0.16 | 7.67 | 0.244 | 0.36 |
| T5_D3_DP1 | lease_2 | 84.25 | 3 | -42.2897 | 145.3523 | 5802.93 | 9.04 | 4.13 | 13.63 | 0.32 | 2.51 | 2.14 | 1.91 | 0.97 | 0.289 | 0.04 |
| T5_D3_DP2 | lease_2 | 84.25 | 15 | -42.2897 | 145.3523 | 34086.00 | 4.28 | 27.81 | 13.96 | 0.00 | 1.28 | 0.34 | 0.18 | 5.63 | 0.19 | 0.07 |
| T5_D3_DP3 | lease_2 | 84.25 | 35 | -42.2897 | 145.3523 | 36972.75 | 0.79 | 30.08 | 14.41 | 0.00 | 1.22 | 1.32 | 0.04 | 7.97 | 0.145 | 0.32 |
| CA_DP1 | Control | >1500 | 3 | -42.2855 | 145.369 | 14858.70 | 6.86 | 11.51 | 13.80 | 0.15 | 2.36 | 1.49 | 1.01 | 1.17 | 0.238 | 0.02 |
| CA_DP2 | Control | >1500 | 15 | -42.2855 | 145.369 | 34385.00 | 3.39 | 28.12 | 13.91 | 0.00 | 1.27 | 0.40 | 0.25 | 4.09 | 0.102 | 0.07 |
| CA_DP3 | Control | >1500 | 35 | -42.2855 | 145.369 | 34954.67 | 0.97 | 28.26 | 14.43 | 0.00 | 1.24 | 0.37 | 0.08 | 7.58 | 0.084 | 0.28 |
| CB_DP1 | Control | >1500 | 4 | -42.2418 | 145.3249 | 9477.53 | 9.67 | 6.98 | 13.52 | 0.01 | 2.25 | 1.09 | 0.96 | 1.8 | 0.292 | 0.02 |
| CB_DP2 | Control | >1500 | 15 | -42.2418 | 145.3249 | 35303.67 | 2.64 | 28.83 | 14.08 | 0.00 | 1.25 | 0.48 | 0.22 | 4 | 0.106 | 0.08 |
| CB_DP3 | Control | >1500 | 37 | -42.2418 | 145.3249 | 36900.75 | 1.37 | 30.04 | 14.38 | 0.00 | 1.21 | 0.52 | 0.1 | 7.75 | 0.118 | 0.26 |
| CC_DP1 | Control | >1500 | 2 | -42.327 | 145.409 | 2920.53 | 10.61 | 2.06 | 12.43 | 2.41 | 3.03 | 3.37 | 0.42 | 0.9 | 0.325 | 0.03 |
| CC_DP2 | Control | >1500 | 15 | -42.327 | 145.409 | 34060.40 | 3.84 | 27.78 | 13.98 | 0.00 | 1.28 | 0.34 | 0.2 | 6.13 | 0.104 | 0.12 |
| CC_DP3 | Control | >1500 | 34 | -42.327 | 145.409 | 36931.83 | 0.59 | 29.98 | 14.50 | 0.00 | 1.27 | 0.39 | 0.13 | 7.52 | 0.136 | 0.27 |

Table S2 - Diversity indices across all samples of both eukaryotic communities analyzed in this study.

|  | Phytoplankton | | | | | Other Protists | | | | |
| --- | --- | --- | --- | --- | --- | --- | --- | --- | --- | --- |
| S**ample** **id** | **Observed** | **Chao1** | **se.chao1** | **Shannon** | **InvSimpson** | **Observed** | **Chao1** | **se.chao1** | **Shannon** | **InvSimpson** |
| T1_D1_DP1 | 10 | 10 | 0.000 | 1.844 | 5.328 | 22 | 22 | 0.000 | 1.702 | 3.120 |
| T2_D2_DP2 | 144 | 144 | 0.249 | 3.752 | 19.432 | 271 | 271 | 0.000 | 3.956 | 19.295 |
| CA_DP2 | 163 | 163 | 0.000 | 3.906 | 24.876 | 286 | 286 | 0.000 | 4.003 | 21.388 |
| T2_D0_DP1 | 85 | 86 | 2.335 | 2.653 | 5.647 | 100 | 100 | 0.000 | 2.616 | 5.561 |
| T4_D1_DP2 | 138 | 138 | 0.000 | 3.000 | 6.142 | 287 | 287 | 0.166 | 3.816 | 16.830 |
| T1_D1_DP3 | 1 | 1 | 0.000 | 0.000 | 1.000 | 2 | 2 | 0.000 | 0.111 | 1.048 |
| T1_D3_DP2 | 116 | 116 | 0.000 | 3.693 | 21.316 | 248 | 248 | 0.083 | 3.961 | 26.509 |
| T2_D3_DP2 | 138 | 139 | 1.297 | 3.498 | 13.877 | 262 | 262 | 0.100 | 4.020 | 24.094 |
| CC_DP2 | 139 | 139 | 0.000 | 3.592 | 14.767 | 274 | 274 | 0.000 | 4.194 | 31.364 |
| T2_D3_DP3 | 85 | 85 | 0.000 | 2.993 | 11.734 | 196 | 196 | 0.499 | 3.695 | 14.886 |
| T2_D2_DP1 | 98 | 101 | 4.642 | 3.467 | 17.328 | 137 | 137 | 0.000 | 2.927 | 6.753 |
| T1_D2_DP1 | 5 | 5 | 0.000 | 0.940 | 1.991 | 8 | 8 | 0.000 | 0.833 | 1.637 |
| T4_D0_DP1 | 59 | 59 | 0.000 | 3.132 | 13.410 | 85 | 85 | 0.000 | 2.980 | 8.342 |
| T4_D1_DP3 | 119 | 119 | 0.000 | 3.810 | 25.886 | 278 | 278 | 0.000 | 3.820 | 18.837 |
| T3_D0_DP1 | 48 | 48 | 0.000 | 3.161 | 12.576 | 44 | 44 | 0.000 | 2.638 | 7.078 |
| T2_D3_DP1 | 80 | 80 | 0.000 | 3.098 | 10.859 | 122 | 122 | 0.000 | 3.067 | 9.959 |
| T1_D0_DP1 | 9 | 9 | 0.000 | 1.635 | 3.889 | 14 | 14 | 0.000 | 0.914 | 1.565 |
| CC_DP1 | 72 | 72 | 0.000 | 2.640 | 8.152 | 94 | 94 | 0.000 | 2.970 | 8.626 |
| T5_D3_DP3 | 107 | 107 | 0.249 | 3.747 | 26.243 | 233 | 233 | 0.000 | 3.470 | 10.666 |
| CC_DP3 | 132 | 132 | 0.000 | 3.867 | 25.980 | 246 | 246 | 0.000 | 3.874 | 18.888 |
| T2_D1_DP2 | 132 | 132 | 0.249 | 3.534 | 16.601 | 259 | 259 | 0.000 | 3.870 | 18.766 |
| CB_DP3 | 83 | 83 | 0.000 | 3.366 | 17.051 | 191 | 191 | 0.000 | 3.934 | 23.100 |
| T1_D2_DP2 | 37 | 37 | 0.000 | 2.530 | 6.914 | 37 | 37 | 0.000 | 0.813 | 1.385 |
| CA_DP1 | 27 | 27 | 0.491 | 2.754 | 11.335 | 39 | 39 | 0.000 | 2.658 | 8.080 |
| T4_D3_DP1 | 102 | 102 | 0.249 | 3.351 | 13.703 | 134 | 134 | 0.000 | 3.189 | 10.856 |
| T5_D2_DP3 | 127 | 127 | 0.125 | 3.946 | 31.173 | 241 | 241 | 0.000 | 3.841 | 19.957 |
| T3_D2_DP2 | 103 | 103 | 0.249 | 3.401 | 14.206 | 224 | 224 | 0.000 | 3.805 | 15.969 |
| T5_D2_DP2 | 137 | 137 | 0.000 | 3.247 | 9.615 | 232 | 232 | 0.000 | 4.041 | 23.481 |
| T5_D3_DP2 | 106 | 107 | 1.297 | 3.575 | 19.478 | 179 | 179 | 0.000 | 3.574 | 16.399 |
| T3_D3_DP1 | 68 | 68 | 0.000 | 3.179 | 13.360 | 76 | 76 | 0.000 | 3.039 | 11.153 |
| T4_D3_DP3 | 116 | 116 | 0.000 | 3.658 | 20.763 | 238 | 238 | 0.000 | 3.721 | 17.123 |
| T2_D1_DP3 | 95 | 95 | 0.000 | 3.528 | 19.945 | 201 | 201 | 0.000 | 3.607 | 14.357 |
| T4_D1_DP1 | 73 | 73 | 0.248 | 2.971 | 9.318 | 124 | 124 | 0.000 | 3.076 | 9.062 |
| T1_D2_DP3 | 5 | 5 | 0.000 | 1.141 | 2.447 | 8 | 8 | 0.000 | 1.320 | 2.821 |
| T4_D2_DP1 | 73 | 73 | 0.166 | 3.124 | 10.300 | 93 | 93 | 0.000 | 2.995 | 8.528 |
| CA_DP3 | 88 | 88 | 0.000 | 3.244 | 13.505 | 194 | 194 | 0.083 | 3.777 | 16.262 |
| T5_D2_DP1 | 62 | 62 | 0.248 | 3.082 | 10.824 | 91 | 91 | 0.000 | 2.804 | 8.248 |
| T5_D1_DP3 | 96 | 96 | 0.000 | 3.653 | 25.833 | 214 | 214 | 0.000 | 3.607 | 14.344 |
| CB_DP1 | 81 | 81 | 0.248 | 3.119 | 9.468 | 102 | 102 | 0.000 | 3.030 | 10.992 |
| T3_D2_DP3 | 129 | 129 | 0.000 | 3.755 | 24.337 | 267 | 267 | 0.000 | 3.767 | 17.659 |
| T5_D0_DP1 | 76 | 76 | 0.248 | 3.041 | 9.571 | 99 | 99 | 0.000 | 3.161 | 11.728 |
| T3_D3_DP3 | 138 | 138 | 0.000 | 3.405 | 12.819 | 277 | 277 | 0.125 | 3.939 | 20.533 |
| T3_D3_DP2 | 85 | 85 | 0.497 | 3.574 | 17.968 | 124 | 124 | 0.000 | 3.048 | 7.756 |
| T3_D1_DP2 | 109 | 109 | 0.000 | 3.271 | 11.243 | 232 | 232 | 0.125 | 4.026 | 22.827 |
| T1_D3_DP1 | 87 | 87 | 0.000 | 3.373 | 15.215 | 121 | 121 | 0.000 | 2.872 | 7.956 |
| CB_DP2 | 140 | 140 | 0.000 | 3.743 | 20.652 | 259 | 259 | 0.000 | 4.184 | 29.522 |
| T3_D2_DP1 | 84 | 84 | 0.000 | 3.165 | 10.993 | 115 | 115 | 0.000 | 3.242 | 12.697 |
| T5_D1_DP1 | 83 | 84 | 1.296 | 3.276 | 13.313 | 96 | 96 | 0.000 | 2.989 | 9.218 |
| T4_D3_DP2 | 133 | 133 | 0.000 | 3.468 | 14.709 | 268 | 268 | 0.000 | 3.976 | 19.510 |
| T3_D1_DP3 | 108 | 108 | 0.000 | 3.572 | 18.480 | 233 | 233 | 0.000 | 3.745 | 18.539 |
| T4_D2_DP3 | 104 | 104 | 0.249 | 3.765 | 27.606 | 269 | 269 | 0.000 | 3.833 | 17.585 |
| T3_D1_DP1 | 101 | 109 | 8.166 | 3.139 | 10.776 | 120 | 120 | 0.000 | 3.353 | 11.267 |
| T4_D2_DP2 | 104 | 104 | 0.000 | 3.501 | 15.716 | 212 | 212 | 0.100 | 3.780 | 14.313 |
| T5_D3_DP1 | 48 | 48 | 0.495 | 3.107 | 12.444 | 45 | 45 | 0.000 | 2.729 | 8.273 |
| T5_D1_DP2 | 157 | 157 | 0.000 | 3.788 | 21.886 | 283 | 283 | 0.000 | 3.957 | 22.703 |
| T1_D3_DP3 | 98 | 98 | 0.000 | 3.365 | 15.732 | 190 | 190 | 0.249 | 3.497 | 11.943 |
| T2_D1_DP1 | 2 | 2 | 0.000 | 0.629 | 1.777 | 2 | 2 | 0.000 | 0.673 | 1.923 |
| T2_D2_DP3 | 120 | 120 | 0.000 | 3.475 | 15.727 | 247 | 247 | 0.000 | 3.589 | 12.733 |
| T1_D1_DP2 | 84 | 85 | 1.296 | 3.501 | 19.304 | 118 | 118 | 0.000 | 2.522 | 4.175 |

Table S3 - Differentially abundant ASVs detected by the two differential abundance approaches (ALDEx2 and DESEq2).

| **Depth** | **Group** | **Phylum** | **Class** | **Order** | **Family** | **Genus** | **Species** | **n** | **Site** |
| --- | --- | --- | --- | --- | --- | --- | --- | --- | --- |
| bottom | Phytoplankton | SAR | Alveolata | Dinoflagellata | unknown | unknown | unknown | 1 | Control |
| bottom | Phytoplankton | SAR | Stramenopiles | Ochrophyta | Mediophyceae | Lauderia | Lauderia annulata | 1 | Lease 1 |
| surface | Phytoplankton | SAR | Alveolata | Dinoflagellata | Dinophyceae | uncultured | uncultured | 1 | Lease 1 |
| surface | Phytoplankton | SAR | Alveolata | Dinoflagellata | Gymnodiniphycidae | Gyrodinium | unknown | 1 | Lease 1 |
| bottom | Phytoplankton | SAR | Stramenopiles | Ochrophyta | Chrysophyceae | uncultured | uncultured | 3 | Lease 2 |
| bottom | Phytoplankton | SAR | Stramenopiles | Ochrophyta | unknown | unknown | unknown | 1 | Lease 2 |
| middle | Phytoplankton | Cryptophyceae | Cryptomonadales | uncultured | unknown | unknown | unknown | 1 | Lease 2 |
| surface | Phytoplankton | Archaeplastida | Chloroplastida | Chlorophyta | Dolichomastigales | Crustomastix | uncultured | 1 | Lease 2 |
| surface | Phytoplankton | SAR | Stramenopiles | Ochrophyta | Chrysophyceae | uncultured | unknown | 1 | Lease 2 |
| bottom | Other protists | SAR | Alveolata | Protalveolata | Syndiniales | Syndiniales_Group_I | uncultured | 1 | Control |
| bottom | Other protists | SAR | Alveolata | Protalveolata | Syndiniales | Syndiniales_Group_I | uncultured | 1 | Control |
| bottom | Other protists | SAR | Alveolata | Protalveolata | Syndiniales | Syndiniales_Group_II | uncultured | 1 | Control |
| middle | Other protists | SAR | Alveolata | Protalveolata | Syndiniales | Amoebophrya | uncultured | 1 | Control |
| middle | Other protists | SAR | Alveolata | Protalveolata | Syndiniales | uncultured | uncultured | 1 | Control |
| bottom | Other protists | SAR | Alveolata | Ciliophora | Scuticociliatia | Philasterides | Philasterides armatalis | 2 | Lease 1 |
| bottom | Other protists | SAR | Alveolata | Ciliophora | Plagiopylea | Plagiopylida | uncultured | 1 | Lease 1 |
| bottom | Other protists | SAR | Rhizaria | Cercozoa | Novel_Clade_2 | uncultured | uncultured | 1 | Lease 1 |
| surface | Other protists | SAR | Alveolata | Ciliophora | Oligotrichia | Strombidium | unknown | 1 | Lease 1 |
| surface | Other protists | SAR | Rhizaria | Cercozoa | Novel_Clade_2 | Uncultured | Uncultured | 1 | Lease 1 |
| surface | Other protists | SAR | Rhizaria | Cercozoa | uncultured | uncultured | uncultured | 1 | Lease 1 |
| bottom | Other protists | SAR | Alveolata | Protalveolata | Syndiniales | Amoebophrya | uncultured | 2 | Lease 2 |
| bottom | Other protists | SAR | Alveolata | Protalveolata | Syndiniales | Syndiniales_Group_II | uncultured | 2 | Lease 2 |
| bottom | Other protists | Opisthokonta | Holozoa | Choanoflagellida | Codonosigidae | Monosiga | uncultured | 1 | Lease 2 |
| bottom | Other protists | SAR | Alveolata | Ciliophora | Cyrtophoria | Pithites | unknown | 1 | Lease 2 |
| bottom | Other protists | SAR | Rhizaria | Cercozoa | Novel_Clade_2 | uncultured | uncultured | 1 | Lease 2 |
| bottom | Other protists | SAR | Rhizaria | Retaria | unknown | unknown | unknown | 1 | Lease 2 |
| bottom | Other protists | SAR | Stramenopiles | MAST-12 | MAST-12E | uncultured | uncultured | 1 | Lease 2 |
| surface | Other protists | SAR | Alveolata | Ciliophora | Choreotrichia | Tintinnidium | Tintinnopsis sp. TK-2011c | 1 | Lease 2 |
| surface | Other protists | SAR | Alveolata | Ciliophora | Haptoria | Hemiophrys | Amphileptus dragescoi | 1 | Lease 2 |
| surface | Other protists | SAR | Alveolata | Ciliophora | Haptoria | Hemiophrys | Hemiophrys macrostoma | 1 | Lease 2 |
| surface | Other protists | SAR | Alveolata | Ciliophora | Suctoria | Acineta | uncultured | 1 | Lease 2 |
| surface | Other protists | SAR | Rhizaria | Cercozoa | unknown | unknown | unknown | 1 | Lease 2 |

Table S4 - The 20 most important ASVs from both communities in the random forest classification models using “site” as class labels. Variable importance was based on Mean Decrease in Gini Index.

| ASV | Importance | Group | Phylum | Class | Order | Family | Genus | Species |
| --- | --- | --- | --- | --- | --- | --- | --- | --- |
| e_asv_663 | 8.75 | Phytoplankton | Cryptophyceae | Cryptomonadales | uncultured | uncultured | uncultured | uncultured |
| e_asv_1471 | 7.11 | Phytoplankton | SAR | Alveolata | Dinoflagellata | Gymnodinium_clade | Chytriodinium | uncultured |
| e_asv_751 | 6.87 | Phytoplankton | SAR | Stramenopiles | Ochrophyta | Dictyochophyceae | Pedinellales | Pedinellales sp. RCC2289 |
| e_asv_78 | 5.78 | Phytoplankton | SAR | Alveolata | Dinoflagellata | Gymnodiniphycidae | Gyrodinium | unknown |
| e_asv_690 | 5.66 | Phytoplankton | Cryptophyceae | Cryptomonadales | Cryptomonas | unknown | unknown | unknown |
| e_asv_1778 | 5.39 | Phytoplankton | SAR | Stramenopiles | Ochrophyta | MOCH-2 | uncultured | uncultured |
| e_asv_714 | 5.22 | Phytoplankton | SAR | Alveolata | Dinoflagellata | unknown | unknown | unknown |
| e_asv_1108 | 5.10 | Phytoplankton | SAR | Stramenopiles | Ochrophyta | Coscinodiscophytina | Actinocyclus | Actinocyclus sp. 1 MPA-2013 |
| e_asv_1884 | 5.06 | Phytoplankton | Archaeplastida | Chloroplastida | Chlorophyta | Mamiellales | Bathycoccus | unknown |
| e_asv_1920 | 4.80 | Phytoplankton | Haptophyta | Prymnesiophyceae | Prymnesiales | OLI51059 | uncultured | uncultured |
| e_asv_320 | 4.80 | Phytoplankton | SAR | Stramenopiles | Ochrophyta | Bolidomonas | uncultured | uncultured |
| e_asv_790 | 4.64 | Phytoplankton | Archaeplastida | Chloroplastida | Chlorophyta | Mamiellales | Ostreococcus | Ostreococcus sp. RCC356 |
| e_asv_1919 | 4.56 | Phytoplankton | SAR | Stramenopiles | Ochrophyta | Bolidomonas | uncultured | uncultured |
| e_asv_437 | 4.30 | Phytoplankton | SAR | Stramenopiles | Ochrophyta | Ochromonadales | Paraphysomonas | unknown |
| e_asv_436 | 4.19 | Phytoplankton | SAR | Alveolata | Dinoflagellata | unknown | unknown | unknown |
| e_asv_175 | 4.13 | Phytoplankton | SAR | Stramenopiles | Ochrophyta | Chrysophyceae | uncultured | uncultured |
| e_asv_1380 | 4.08 | Phytoplankton | Archaeplastida | Chloroplastida | Chlorophyta | Pyramimonadales | Pterosperma | Pterosperma cristatum |
| e_asv_551 | 4.06 | Phytoplankton | SAR | Stramenopiles | Ochrophyta | Bolidomonas | uncultured | uncultured |
| e_asv_268 | 4.01 | Phytoplankton | Cryptophyceae | Cryptomonadales | uncultured | unknown | unknown | unknown |
| e_asv_579 | 3.66 | Phytoplankton | SAR | Alveolata | Dinoflagellata | unknown | unknown | unknown |
| e_asv_308 | 4.83 | Other protists | SAR | Rhizaria | Cercozoa | Novel_Clade_2 | uncultured | uncultured |
| e_asv_931 | 4.83 | Other protists | SAR | Alveolata | Protalveolata | Syndiniales | Amoebophrya | uncultured_eukaryote |
| e_asv_1394 | 4.71 | Other protists | SAR | Alveolata | Protalveolata | Syndiniales | Amoebophrya | uncultured_eukaryote |
| e_asv_1654 | 4.55 | Other protists | SAR | Alveolata | unknown | unknown | unknown | unknown |
| e_asv_738 | 4.43 | Other protists | SAR | Alveolata | Protalveolata | Syndiniales | Syndiniales_Group_II | uncultured_eukaryote |
| e_asv_669 | 4.29 | Other protists | Centrohelida | unknown | unknown | unknown | unknown | unknown |
| e_asv_2541 | 4.28 | Other protists | Amoebozoa | Tubulinea | Euamoebida | unknown | unknown | unknown |
| e_asv_1436 | 3.70 | Other protists | SAR | Stramenopiles | Labyrinthulomycetes | Thraustochytriaceae | uncultured | uncultured |
| e_asv_3555 | 3.56 | Other protists | SAR | Stramenopiles | Incertae_Sedis | Pirsonia | unknown | unknown |
| e_asv_760 | 3.56 | Other protists | SAR | Alveolata | unknown | unknown | unknown | unknown |
| e_asv_1739 | 3.40 | Other protists | Incertae_Sedis | Rigifilida | Micronuclearia | Micronuclearia podoventralis | Micronuclearia podoventralis | Micronuclearia podoventralis |
| e_asv_2664 | 3.37 | Other protists | SAR | Stramenopiles | MAST-3 | MAST-3A | uncultured | uncultured |
| e_asv_12 | 3.34 | Other protists | SAR | Alveolata | Ciliophora | Prostomatea | Cryptocaryon | Askenasia sp. LWW2010032604 |
| e_asv_1006 | 3.29 | Other protists | SAR | Rhizaria | Cercozoa | Novel_Clade_2 | uncultured | uncultured |
| e_asv_741 | 3.22 | Other protists | SAR | unknown | unknown | unknown | unknown | unknown |
| e_asv_780 | 3.21 | Other protists | Excavata | Discoba | Discicristata | unknown | unknown | unknown |
| e_asv_81 | 3.15 | Other protists | SAR | Alveolata | Ciliophora | Oligotrichia | Strombidium | unknown |
| e_asv_426 | 3.14 | Other protists | SAR | unknown | unknown | unknown | unknown | unknown |
| e_asv_446 | 3.11 | Other protists | SAR | Alveolata | Ciliophora | Hypotrichia | Trachelostyla | Trachelostyla_pediculiformis |
| e_asv_851 | 3.09 | Other protists | SAR | Stramenopiles | Labyrinthulomycetes | Thraustochytriaceae | uncultured | uncultured |

Table S5 - The 20 most important ASVs from both communities in the random forest classification models using “depth” as class labels. . Variable importance was based on Mean Decrease in Gini Index.

| ASV | Importance | Group | Phylum | Class | Order | Family | Genus | Species |
| --- | --- | --- | --- | --- | --- | --- | --- | --- |
| e_asv_26 | 5.80 | Phytoplankton | SAR | Alveolata | Dinoflagellata | Thoracosphaeraceae | Scrippsiella | Peridiniopsis polonicum |
| e_asv_9 | 5.75 | Phytoplankton | SAR | Alveolata | Dinoflagellata | Thoracosphaeraceae | Pfiesteria | uncultured |
| e_asv_31 | 5.33 | Phytoplankton | SAR | Alveolata | Dinoflagellata | Dinophyceae | uncultured | unknown |
| e_asv_84 | 5.25 | Phytoplankton | SAR | Alveolata | Dinoflagellata | Peridiniales | Heterocapsa | uncultured |
| e_asv_14 | 5.02 | Phytoplankton | SAR | Alveolata | Dinoflagellata | Gymnodinium_clade | Chytriodinium | uncultured |
| e_asv_221 | 4.98 | Phytoplankton | SAR | Alveolata | Dinoflagellata | Gymnodiniphycidae | Amphidinium | Amphidinium longum |
| e_asv_33 | 4.96 | Phytoplankton | SAR | Alveolata | Dinoflagellata | unknown | unknown | unknown |
| e_asv_87 | 4.88 | Phytoplankton | SAR | Alveolata | Dinoflagellata | Gymnodinium_clade | Chytriodinium | uncultured |
| e_asv_156 | 4.87 | Phytoplankton | SAR | Alveolata | Dinoflagellata | unknown | unknown | unknown |
| e_asv_289 | 4.76 | Phytoplankton | SAR | Stramenopiles | Ochrophyta | Chrysophyceae | uncultured | uncultured |
| e_asv_201 | 4.73 | Phytoplankton | SAR | Stramenopiles | Ochrophyta | Chrysophyceae | uncultured | uncultured |
| e_asv_15 | 4.70 | Phytoplankton | SAR | Alveolata | Dinoflagellata | unknown | unknown | unknown |
| e_asv_194 | 4.66 | Phytoplankton | SAR | Stramenopiles | Ochrophyta | Chrysophyceae | uncultured | uncultured |
| e_asv_198 | 4.40 | Phytoplankton | SAR | Stramenopiles | Ochrophyta | Ochromonadales | Dinobryon | uncultured |
| e_asv_666 | 4.25 | Phytoplankton | Archaeplastida | Chloroplastida | Chlorophyta | Dolichomastigales | Crustomastix | unknown |
| e_asv_285 | 4.03 | Phytoplankton | SAR | Alveolata | Dinoflagellata | Gymnodiniphycidae | uncultured | uncultured |
| e_asv_210 | 3.85 | Phytoplankton | SAR | Stramenopiles | Ochrophyta | unknown | unknown | unknown |
| e_asv_371 | 3.58 | Phytoplankton | SAR | Stramenopiles | Ochrophyta | Mediophyceae | Chaetoceros | Chaetoceros sp. UNC1415 |
| e_asv_38 | 3.52 | Phytoplankton | SAR | Alveolata | Dinoflagellata | Gymnodinium_clade | unknown | unknown |
| e_asv_83 | 3.45 | Phytoplankton | SAR | Stramenopiles | Ochrophyta | Chrysophyceae | uncultured | uncultured |
| e_asv_22 | 7.11 | Other protists | SAR | Alveolata | Protalveolata | unknown | unknown | unknown |
| e_asv_191 | 5.37 | Other protists | SAR | Alveolata | Ciliophora | Choreotrichia | uncultured | uncultured |
| e_asv_118 | 5.13 | Other protists | SAR | Stramenopiles | Bicosoecida | Pseudobodo | uncultured | uncultured |
| e_asv_94 | 4.83 | Other protists | SAR | Alveolata | Protalveolata | Syndiniales | Amoebophrya | uncultured |
| e_asv_32 | 4.68 | Other protists | SAR | Alveolata | Ciliophora | Oligotrichia | Strombidium | uncultured |
| e_asv_280 | 4.60 | Other protists | SAR | Alveolata | Protalveolata | Syndiniales | Syndiniales_Group_II | uncultured |
| e_asv_318 | 4.47 | Other protists | SAR | Stramenopiles | MAST-3 | MAST-3J | unknown | unknown |
| e_asv_34 | 4.46 | Other protists | Cryptophyceae | Kathablepharidae | Leucocryptos | uncultured | uncultured | uncultured |
| e_asv_213 | 4.25 | Other protists | SAR | Alveolata | Protalveolata | Syndiniales | Syndiniales_Group_II | uncultured |
| e_asv_682 | 3.98 | Other protists | SAR | Alveolata | Protalveolata | Syndiniales | Syndiniales_Group_II | unknown |
| e_asv_164 | 3.98 | Other protists | SAR | Alveolata | uncultured | unknown | unknown | unknown |
| e_asv_299 | 3.91 | Other protists | SAR | Alveolata | Protalveolata | Syndiniales | Amoebophrya | uncultured |
| e_asv_183 | 3.81 | Other protists | Incertae_Sedis | Telonema | uncultured | uncultured | uncultured | uncultured |
| e_asv_2 | 3.67 | Other protists | SAR | Alveolata | Protalveolata | Syndiniales | Syndiniales_Group_I | uncultured |
| e_asv_233 | 3.66 | Other protists | SAR | Alveolata | Ciliophora | Prostomatea | Cryptocaryon | uncultured |
| e_asv_230 | 3.59 | Other protists | SAR | Alveolata | unknown | unknown | unknown | unknown |
| e_asv_113 | 3.56 | Other protists | SAR | Alveolata | Protalveolata | Syndiniales | Syndiniales_Group_II | uncultured |
| e_asv_330 | 3.31 | Other protists | SAR | Stramenopiles | Bicosoecida | Pseudobodo | Pseudobodo_tremulans | Pseudobodo tremulans |
| e_asv_202 | 3.22 | Other protists | SAR | Alveolata | Protalveolata | Syndiniales | Syndiniales_Group_II | uncultured |
| e_asv_211 | 3.21 | Other protists | SAR | Rhizaria | Cercozoa | Novel_Clade_2 | uncultured | uncultured |

Table S6 - The common ASVs detected in at least two out of three methods used in this study (ALDEX2, DESEq2 and random forest classification model - RF) using “site” as class labels. Relative abundance (%) and depth are displayed at the last four right columns.

| **^ASV^** | **^name^** | **^Group^** | **^Phylum^** | **^Class^** | **^Order^** | **^Family^** | **^Genus^** | **^Species^** | **^Control^** | **^lease_1^** | **^lease_2^** | **^Depth^** |
| --- | --- | --- | --- | --- | --- | --- | --- | --- | --- | --- | --- | --- |
| ^e_asv_127^ | ^ALDEx2..DESeq2^ | ^Phytoplankton^ | ^SAR^ | ^Stramenopiles^ | ^Ochrophyta^ | ^Phaeophyceae^ | ^Ectocarpales^ | ^Ectocarpus sp.^ | ^0.0%^ | ^44.5%^ | ^55.5%^ | ^surface^ |
| ^e_asv_147^ | ^ALDEx2..DESeq2^ | ^Phytoplankton^ | ^SAR^ | ^Stramenopiles^ | ^Ochrophyta^ | ^Chrysophyceae^ | ^uncultured^ | ^uncultured^ | ^12.3%^ | ^0.5%^ | ^87.2%^ | ^bottom^ |
| ^e_asv_181^ | ^ALDEx2..DESeq2^ | ^Phytoplankton^ | ^SAR^ | ^Stramenopiles^ | ^Ochrophyta^ | ^Chrysophyceae^ | ^uncultured^ | ^uncultured^ | ^40.3%^ | ^1.7%^ | ^58.0%^ | ^bottom^ |
| ^e_asv_97^ | ^RF..ALDEx2^ | ^Phytoplankton^ | ^SAR^ | ^Stramenopiles^ | ^Ochrophyta^ | ^Chrysophyceae^ | ^uncultured^ | ^Spumella sp.^ | ^20.8%^ | ^52.9%^ | ^26.3%^ | ^middle^ |
| ^e_asv_175^ | ^RF..ALDEx2..DESeq2^ | ^Phytoplankton^ | ^SAR^ | ^Stramenopiles^ | ^Ochrophyta^ | ^Chrysophyceae^ | ^uncultured^ | ^uncultured^ | ^2.5%^ | ^7.9%^ | ^89.6%^ | ^surface^ |
| ^e_asv_78^ | ^RF..DESeq2^ | ^Phytoplankton^ | ^SAR^ | ^Alveolata^ | ^Dinoflagellata^ | ^Gymnodiniphycidae^ | ^Gyrodinium^ | ^uncultured^ | ^0.0%^ | ^96.4%^ | ^3.6%^ | ^surface^ |
| ^e_asv_268^ | ^RF..DESeq2^ | ^Phytoplankton^ | ^Cryptophyceae^ | ^Cryptomonadales^ | ^uncultured^ | ^uncultured^ | ^uncultured^ | ^uncultured^ | ^0.0%^ | ^27.2%^ | ^72.8%^ | ^middle^ |
| ^e_asv_203^ | ^ALDEx2..DESeq2^ | ^Other protists^ | ^SAR^ | ^Alveolata^ | ^Protalveolata^ | ^Syndiniales^ | ^Syndiniales_Group_I^ | ^uncultured^ | ^0.0%^ | ^53.5%^ | ^46.5%^ | ^middle^ |
| ^e_asv_184^ | ^ALDEx2..DESeq2^ | ^Other protists^ | ^SAR^ | ^Alveolata^ | ^Protalveolata^ | ^Syndiniales^ | ^Syndiniales_Group_I^ | ^uncultured^ | ^89.7%^ | ^0.0%^ | ^10.3%^ | ^bottom^ |
| ^e_asv_61^ | ^ALDEx2..DESeq2^ | ^Other protists^ | ^SAR^ | ^Alveolata^ | ^Protalveolata^ | ^Syndiniales^ | ^Syndiniales_Group_II^ | ^uncultured^ | ^49.8%^ | ^4.2%^ | ^46.0%^ | ^bottom^ |
| ^e_asv_94^ | ^ALDEx2..DESeq2^ | ^Other protists^ | ^SAR^ | ^Alveolata^ | ^Protalveolata^ | ^Syndiniales^ | ^Amoebophrya^ | ^uncultured^ | ^45.6%^ | ^5.1%^ | ^49.4%^ | ^bottom^ |
| ^e_asv_114^ | ^ALDEx2..DESeq2^ | ^Other protists^ | ^SAR^ | ^Rhizaria^ | ^Cercozoa^ | ^Novel_Clade_2^ | ^uncultured^ | ^uncultured^ | ^40.6%^ | ^4.7%^ | ^54.7%^ | ^bottom^ |
| ^e_asv_121^ | ^ALDEx2..DESeq2^ | ^Other protists^ | ^SAR^ | ^Alveolata^ | ^Protalveolata^ | ^Syndiniales^ | ^Amoebophrya^ | ^uncultured^ | ^34.9%^ | ^2.0%^ | ^63.0%^ | ^bottom^ |
| ^e_asv_309^ | ^ALDEx2..DESeq2^ | ^Other protists^ | ^SAR^ | ^Alveolata^ | ^Ciliophora^ | ^Scuticociliatia^ | ^Philasterides^ | ^Philasterides armatalis^ | ^0.0%^ | ^71.2%^ | ^28.8%^ | ^bottom^ |
| ^e_asv_108^ | ^ALDEx2..DESeq2^ | ^Other protists^ | ^SAR^ | ^Alveolata^ | ^Ciliophora^ | ^Plagiopylea^ | ^Plagiopylida^ | ^uncultured^ | ^0.0%^ | ^99.7%^ | ^0.3%^ | ^bottom^ |
| ^e_asv_1006^ | ^RF..ALDEx2^ | ^Other protists^ | ^SAR^ | ^Rhizaria^ | ^Cercozoa^ | ^Novel_Clade_2^ | ^uncultured^ | ^uncultured^ | ^0.0%^ | ^100.0%^ | ^0.0%^ | ^bottom^ |
| ^e_asv_81^ | ^RF..ALDEx2..DESeq2^ | ^Other protists^ | ^SAR^ | ^Alveolata^ | ^Ciliophora^ | ^Oligotrichia^ | ^Strombidium^ | ^uncultured^ | ^0.0%^ | ^82.0%^ | ^18.0%^ | ^surface^ |
| ^e_asv_144^ | ^RF..DESeq2^ | ^Other protists^ | ^SAR^ | ^Alveolata^ | ^Protalveolata^ | ^Syndiniales^ | ^Amoebophrya^ | ^uncultured^ | ^32.8%^ | ^5.6%^ | ^61.6%^ | ^bottom^ |
| ^e_asv_379^ | ^RF..DESeq2^ | ^Other protists^ | ^SAR^ | ^Alveolata^ | ^Protalveolata^ | ^Syndiniales^ | ^Syndiniales_Group_II^ | ^uncultured^ | ^47.5%^ | ^0.0%^ | ^52.5%^ | ^bottom^ |
